# Porcine intestinal innate lymphoid cells and lymphocyte spatial context revealed through single-cell RNA sequencing

**DOI:** 10.1101/2022.01.09.475571

**Authors:** Jayne E. Wiarda, Julian M. Trachsel, Sathesh K. Sivasankaran, Christopher K. Tuggle, Crystal L. Loving

## Abstract

Intestinal lymphocytes are crucial members of the mucosal immune system with impact over outcomes of intestinal health versus dysbiosis. Resolving intestinal lymphocyte complexity and function is a challenge, as the intestine provides cellular snapshots of a diverse spectrum of immune states. In pigs, intestinal lymphocytes are poorly described relative to humans or traditional model species. Enhanced understanding of porcine intestinal lymphocytes will promote food security and improve utility of pigs as a biomedical model for intestinal research. Single-cell RNA sequencing (scRNA-seq) was performed to provide transcriptomic profiles of lymphocytes in porcine ileum, with 31,983 cells annotated into 26 cell types. Deeper interrogation revealed previously undescribed cells in porcine intestine, including *SELL*^hi^ *γδ* T cells, group 1 and group 3 innate lymphoid cells (ILCs), and four subsets of B cells. Single-cell transcriptomes in ileum were compared to those in porcine blood, and subsets of activated lymphocytes were detected in ileum but not periphery. Comparison to scRNA-seq human and murine ileum data revealed a general consensus of ileal lymphocytes across species. Lymphocyte spatial context in porcine ileum was conferred through differential tissue dissection prior to scRNA-seq. Antibody-secreting cells, B cells, follicular CD4 *αβ* T cells, and cycling T/ILCs were enriched in ileum with Peyer’s patches, while non-cycling *γδ* T, CD8 *αβ* T, and group 1 ILCs were enriched in ileum without Peyer’s patches. scRNA-seq findings were leveraged to develop advanced toolsets for further identification of ILCs in porcine ileum via flow cytometry and *in situ* staining. Porcine ileal ILCs identified via scRNA-seq did not transcriptionally mirror peripheral porcine ILCs (corresponding to natural killer cells) but instead had gene signatures indicative of tissue- and activation-specific functions, indicating potentially similar roles to intestinal ILCs identified in humans. Overall, the data serve as a highly-resolved transcriptomic atlas of the porcine intestinal immune landscape and will be useful in further understanding intestinal immune cell function.

## INTRODUCTION

The intestine is a selectively permeable barrier that absorbs nutrients while simultaneously limiting entry of potentially harmful external organisms and compounds. Thus, the intestinal immune system continuously deciphers between innocuous and dangerous stimuli. Coordination of immune responses is crucial for maintaining intestinal homeostasis; dysregulation of even a small number of cells can negatively impact intestinal health, as evidenced in non-pathogenic inflammatory conditions such as celiac disease, Crohn’s disease, and ulcerative colitis (reviewed by Caio et al., 2019; Caminero and Pinto-Sanchez, 2020; Caruso et al., 2020; Mowat and Agace, 2014). Intestinal lymphocytes include B cells, T cells, and innate lymphoid cells (ILCs). The importance of lymphocytes in promoting intestinal homeostasis is well-documented in cases of intestinal dysbiosis in individuals naturally lacking at least some lymphocyte populations (reviewed by Agarwal and Cunningham-Rundles, 2019) or experimental models where lymphocytes are depleted or immune pathways disrupted (Gärdby and Lycke, 2000; Hepworth et al., 2015; Kühn et al., 1993; Laroux et al., 2004; Mombaerts et al., 1993; Sadlack et al., 1993; Strober and Ehrhardt, 1993; Wang et al., 2017). Lymphocytes can be directed to provide protective adaptive immunity through mucosal vaccination strategies (reviewed by Lavelle and Ward, 2021; Li et al., 2020), while immune protection against a broad range of microorganisms may be achieved through non-conventional innate memory in some lymphocytes (reviewed by Wang et al., 2019b). Resultingly, there is pan-disciplinary interest in promoting health through modulation of intestinal lymphocytes, but decoding the complexity and function of these cells is an ongoing challenge. The intestine is a site of vast immune cellular diversity difficult to holistically characterize, yet defining heterogeneity within the cellular landscape of intestinal immune cells, such as lymphocytes, is one initial step to be taken towards better understanding intestinal immune dynamics and resulting effects on health.

Pigs (*Sus scrofa*) are a promising biomedical model and major global food source, yet the porcine intestinal immune cell landscape is poorly defined relative to humans and rodent models. Deeper exploration of the porcine intestinal immune system, particularly intestinal lymphocytes, will enhance utility of pigs as a well-defined and highly comparable biomedical model for gut health and/or disease (reviewed by Gonzalez et al., 2015; Käser, 2021; Roura et al., 2016; Ziegler et al., 2016), as pigs have greater physiologic and genetic similarities to humans than rodent models and are less expensive and more easily obtained than non-human primates (reviewed by Gün and Kues, 2014; Kobayashi et al., 2018; Swindle et al., 2011). Enhanced characterization of porcine intestinal lymphocytes will also provide insight into promoting gut health and associated overall pig health to ultimately decrease disease susceptibility and strengthen pork as a major global food source. Though previous work has described porcine lymphocytes at the protein level (reviewed by Piriou-Guzylack and Salmon, 2008), annotations are confined by a limited toolbox of available porcine protein-specific immunoreagents (reviewed by Entrican et al., 2020). Thus, definitions of porcine lymphocytes lack cellular resolution comparable to that of humans. This is particularly true for B cells, as a pan-B cell-specific extracellular protein marker is not available (reviewed by Piriou-Guzylack and Salmon, 2008; Sinkora and Butler, 2009), and ILCs, for which only natural killer (NK) cells have been identified (reviewed by Gerner et al., 2009). Approaches to resolve the porcine immune cell landscape at the transcriptional level have also been employed; however, traditional bulk RNA sequencing (RNA-seq) or microarray approaches fail to provide cellular resolution needed to decode such a complex cellular community (Herrera-Uribe & Wiarda et al., 2021), especially when immunoreagents for sorting of cells into more homogenous populations are lacking.

Numerous studies have assessed transcriptional dynamics in the porcine intestinal tract but did not attempt to deconvolute cells into specific populations, a critical step in understanding functions of specific cells (Beiki et al., 2019; Freeman et al., 2012; Inoue et al., 2015; Jin et al., 2021; Mach et al., 2014; Maroilley et al., 2018; Meng et al., 2020; Pan et al., 2021; Summers et al., 2020; Tan et al., 2017; Wang et al., 2008; Wang et al., 2019a; Zhu et al., 2014). Some bulk RNA-seq studies have sorted porcine immune cells into specific populations based on cell surface markers but primarily focused on studying cells from the periphery and non-intestinal tissues (Auray et al., 2016; Auray et al., 2020; Foissac et al., 2019; Herrera-Uribe & Wiarda et al., 2021; Kim et al., 2021). Consequently, it remains to be determined whether existing data adequately portray the transcriptional heterogeneity of intestinal immune cells or if novelties exist in the context of the porcine intestine.

Single-cell RNA-seq (scRNA-seq) has been utilized to describe porcine immune cell transcriptomes at granularity unparalleled by bulk RNA-seq or microarray approaches, including in peripheral blood (Herrera-Uribe & Wiarda et al., 2021), lung (Zhang et al., 2021b), skin (Han et al., 2021), brain (Chen et al., 2019; Zhu et al., 2021), and embryos (Kong et al., 2020; Liu et al., 2021; Ramos-Ibeas et al., 2019). Additionally, epithelial cells in the porcine intestine were recently queried via scRNA-seq, and results provide new insight into biological development and epithelial cell functions (Meng et al., 2021). However, high-resolution transcriptomic analysis of porcine intestinal immune cells remains to be completed. We therefore utilized scRNA-seq to provide the first high-resolution, global transcriptomic profiles of porcine intestinal lymphocytes. Interrogation was focused to the ileum, the most distal segment of the small intestine, which contains a unique combination of not only lymphocytes residing in the lamina propria and epithelium but also lymphocytes found in association with gut-associated lymphoid tissue (GALT) called Peyer’s patches. Peyer’s patches are major sites of immune induction not highly prevalent in other intestinal segments (Bonnardel et al., 2015; Fujihashi et al., 2001; Keren et al., 1978; Kiriya et al., 2007; Kwa et al., 2006; Mora et al., 2003; Nagai et al., 2007). In pigs, ileal Peyer’s patches present as a continuous longitudinal strip along the length of the distal small intestine (Binns and Licence, 1985; Rothkötter, 2009) and are more easily identified and obtained compared to Peyer’s patches in humans and rodents, the species in which scRNA-seq approaches have been mostly employed. Consequently, pigs are an ideal candidate for studying Peyer’s patches due to easier gross identification and isolation for further study.

By performing scRNA-seq on porcine ileal-derived cells, we documented and showcased previously undescribed levels of cellular heterogeneity for multiple populations of lymphocytes, as well as some non-lymphocytes. Profiling of porcine ileal cells was completed with multiple approaches, including cross-location and cross-species analyses. Data were compared to an annotated reference scRNA-seq dataset of porcine peripheral blood mononuclear cells (PBMCs; Herrera-Uribe & Wiarda et al., 2021) to reveal transcriptional differences between porcine intestinal-derived cells and circulating counterparts. Comparison to human and murine ileum reference datasets (Elmentaite & Ross et al., 2020; Xu et al., 2019) unveiled similarities and differences for cells of the same intestinal location across species. We further recognized cells associated specifically with Peyer’s patches or the epithelium/lamina propria and confirmed findings by *in situ* and *ex vivo* detection using available canonical cell markers with locational context to further infer potential cell functions. Previously undescribed lymphocyte populations in pigs, particularly intestinal ILCs, were identified and characterized. We further leveraged our single-cell gene expression profiles to develop new cell marker combinations with currently available immunoreagents to label novel populations. ILC locational context within the ileum was determined, and transcriptional distinctions from circulating NK cells was denoted. Collectively, the data serve as the highest-resolved transcriptomic atlas of the porcine intestinal immune landscape and may be used to further decode cellular phenotype and function within the intestinal tract.

## RESULTS

### Experimental overview

From each of two pigs, distal ileum was grossly dissected into three distinct sections for cell isolation: (1) ileal tissue containing only regions with Peyer’s patches (PP), (2) ileal tissue excluding regions with Peyer’s patches (non-PP), and (3) a complete cross section of ileal tissue containing both regions (whole ileum; **Figure 1A**; **Supplement to Figure 1-Figure 1A-B**). For each region, a single-cell suspension of combined epithelium, lamina propria, Peyer’s patches (if present), and submucosa was retrieved, enriched for viable lymphocytes, and submitted for scRNA-seq, as described in materials and methods. Sequencing and further processing/quality control of scRNA-seq data are fully described in materials and methods and are shown in **Supplement to Figure 1-Figure 2A-D** and **Supplementary Data 1**. Our final dataset contained 31,983 total cells from six ileal samples (**Figure 1B**). Cells were classified into four cell lineages and further annotated as 26 cell types (**Figure 1C-D**; **Supplement to Figure 1-Figure 3**) using a multi-method annotation approach described fully in materials and methods sections, ‘*Ileal cell lineage annotations’* and *‘Ileal cell type annotations’*, and shown in **Supplement to Figure 1 - Figures 4-10** and **Supplementary Data 2-8.**

**Figure 1.**
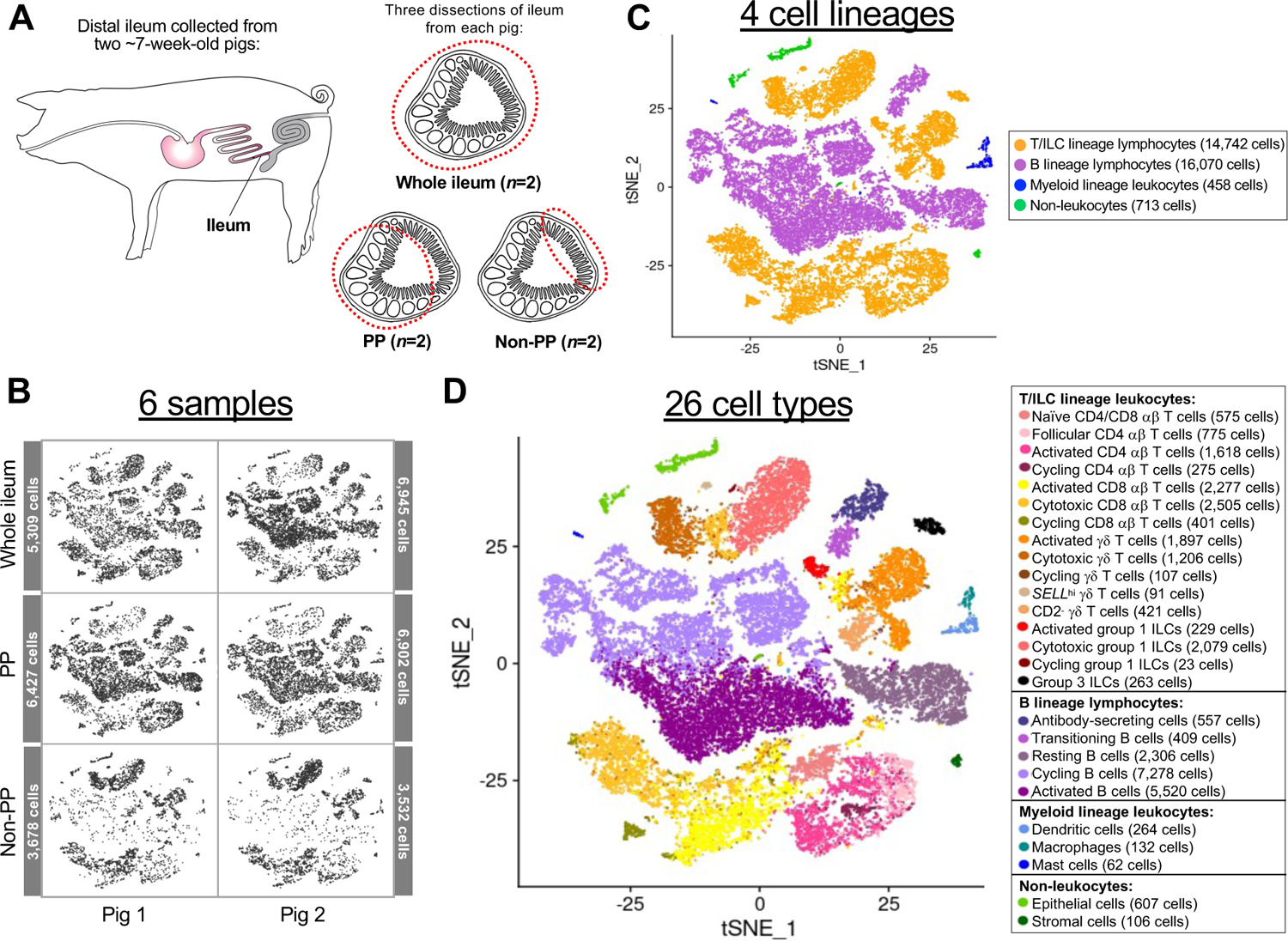
Experimental overview and annotation of cells recovered from scRNA-seq of porcine ileum. **(A)** Ileal samples collected from two seven-week-old pigs for scRNA-seq. Left: representative image of tissue collection site from ileum of the distal small intestine within the porcine gastrointestinal tract. Right: representative images of tissue dissections from transverse cross sections of ileum. Dissections from each pig included a cross section of whole ileum including areas with and without Peyer’s patches (whole ileum), ileum containing only regions with Peyer’s patches (PP), and ileum containing only regions without Peyer’s patches (non-PP), resulting in a total of six samples processed for scRNA-seq. **(B-D)** Two-dimensional t-SNE visualization of 31,983 cells isolated from porcine ileal samples described in **A**, subjected to scRNA-seq, and included in the final dataset following data processing and quality filtering. Each point represents a single cell. Plots show which sample cells are derived from **(B)** and cell lineage **(C)** or cell type **(D)** annotations. In **B**, cells in individual panels are derived from a specified sample. In **C-D**, the color of a cell indicates cell lineage **(C)** or cell type **(D)** annotation. The number of cells belonging to each sample (**B**), cell lineage (**C**), and cell type (**D**) are listed next to corresponding panels. Abbreviations: ILC (innate lymphoid cell); PP (Peyer’s patch); scRNA-seq (single-cell RNA sequencing); t-SNE (t-distributed stochastic neighbor embedding)

**Figure 2.**
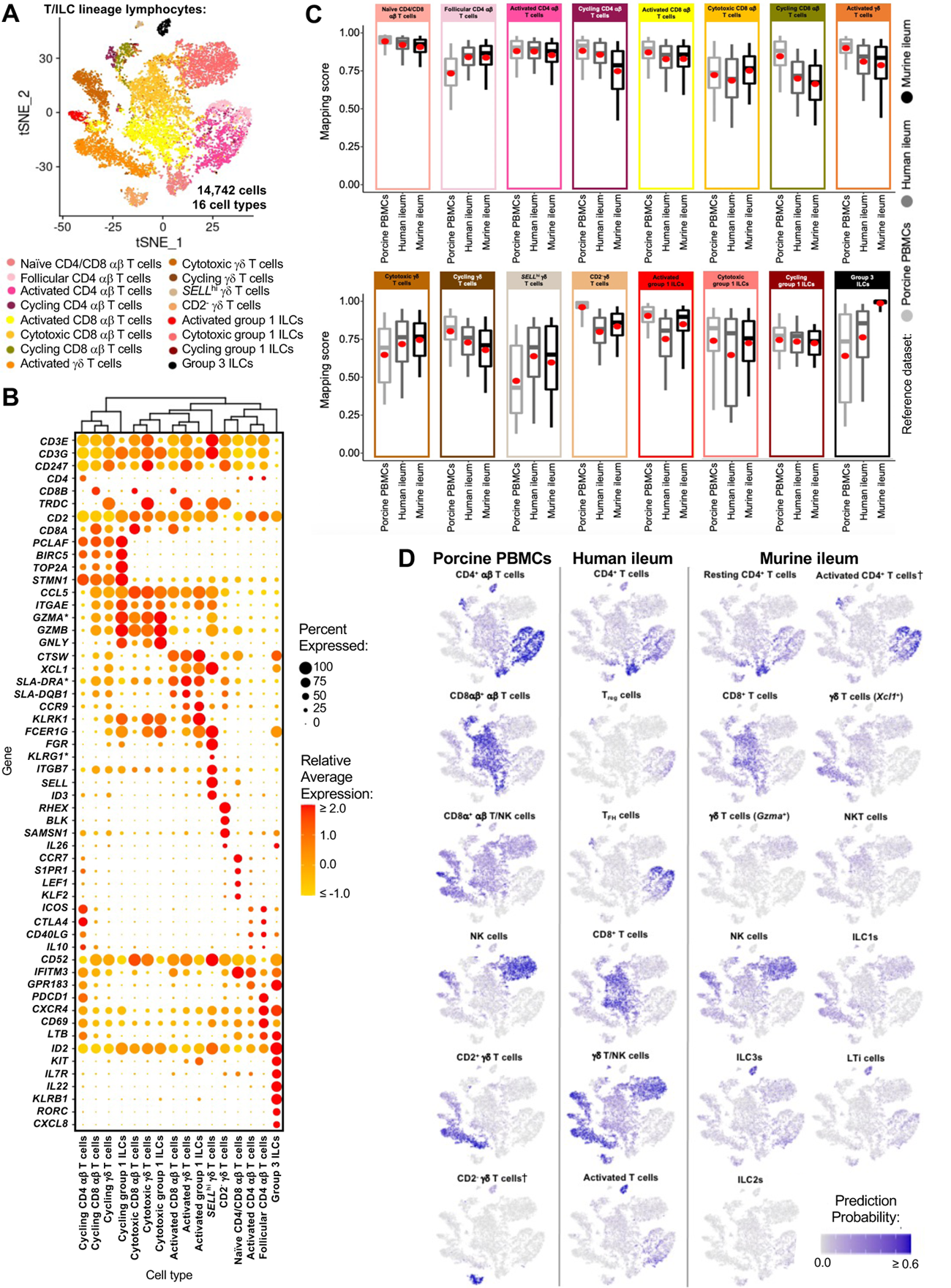
scRNA-seq profiles of T/ILC lineage lymphocytes in porcine ileum. **(A)** Two-dimensional t-SNE visualization of 14,742 cells recovered from porcine ileum via scRNA-seq that were classified as T/ILC lineage lymphocytes in Figure 1C & **Supplement to Figure 1-Figure 4B** and further annotated into 16 cell types in Figure 1D & **Supplement to Figure 1-Figures 5-8**. Each point represents a single cell; the color of each point indicates cell types shown in Figure 1D. **(B)** Hierarchical relationship of T/ILC lineage lymphocyte cell types from porcine ileum shown in a dendrogram (upper) and expression patterns of selected genes within each cell type shown in a dot plot (lower). In the dot plot, selected genes are listed on the y-axis, and cell types are listed on the x-axis. Within the dot plot, size of a dot corresponds to the percentage of cells expressing a gene within an annotated cell type; color of a dot corresponds to average expression level of a gene for those cells expressing it within a cell type relative to all other cells in the dataset shown in **A**. **(C)** Box plots of the distribution of mapping scores for T/ILC lineage lymphocyte cell types from porcine ileum mapped to each reference scRNA-seq dataset. Results for a single cell type are located within a single box, with color of the box corresponding to colors used for cell types in **A**. The color of each box in a plot corresponds to the reference dataset porcine ileal cells were mapped to, including porcine PBMCs (light grey), human ileum (medium grey), and murine ileum (black). Boxes span the interquartile range (IQR) of the data (25^th^ and 75^th^ percentiles), with the median (50^th^ percentile) indicated by a horizontal line. Whiskers span the 5^th^ and 95^th^ percentiles of the data. A red dot represents the data mean. **(D)** Prediction probabilities for porcine ileum scRNA-seq query data from label transfer of selected annotated T/ILC types in reference scRNA-seq datasets of porcine PBMCs (left), human ileum (middle), and murine ileum (right) overlaid onto two-dimensional t-SNE visualization shown in **A**. Each point represents a single cell; the color of each point indicates prediction probability to a corresponding cell type from reference data, as indicated directly above each t-SNE plot. A higher prediction probability indicates higher similarity to a specified annotated cell type in a reference scRNA-seq dataset. scRNA-seq data shown in **A-D** were derived from ileum of two seven-week-old pigs. *Ensembl identifiers found in gene annotation were converted to gene symbols; refer to methods section ‘*Gene name modifications*’ for more details † Identical cell type annotations were given to cells in both porcine ileum and a reference scRNA-seq dataset. Cell type annotations were given to each dataset by independent rationales, and identical annotations do not necessarily indicate identical cell types were recovered from both porcine ileum and reference data. Abbreviations: ILC (innate lymphoid cell); IQR (interquartile range); LTi (lymphoid tissue inducer); NK (natural killer); NKT (natural killer T); PBMC (peripheral blood mononuclear cell); scRNA-seq (single-cell RNA sequencing); t-SNE (t-distributed stochastic neighbor embedding); T_FH_ (T follicular helper); T_reg_ (T regulatory)

**Figure 3.**
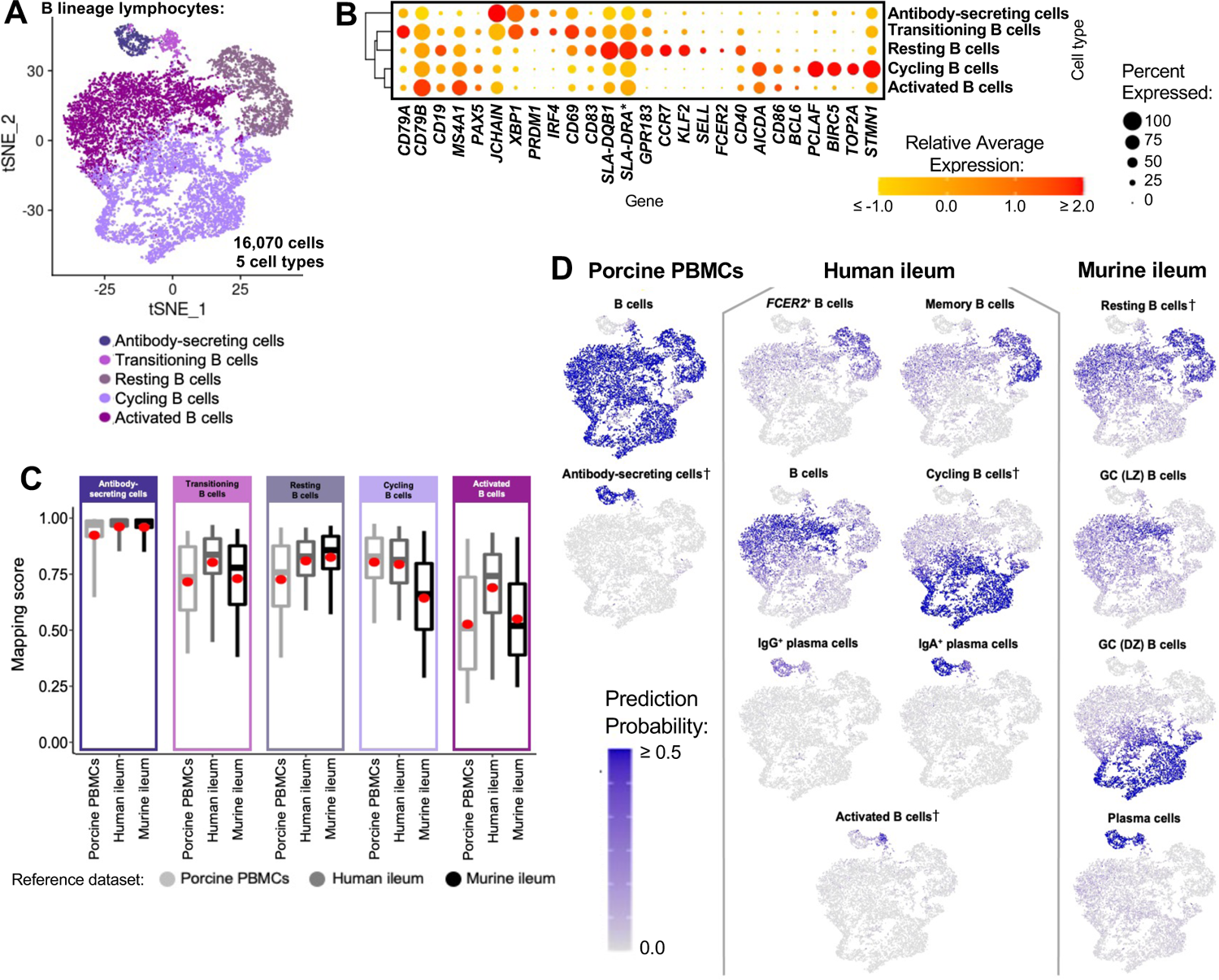
scRNA-seq profiles of B lineage lymphocytes in porcine ileum. **(A)** Two-dimensional t-SNE visualization of 16,070 cells recovered from porcine ileum via scRNA-seq that were classified as B lineage lymphocytes in Figure 1C & **Supplement to Figure 1-Figure 4B** and further annotated into five cell types in Figure 1D & **Supplement to Figure 1 - Figure 9**. Each point represents a single cell; the color of each point indicates cell types shown in Figure 1D. **(B)** Hierarchical relationship of B lineage lymphocyte cell types from porcine ileum shown in a dendrogram (left) and expression patterns of selected genes within each cell type shown in a dot plot (right). In the dot plot, selected genes are listed on the x-axis, and cell types are listed on the y-axis. Within the dot plot, size of a dot corresponds to the percentage of cells expressing a gene within an annotated cell type; color of a dot corresponds to average expression level of a gene for those cells expressing it within a cell type relative to all other cells in the dataset shown in **A**. **(C)** Box plots of the distribution of mapping scores for B lineage lymphocyte cell types from porcine ileum mapped to each reference scRNA-seq dataset. Results for a single cell type are located within a single box, with color of the box corresponding to colors used for cell types in **A**. The color of each box in a plot corresponds to the reference dataset porcine ileal cells were mapped to, including porcine PBMCs (light grey), human ileum (medium grey), and murine ileum (black). Boxes span the interquartile range (IQR) of the data (25^th^ and 75^th^ percentiles), with the median (50^th^ percentile) indicated by a horizontal line. Whiskers span the 5^th^ and 95^th^ percentiles of the data. A red dot represents the data mean. **(D)** Prediction probabilities for porcine ileum scRNA-seq query data from label transfer of selected annotated B/antibody-secreting cell types in reference scRNA-seq datasets of porcine PBMCs (left), human ileum (middle), and murine ileum (right) overlaid onto two-dimensional t-SNE visualization shown in **A**. Each point represents a single cell; the color of each point indicates prediction probability to a corresponding cell type from reference data, as indicated directly above each t-SNE plot. A higher prediction probability indicates higher similarity to a specified annotated cell type in a reference scRNA-seq dataset. scRNA-seq data shown in **A-D** were derived from ileum of two seven-week-old pigs. *Ensembl identifiers found in gene annotation were converted to gene symbols; refer to methods section ‘*Gene name modifications*’ for more details † Identical cell type annotations were given to cells in both porcine ileum and a reference scRNA-seq dataset. Cell type annotations were given to each dataset by independent rationales, and identical annotations do not necessarily indicate identical cell types were recovered from both porcine ileum and reference data. Abbreviations: DZ (dark zone); GC (germinal center); IQR (interquartile range); LZ (light zone); PBMC (peripheral blood mononuclear cell); scRNA-seq (single-cell RNA sequencing); t-SNE (t-distributed stochastic neighbor embedding)

**Figure 4.**
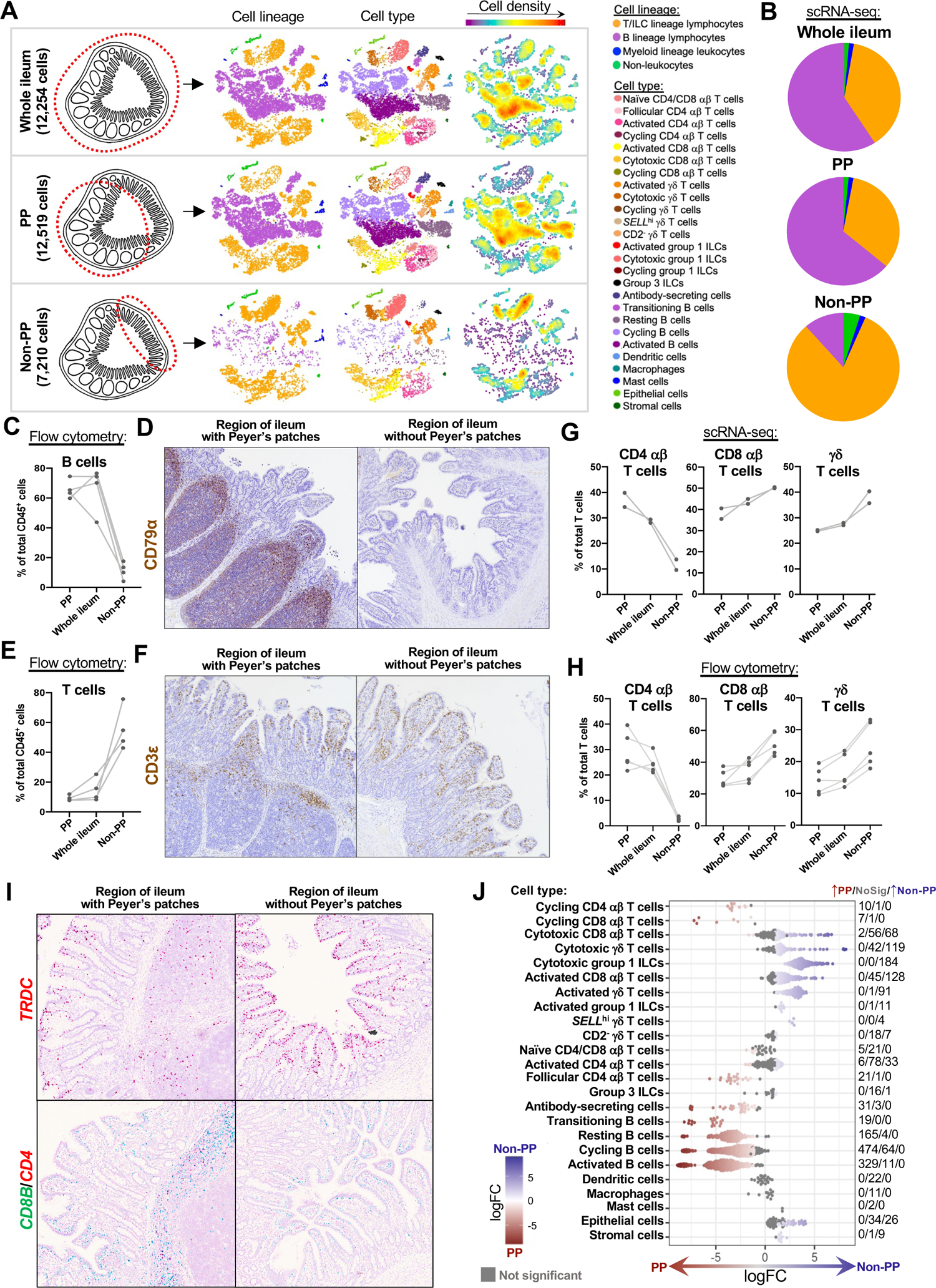
Compositional differences in lymphocytes from ileum with or without Peyer’s patches. **(A)** Cell compositions of scRNA-seq data from whole ileum (top), PP (middle), and non-PP (bottom) samples. Cells from each sample type (depicted on the far left) were combined from a total of two animals and overlaid onto t-SNE coordinates originally presented in Figure 1B-D. The total numbers of cells derived from the total of two animals for each sample type are listed on the far left. On the t-SNE plots, each point represents a single cell; the color of each point corresponds to cell lineage (left t-SNE), cell type (center t-SNE), or cell density (right t-SNE). **(B)** Pie charts showing proportions of cells from each annotated cell lineage within total cells derived from each sample type in **A**. The color of a pie slice indicates cell lineage. The total area of each pie chart is not proportional to the total number of cells derived from each sample type. Proportions were calculated from total cells derived from two pigs for each sample type. **(C)** Plot of the percentage of B cells (CD79α^+^) within total leukocytes (CD45^+^; y-axis) from samples of whole ileum, PP, and non-PP (x-axis), as assessed by flow cytometry gating shown in **Supplement to Figure 4 - Figure 2A**. Measurements from different sample types derived from the same animal are connected by a light grey line. **(D)** IHC staining for B cell CD79α protein (brown) in a region of ileum with Peyer’s patches (left) or without Peyer’s patches (right). **(E)** Plot of the percentage of T cells (CD3ε^+^) within total leukocytes (CD45^+^; y-axis) from samples of whole ileum, PP, and non-PP (x-axis), as assessed by flow cytometry gating shown in **Supplement to Figure 4-Figure 2A**. Measurements from different sample types derived from the same animal are connected by a light grey line. **(F)** IHC staining for T cell CD3ε protein (brown) in a region of ileum with Peyer’s patches (left) or without Peyer’s patches (right). **(G)** Plot of the percentage of CD4 αβ T cells (left), CD8 αβ T cells (center), or γδ T cells (right) within total T cells (y-axis) of the porcine ileum scRNA-seq dataset. Percentages from samples of whole ileum, PP, and non-PP are shown on the x-axis. CD4 αβ T cells included cells annotated as follicular CD4 αβ T cells, activated CD4 αβ T cells, or cycling CD4 αβ T cells and cells annotated as naïve CD4/CD8 αβ T cells with prediction probability to porcine PBMC CD4^+^ αβ T cells > prediction probability to porcine PBMC CD8αβ^+^ αβ T cells. CD8 αβ T cells included cells annotated as activated CD8 αβ T cells, cytotoxic CD8 αβ T cells, or cycling CD8 αβ T cells and cells annotated as naïve CD4/CD8 αβ T cells with prediction probability to porcine PBMC CD8αβ^+^ αβ T cells > prediction probability to porcine PBMC CD4^+^ αβ T cells. γδ T cells included cells annotated as activated γδ T cells, cytotoxic γδ T cells, cycling γδ T cells, *SELL*^hi^ γδ T cells, and CD2^-^ γδ T cells. Measurements from different sample types derived from the same animal are connected by a light grey line. **(H)** Plot of the percentage of CD4 αβ T cells (γδTCR^-^CD4^+^; left), CD8 αβ T cells (γδTCR^-^ CD8β^+^; center), or γδ T cells (γδTCR^+^; right) within total T cells (CD3ε^+^; y-axis) from samples of whole ileum, PP, and non-PP (x-axis), as assessed by flow cytometry gating shown in **Supplement to Figure 4-Figure 2B**. Measurements from different sample types derived from the same animal are connected by a light grey line. **(I)** RNA ISH staining for *TRDC* (top, red), *CD8B* (bottom, green), or *CD4* (bottom, red) transcripts in regions of ileum with Peyer’s patches (left) or regions of ileum without Peyer’s patches (right). **(J)** Differential abundance analysis of cell types from porcine ileum scRNA-seq PP versus non-PP samples. Annotated cell types are listed on the y-axis. Each point represents an individual cell neighborhood, where a neighborhood was assigned as a specific cell type if >70% of cells within the neighborhood belonged to the specified cell type annotation. Cell neighborhoods with <70% of cells belonging to a single cell type are not shown. Grey points indicate cell neighborhoods that were not significantly more abundant in a specific sample type. Non-grey points indicate cell neighborhoods exhibiting differential abundance (p<0.1), and red/blue fill of differentially abundant points corresponds to the magnitude and direction of logFC (also corresponding to values listed on the x-axis). Red indicates increased abundance in PP samples, while blue indicates increased abundance in non-PP samples. On the far right, counts of cell neighborhoods with increased abundance in PP samples/no differential abundance/increased abundance in non-PP samples are shown for each cell type. Cycling γδ T cells and cycling group 1 ILCs are not shown on the y-axis due to no cell neighborhoods being assigned to these cell types. scRNA-seq data shown in **A-B, G, & J** were derived from ileum of two seven-week-old pigs. Images shown in **I** were also taken from a seven-week-old pig used for ileum scRNA-seq. Flow cytometry and IHC experiments were not performed on animals used for scRNA-seq. Flow cytometry experiments shown in **C** & **E** were conducted using four six-week-old pigs. Flow cytometry data shown in **H** was performed using five nine-week-old pigs. IHC staining in **D** was completed on a six-week-old pig. IHC staining in **F** was completed on a nine-week-old pig. Abbreviations: IHC (immunohistochemistry); ILC (innate lymphoid cell); ISH (*in situ* hybridization); logFC (log fold-change); No Sig (no significance); PBMC (peripheral blood mononuclear cell); PP (Peyer’s patch); scRNA-seq (single-cell RNA sequencing); t-SNE (t-distributed stochastic neighbor embedding); TCR (T cell receptor)

Annotated porcine ileal cells were next treated as query data for comparison to existing scRNA-seq datasets (as described in materials and methods section, *‘Reference-based label transfer and mapping*’) to provide greater insight into annotated cell identities. Since a comparable porcine intestinal scRNA-seq dataset was not available, scRNA-seq reference data of healthy porcine PBMCs (Herrera-Uribe & Wiarda et al., 2021), human ileum (Elmentaite & Ross et al., 2020), and murine ileum (Xu et al., 2019) were utilized to provide intra-species/inter-tissue and inter-species/intra-tissue comparisons. Degree of similarity between query and reference cells was determined by calculating mapping scores via reference-based cell mapping (**Supplement to Figure 1-Figure 11**). Transfer of cell labels from reference onto query single-cells provided prediction probabilities to cell types described in each reference dataset (**Supplement to Figure 1-Figure 12**). Gene expression profiles, enrichment of biological processes, and reference-based mapping and cell type prediction results for lymphocytes are presented in the next two results sections.

The main purpose of this work was to deeply characterize porcine ileal lymphocytes, and the majority of cells across all six ileal samples were annotated as belonging to B (50.25%) or T/ILC (46.09%) lymphocyte lineages. However, some non-lymphocytes were also identified, including myeloid lineage leukocytes (1.43%) and non-leukocytes (2.23%). Myeloid lineage leukocytes were composed of dendritic cells (DCs; 264 cells), macrophages (132 cells), and mast cells (62 cells). Identified non-leukocytes included epithelial (607 cells) and stromal (106 cells) cells. Since characterization of non-lymphocytes was not our primary intent for this work, non-leukocytes are not discussed further, but data is available for deeper inquiry (see **Supplement to Figure 1-Figure 10**; **Supplementary Data 6-7**; and our data availability statement).

### Defining the porcine ileal immune landscape: T cells and ILCs

Similar to scRNA-seq results described elsewhere (Elmentaite et al., 2021; Guo et al., 2021; Herrera-Uribe & Wiarda et al., 2021; Patel et al., 2021; Zhao et al., 2020), T cells and ILCs were so transcriptionally similar to one another that they were annotated into a single cell lineage and further resolved into 16 cell types (**Figure 2A**; **Supplement to Figure 2-Figure 1**). T cells were identified by expression of the porcine pan-T cell marker *CD3E* (reviewed by Gerner et al., 2009; Piriou-Guzylack and Salmon, 2008) and included subsets of CD4 αβ, CD8 αβ, and γδ T cells expressing *CD4, CD8B,* and *TRDC*, respectively. ILCs largely lacked *CD3E* and included subsets of group 1 and group 3 ILCs based on expression of genes associated with type 1 or type 3 immunity, respectively (described in subsequent cell type descriptions below; **Figure 2B**). By hierarchical analysis, T/ILC types were more closely related by inferred function (e.g. cell cycling, activation, cytotoxicity) rather than traditional T/ILC phenotypes (e.g. CD4 αβ T cells, CD8 αβ T cells, γδ T cells, group 1 ILCs, group 3 ILCs; **Figure 2B**), which are classically defined based on expression of a series of cell surface markers.

### Cycling T cells and ILCs

One hierarchical grouping of T/ILC types in **Figure 2B** was composed of all cycling T/ILCs, including cycling CD4 αβ T, CD8 αβ T, γδ T, and group 1 ILCs. All cycling cells had significantly increased expression of genes associated with replication/division (e.g. *PCLAF, BIRC5, TOP2A, STMN1* (Dabydeen et al., 2019; Giotti et al., 2019) and enrichment of related biological processes (e.g. establishment of mitotic spindle orientation [*GO:0000132*], regulation of mitotic centrosome separation [*GO:0046602*], DNA duplex unwinding [*GO:0032508*]) relative to other T/ILC types (**Figure 2B**; **Supplementary Data 8**). Cycling T/ILCs had highest average mapping scores to reference porcine PBMCs (≥0.737), followed by human ileum (≥0.699) and murine ileum (≥0.664; **Figure 2C**; **Supplement to Figure 2-Figure 2**). Though predictions of many cycling T/ILC types were to similarly-annotated T/ILCs in reference datasets (e.g. porcine ileal cycling CD4 αβ T cells having highest average predictions to reference CD4 αβ T cell types; **Figure 2D**; **Supplement to Figure 2-Figures 3-5**), several cycling T/ILC types had high prediction to B cells in porcine PBMCs or cycling B cells in human ileum (**Supplement to Figure 2-Figures 3-4**). For instance, cycling CD8 αβ T, γδ T, and group 1 ILCs all had first or second highest average prediction probabilities to cycling B cells in human ileum (**Supplement to Figure 2-Figure 4**), indicating cycling T/ILCs share transcriptional similarities to cycling B cells in human ileum, likely due to shared replication/division-specific gene expression as opposed to shared expression of genes involved in lymphocyte lineage-specific immune functions of the cell.

**Figure 5.**
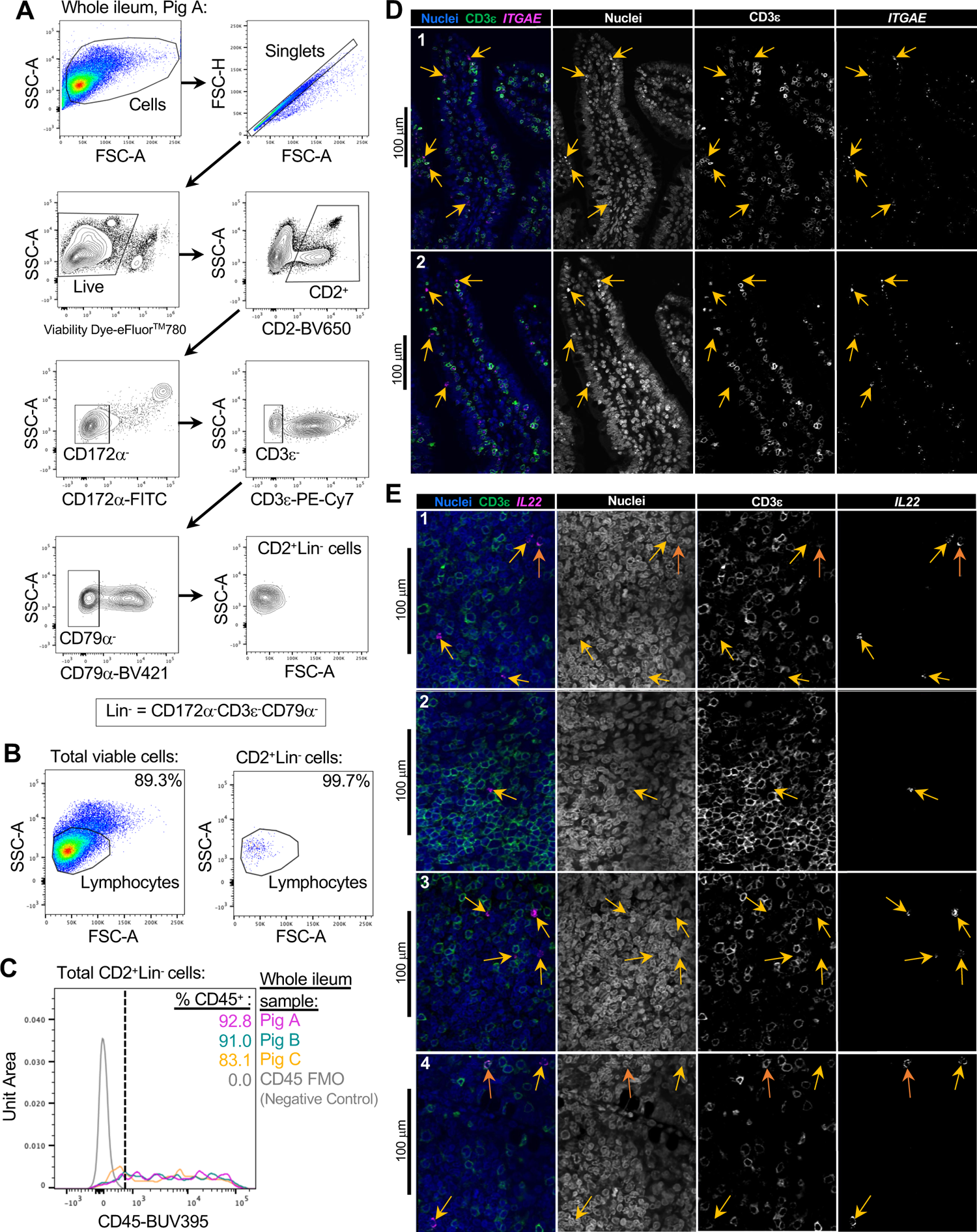
*Ex vivo* and *in situ* identification of ILCs in porcine ileum. **(A)** Flow cytometry gating strategy used to identify CD2^+^Lin^-^ (Lin^-^ = CD172α^-^CD3ε^-^CD79α^-^) within total viable cells of porcine ileal samples. Gating is shown for a whole ileum sample (containing both regions with and without Peyer’s patches) for pig A (corresponding to pig IDs in **C**). **(B)** Flow cytometry forward- and side-scatter plots of total viable cells (left) and CD2^+^Lin^-^ cells (right) within a sample of porcine whole ileum shown in **A**. A gate identifying cells with scatter profiles consistent with lymphocytes is shown, with percentages of cells within the lymphocyte gate listed in the top right of each plot. **(C)** Histogram of the percentage of CD45^+^ cells within CD2^+^Lin^-^ cells identified from samples of porcine ileum using the flow cytometry gating strategy shown in **A**. A fluorescence-minus-one (FMO) sample lacking α-CD45 antibody staining was used as a negative control. **(D)** Dual fluorescent staining of CD3ε protein and *ITGAE* RNA in villi (epithelium + lamina propria) of porcine ileum. Left column: overlay of all stains, including CD3ε protein (green), *ITGAE* RNA (magenta), and nuclei (DAPI staining; blue). Additional columns show individual stain overlays in white, including (from left to right) nuclei, CD3ε protein, and *ITGAE* RNA. Panels of two separate villi are shown in each row. Panels were selected from larger stitched images as shown in **Supplement to Figure 5-Figure 2A**. Yellow arrows indicate location of *ITGAE*^+^CD3ε^-^ cells. **(E)** Dual fluorescent staining of CD3ε protein and *IL22* RNA in lamina propria/GALT of porcine ileum. Left column: overlay of all stains, including CD3ε protein (green), *IL22* RNA (magenta), and nuclei (DAPI staining; blue). Additional columns show individual stain overlays in white, including (from left to right) nuclei, CD3ε protein, and *IL22* RNA. Panels of four separate tissue locations are shown in each row. Panels were selected from larger stitched images as shown in **Supplement to Figure 5-Figure 2B**. Yellow arrows indicate location of *IL22*^+^CD3ε^-^ cells. Orange arrows indicate location of *IL22*^+^CD3ε^+^ cells. Flow cytometry experiments shown in **A-C** were conducted using three six-week-old pigs. Dual IF/ISH experiments shown in **D-E** were conducted using a seven-week-old pig used for ileum scRNA-seq. Abbreviations: FMO (fluorescence-minus-one); FSC-A (forward scatter area); FSC-H (forward scatter height); GALT (gut-associated lymphoid tissue); IF (immunofluorescence); ISH (*in situ* hybridization); ILC (innate lymphoid cell); scRNA-seq (single-cell RNA sequencing); SSC-A (side scatter area)

### Cytotoxic T cells and ILCs

Cytotoxic CD8 αβ T, γδ T, and group 1 ILCs were most closely related to one another and had significantly elevated expression of genes encoding for cytotoxic molecules, including *GZMA*\*, *GZMB,* and *GNLY* (Hidalgo et al., 2008), relative to other T/ILC types (**Figure 2B**; **Supplementary Data 8**). The biological process leukocyte mediated cytotoxicity (*GO:0001909*) was enriched in cytotoxic CD8 αβ T cells, while regulation of natural killer cell mediated cytotoxicity (*GO:0042269*) was enriched in cytotoxic γδ T and group 1 ILCs (**Supplementary Data 8**). Cytotoxic cell types had some of the lowest average mapping scores to reference porcine PBMCs (range of means 0.645 to 0.732), indicating dissimilarity between cytotoxic ileal cells from any cells in circulation (**Figure 2C**; **Supplement to Figure 2-Figure 2**). Though cytotoxic cell types had lower mapping scores to porcine PBMCs, cytotoxic CD8 αβ T cells and group 1 ILCs still had highest prediction to comparable cell types in porcine peripheral blood: CD8αβ^+^ αβ T cells and NK cells, respectively (**Figure 2D**; **Supplement to Figure 2-Figure 3**). Ileal cytotoxic γδ T cells had highest average prediction to innate-like CD8α^+^ αβ T cells and NK cells from porcine peripheral blood rather than to peripheral CD2^+^ γδ T cells, further supporting poor representation of ileal cytotoxic γδ T cells by porcine peripheral γδ T cells and suggesting greater similarities to other peripheral innate or innate-like T/ILC types instead (**Figure 2D**; **Supplement to Figure 2-Figure 3**). A similar pattern was observed in murine ileum, where porcine ileal cytotoxic γδ T cells had highest prediction to reference NK cells rather than *Gzma*+γδ T cells, suggesting again that porcine ileal cytotoxic γδ T cells had greater similarity to other innate/innate-like T/ILC types rather than γδ T cells (**Figure 2D**; **Supplement to Figure 2-Figure 5**).

### Activated γδ T cells, CD8 αβ T cells, and group 1 ILCs

Activated γδ T, CD8 αβ T, and group 1 ILCs also formed a hierarchical grouping closely related to cytotoxic cell counterparts in **Figure 2B**. In contrast to cytotoxic T/ILCs, activated γδ T, CD8 αβ T, and group 1 ILCs had lower expression of genes encoding cytotoxic molecules (e.g. *GZMA*, GZMB, GNLY*) but significantly elevated expression of other genes indicative of previous or recent cell activation, including *CTSW, XCL1, SLA-DRA*, SLA-DQB1,* and *CCR9* (**Figure 2B**; **Supplementary Data 8**; Boismenu et al., 1996; Gerner et al., 2009; Iwata et al., 2004; Kelner Gregory et al., 1994; Ondr and Pham, 2004; Stoeckle et al., 2009; Svensson et al., 2002; Uehara et al., 2002). Activated γδ T, CD8 αβ T, and group 1 ILCs were all enriched for the biological processes positive regulation of T cell differentiation (*GO:0045582*) and positive regulation of T cell mediated immunity (*GO:0002711*), further supporting an activated cell state (**Supplementary Data 8**). Activated γδ T, CD8 αβ T, and group 1 ILCs had higher average mapping scores to all reference datasets (range of means 0.872 to 0.902) than did corresponding cytotoxic T/ILCs, indicating better representation by reference data (**Figure 2C**; **Supplement to Figure 2-Figure 2**). Unlike cytotoxic γδ T cells, activated γδ T cells had highest prediction to γδ T cell types in reference datasets, including CD2^+^ γδ T cells (porcine PBMCs), γδ T/NK cells (human ileum), and *Xcl1*+ γδ T cells (murine ileum), suggesting greater similarity of porcine ileal activated γδ T cells to reference γδ τ cell populations than observed for porcine ileal cytotoxic γδ T cells (**Figure 2D**; **Supplement to Figure 2-Figures 3-5**). Activated CD8 αβ T cells had highest average predictions to CD8 αβ T cell types in reference datasets, as did activated group 1 ILCs to reference ILC types (**Figure 2D**; **Supplement to Figure 2-Figures 3-5**). Though both activated and cytotoxic group 1 ILCs had highest prediction to reference group 1 ILC types, activated group 1 ILCs had highest prediction to porcine peripheral CD8α^+^ αβ T/NK cells and murine ileal ILC1s. In contrast, cytotoxic group 1 ILCs had highest prediction to NK cells in the same reference datasets, delineating transcriptional distinctions between cytotoxic and activated group 1 ILCs that correspond better to different reference cell types (**Figure 2D**; **Supplement to Figure 2-Figures 3, 5**).

### SELL^hi^ γδ T cells

*SELL*^hi^ γδ T cells were a minor fraction of porcine ileal γδ T cells (91 cells total) that shared a node with cytotoxic and activated T/ILCs expressing effector/activation molecules including *CCL5* and *ITGAE* (**Figure 2B**; Ling et al., 2007; Szabo et al., 2019a)*. SELL*^hi^ γδ T cells nearly ubiquitously expressed *SELL* (encoding CD62L) and genes related to cytotoxicity (e.g. *GZMA*, GZMB*), but also some genes expressed by activated T/ILCs, such as *XCL1* (**Figure 2B**). *SELL*^hi^ γδ T cells expressed genes encoding innate receptors, including *FCER1G* and *KLRG1*,* but lacked expression of others, such as *KLRK1* (**Figure 2B**). Moreover, *SELL*^hi^ γδ T cells had significantly higher expression of genes encoding for adhesion molecules (*SELL, ITGB1, ITGB7*), and the transcriptional regulator and γδ T cell fate determinator, *ID3* (**Figure 2B**; **Supplementary Data 8**; Lauritsen et al., 2009). Five of the top eight enriched biological processes for *SELL*^hi^ γδ T cells (as determined by smallest p-values) included actin filament depolymerization (*GO:0030042*), positive regulation of actin filament polymerization (*GO:003038*), establishment or maintenance of cell polarity (*GO:0007163*), integrin-mediated signaling pathway (*GO:0007229*), and natural killer cell activation (*GO:0030101*), indicating a highly activated state potentially related to cell receptor engagement/signaling (**Supplementary Data 8**). *SELL*^hi^ γδ T cells had the lowest average mapping scores of all cell types in comparison to each reference dataset (range of means 0.473 to 0.636; **Figure 2C**; **Supplement to Figure 2-Figure 2**), suggesting they were unique to porcine ileum.

### CD2^-^ γδ T cells

CD2^-^ γδ T cells (characterized as *TRDC*-expressing cells that lacked *CD2* expression; **Figure 2B**) are a unique porcine γδ T cell subset, as they are absent from humans and mice (Stepanova and Sinkora, 2013). Correspondingly, CD2^-^ γδ T cells had higher average mapping scores to porcine PBMCs (0.959) than to ileal cells from human (0.796) or mouse (0.832) (**Figure 2C**; **Supplement to Figure 2-Figure 2**). CD2^-^ γδ T cells had the highest average mapping scores to porcine PBMCs of all T/ILC types, indicating CD2^-^ γδ T cells to be the porcine ileal T/ILC type best represented by cells in the porcine periphery. CD2^-^ γδ T cells are considered naïve cells in pigs (Käser, 2021; Rodríguez-Gómez et al., 2019; Sedlak et al., 2014), and, in support, CD2^-^ γδ T cells were most closely related to naïve CD4/CD8 αβ T cells in porcine ileum by hierarchical clustering (**Figure 2B**). Besides lacking *CD2* expression, ileal CD2^-^ γδ T cells had significantly elevated expression of *RHEX, BLK, SAMSN1,* and *IL26* (**Figure 2B**; **Supplementary Data 8**), which were also highly expressed by CD2^-^ γδ T cells in the porcine periphery (Herrera-Uribe & Wiarda et al., 2021). CD2^-^ γδ T cells were predicted most similar to corresponding CD2^-^ γδ T cells in porcine peripheral blood and to γδ T/NK cells in human ileum, while in murine ileum, predictions were lowly distributed across multiple T/ILC subsets (**Figure 2D**; **Supplement to Figure 2-Figures 3-5**). Thus, CD2^-^ γδ T cells can be found in both ileum and periphery of pigs and are less well represented in human or murine ileum.

### Naïve CD4/CD8 αβ T cells

Naïve CD4 and CD8 αβ T cells had significantly higher expression of genes related to cell circulation and a naïve T cell phenotype, including *CCR7, S1PR1, LEF1, KLF2* (**Figure 2B**; **Supplementary Data 8**; Cano-Gamez et al., 2020; Sebzda et al., 2008; Shan et al., 2021; Skon et al., 2013; Willinger et al., 2006). Of all T/ILCs, naïve CD4/CD8 αβ T cells had the second-highest average mapping scores to porcine PBMCs (0.944; **Figure 2C**; **Supplement to Figure 2-Figure 2**), indicating naïve CD4 and CD8 αβ T cells to be the porcine ileal T/ILC type second-best represented by cells in the porcine periphery, trailing only behind CD2^-^ γδ T cells with a similar naïve T cell state. High mapping scores to human and murine ileum (means 0.921 and 0.917, respectively) were also noted, indicating good representation of naïve CD4 and CD8 αβ T cells in ileum of both human and mouse. Porcine ileal naïve CD4/CD8 αβ T cells had highest prediction to corresponding populations in reference datasets, including CD4 and CD8 αβ T cell populations derived from porcine PBMCs or human ileum and resting CD4 and CD8 T cells derived from murine ileum (**Figure 2D**; **Supplement to Figure 2-Figure 3-5**).

### Activated and follicular CD4 αβ T cells

Remaining non-naïve/non-cycling CD4 αβ T cells in porcine ileum included follicular and activated CD4 αβ T cells, which were most closely related to one another (**Figure 2B**). Activated CD4 αβ T cells did not share elevated expression of several genes highly expressed by other activated T/ILC types (e.g. *CCL5, ITGAE, CTSW, XCL1, SLA-DRA*, SLA-DQB1, CCR9*) but instead had significantly elevated expression of genes associated with CD4 αβ T cell activation (e.g. *ICOS, CTLA4,* and *CD40LG*; Cano-Gamez et al., 2020; Hutloff et al., 1999; Jaiswal et al., 1996; Linsley and Golstein, 1996; Miragaia et al., 2019), which were also elevated in follicular CD4 αβ T cells (**Figure 2B**; **Supplementary Data 8**). However, activated CD4 αβ T cells had higher expression of activation-associated genes *IFITM3* and *GPR183* (Bedford et al., 2019; Clottu et al., 2017; Szabo et al., 2019a) relative to follicular CD4 αβ T cells. Follicular CD4 αβ T cells were characterized by higher expression of *PDCD1, CXCR4, CD69,* and *LTB*, all genes highly expressed by follicle-associated T cells (Haynes et al., 2007; Schaerli et al., 2000; Shi et al., 2018), such as T follicular helper (T_FH_) or T follicular regulatory (T_FR_) cells (**Figure 2B**). The top two enriched biological processes in follicular CD4 αβ T cells (smallest p-values) were related to B cell activation/humoral immunity, including humoral immune response mediated by circulating immunoglobulin (*GO:0002455*) and plasma cell differentiation (*GO:0002317*; **Supplementary Data 8**). Follicular CD4 αβ T cells had lower mapping scores to porcine PBMCs (mean 0.733) than did activated CD4 αβ T cells (mean 0.880; **Figure 2C**; **Supplement to Figure 2-Figure 3**), indicating greater dissimilarity of follicular CD4 αβ T cells than activated CD4 αβ T cells to circulating cells in pigs. Porcine follicular CD4 αβ T cells had highest prediction to T_FH_ cells in human ileum and activated CD4 T cells in murine ileum (**Figure 2D**; **Supplement to Figure 2-Figure 4-5**), further supporting an activated role associated with follicular helper/regulatory functions. Activated CD4 αβ T cells were largely predicted as activated CD4 T cells in murine ileum and more so as CD4 T than T_FH_ in human ileum (**Figure 2D**; **Supplement to Figure 2-Figure 4-5**), supporting an activated cell state.

### Group 3 ILCs

Group 3 ILCs expressed many genes characteristic of type 3 immunity, including *IL22, RORC,* and *CXCL8* (Haynes et al., 2007; Qi et al., 2021; Schaerli et al., 2000; Shi et al., 2018), and were more closely related to non-cycling CD4 αβ and naïve T cell subsets than to any type of group 1 ILC (**Figure 2B**). Though ILCs largely lacked expression of pan-T cell marker *CD3E*, group 1 ILCs still expressed other CD3 complex-associated genes, such as *CD3G* and *CD247* (encoding CD3γ and CD3ζ, respectively). In contrast, group 3 ILCs largely lacked expression of all aforementioned CD3 subunit-encoding genes and also had significantly higher expression of classical ILC gene markers, including *KIT, ID2, IL7R,* and *KLRB1* (**Figure 2B**; **Supplementary Data 8**; Boos et al., 2007; Satoh-Takayama et al., 2010; Spits et al., 2013; Yokota et al., 1999; Yoshida et al., 1999), though these markers are already known to be variably expressed by intestinal group 1 ILCs based on species and regional location (Meininger et al., 2020; Robinette et al., 2015; Simoni et al., 2017; Simoni and Newell, 2017; Van Acker et al., 2017). Group 3 ILCs mapped best to cells of the murine ileum (mean mapping score 0.979) and were predicted most similar to corresponding group 3 ILC populations of ILC3s or lymphoid tissue inducer (LTi) cells (**Figure 2C-D**; **Supplement to Figure 2-Figure 5**). In contrast, group 3 ILCs did not have as close a counterpart in porcine PBMCs or human ileum, as indicated by lower average mapping scores (0.632 and 0.754, respectively) and prediction most similar to CD4 αβ T cells or activated T cells, respectively (**Figure 2C-D**; **Supplement to Figure 2-Figures 3-4**).

### Defining the porcine ileal immune landscape: B cells and antibody-secreting cells

B lineage lymphocytes were annotated as antibody-secreting cells (ASCs), B cells transitioning into ASCs (referred to as transitioning B cells), and three additional populations of B cells, including resting, cycling, and activated B cells (**Fig. 3A**; **Supplement to Figure 1-Figure 9G**).

### Antibody-secreting cells

ASCs were most distantly related from other B cell types by hierarchical clustering and had significantly lower expression of several canonical B cell genes, including *CD19, CD79A, CD79B, MS4A1, PAX5* (**Figure 3B**; **Supplementary Data 9**; Herrera-Uribe & Wiarda et al., 2021; Lee et al., 2021). Genes known to be expressed by porcine peripheral ASCs (e.g. *JCHAIN, XBP1, IRF4, PRDM1*; Herrera-Uribe & Wiarda et al., 2021) had elevated expression in ileal ASCs as well. The top two enriched biological processes in ASCs relative to other B cells were related to B cell activation (positive regulation of B cell activation [*GO:0050871*]) and protein production, such as required for producing antibodies (positive regulation of protein exit from endoplasmic reticulum [*GO:0070863*]; **Supplementary Data 9**). ASCs were well-represented by all reference datasets, as indicated by high mapping scores (means ≥0.927; **Figure 3C**; **Supplement to Figure 3-Figure 1**), and were almost unanimously predicted as ASC/plasma cell types from all reference datasets (**Figure 3D**; **Supplement to Figure 3-Figures 2-4**).

### Transitioning B cells

Similar to ASCs, transitioning B cells had high expression of genes characteristic of porcine ASCs, including *JCHAIN, XBP1, IRF4, PRDM1,* and were enriched for biological processes supporting antibody production, including the top three enriched processes of posttranslational protein targeting to endoplasmic reticulum membrane (*GO:0006620*), protein N-linked glycosylation (*GO:0006487*), and glycoprotein catabolic process (*GO:0006516*; **Figure 3B**; **Supplementary Data 9**). Porcine ileal ASCs had highest average prediction to ASC/plasma cell types in reference datasets; however, prediction scores to ASC/plasma cell types for transitioning B cells were lower than those observed in ASCs, and transitioning B cells also had high prediction to activated B cells in human ileum (**Figure 3D**; **Supplement to Figure 3-Figures 2-4**). Transitioning B cells had lower average mapping scores to all reference datasets than did ASCs (means ≥0.718; **Figure 3C**; **Supplement to Figure 3-Figure 1**), indicating poorer representation by the reference data. In contrast to ASCs, transitioning B cells had greater expression of canonical B cell genes (*CD19, CD79A, CD79B, MS4A1, PAX5*) and higher expression of several markers of early B cell activation, including *CD69, CD83, SLA-DQB1,* and *SLA-DRA\** (**Figure 3B**; Ashouri and Weiss, 2017; Breloer et al., 2007; Prazma et al., 2007; Rahe and Murtaugh, 2017; Van der Stede et al., 2005), supporting functional inference that cells were a subset of more recently activated B cells transitioning to produce and secrete antibody.

### Resting B cells

Remaining B cell types (resting, cycling, activated B cells) all had greater expression of B cell canonical genes (*CD19, CD79A, CD79B, MS4A1, PAX5*) than did ASCs and lesser expression of aforementioned genes expressed by both ASCs and transitioning B cells (*JCHAIN, XBP1, IRF4, PRDM1*; **Figure 3B**). Resting B cells were most closely related to transitioning B cells in porcine ileum but, unlike remaining cycling/activated B cell subsets, lacked expression of several genes associated with activation and/or germinal centers, including *AICDA, BCL6, CD86* (Allman et al., 1996; Engel et al., 1994; Lee et al., 2021; Muramatsu et al., 1999), indicating cells in a resting state (**Figure 3B**). Resting B cells had increased expression of genes characteristic of cell circulation and naïve/memory B cells, including *KLF2, SELL* (CD62L)*, CCR7, FCER2* (CD23)*, CD40* (**Figure 3B**; Bhattacharya et al., 2007; Förster et al., 1999; Lee et al., 2021; Rahe and Murtaugh, 2017; Waldschmidt et al., 1988; Winkelmann et al., 2011; Zhang et al., 2021a); however, it remained indiscriminate as to whether resting B cells were naïve, memory, or a combination of both, as many of the same genes are expressed by both naïve and memory B cell subsets. In comparison to reference datasets, resting B cells were mostly predicted as memory B cell types (memory or *FCER2^+^* B cells) in human ileum and as resting B cells in murine ileum (**Figure 3D**; **Supplement to Figure 3-Figures 3-4**).

### Activated and cycling B cells

The remaining two B cell types in porcine ileum included cycling and activated B cells, which were most closely related to one another in **Figure 3B**. Both cell types had high expression of genes related to B cell activation and/or germinal center-associated responses (e.g. *AICDA, BCL6, CD86*; Muramatsu et al., 1999; Victora et al., 2010; Ye et al., 1997), but cycling B cells also had characteristics of cellular replication/division, including elevated expression of *PCLAF, BIRC5, TOP2A, STMN1* (Dabydeen et al., 2019; Giotti et al., 2019) and enrichment of biological processes such as nucleosome organization (*GO:0034728*), centriole-centriole cohesion (*GO:0010457*), and mitotic spindle organization (*GO:0007052*; **Figure 3B**; **Supplementary Data 9**). Porcine ileal cycling B cells had highest prediction to cycling B cells in human ileum and germinal center dark zone (GC DZ) B cells in murine ileum, while activated B cells instead had highest prediction to cells labeled as B cells in human ileum and germinal center light zone (GC LZ) or resting B cells in murine ileum (**Figure 3D**; **Supplement to Figure 3-Figures 3-4**). A subset of cycling B cells had higher prediction scores to porcine peripheral T/ILC lineage lymphocytes and were more specifically predicted to be CD8αβ^+^ αβ T cells (**Supplement to Figure 3-Figure 2**). Of all B/ASC types, porcine ileal activated B cells had the lowest average mapping scores to all reference datasets (range of means 0.530 to 0.692), suggesting lack of a similar cell population in porcine circulation and human or murine ileum (**Figure 3C**; **Supplement to Figure 3-Figure 1**).

### B lineage and cycling lymphocytes enriched in ileum containing Peyer’s patches

Since Peyer’s patches are niches of GALT with specialized cellular functions different from those performed by cells in the lamina propria or epithelium, we assessed the impact of inclusion versus exclusion of Peyer’s patches on cellular compositions recovered from porcine ileum. As already shown in **Figure 1A** and **Supplement to Figure 1-Figure 1B**, ileal tissue was dissected into sections with Peyer’s patches (PP), without Peyer’s patches (non-PP), and a whole cross section of ileum (whole ileum) for cell isolation and scRNA-seq. At pseudo-bulk RNA-seq rather than scRNA-seq resolution, overall gene expression profiles of PP and whole ileum were distinct from non-PP samples both before and after data quality control/filtering (**Supplement to Figure 4-Figure 1A**). Analysis at single-cell resolution revealed similar results, whereby cell type proportions and overall cell numbers in whole ileum samples more closely resembled PP than non-PP samples (**Figure 4A**; **Supplement to Figure 4-Figure 1B-C**). At the cell lineage level, whole ileum and PP samples were composed primarily of B lineage lymphocytes (59.12% and 63.89%, respectively), followed by T/ILC lineage lymphocytes (38.13% and 33.17%, respectively; **Figure 4B**). In contrast, most cells from non-PP samples were T/ILC lineage lymphocytes (82.07%), and only 11.45% were B lineage lymphocytes (**Figure 4B**).

Presence and abundance of selected lymphocyte populations in different ileal regions were further validated *ex vivo* and *in situ*. Flow cytometry was used to assess B cell abundance via intracellular CD79α protein expression. Larger proportions of CD45^+^ leukocytes were CD79α^+^ in PP and whole ileum samples when compared to non-PP samples (**Figure 4C**; **Supplement to Figure 4-Figure 2A**). Immunohistochemistry (IHC) labeling revealed CD79α protein primarily in follicular areas of Peyer’s patches but largely absent in lamina propria and epithelium (**Figure 4D**), indicating minimal CD79α detected in regions representative of non-PP samples. Since a dependable marker has not yet been established to identify ILCs in pigs, CD3ε protein staining was performed to label only T cells. By flow cytometry, higher percentages of CD3ε^+^ cells were detected within total CD45^+^ leukocyte populations of non-PP samples compared to PP or whole ileum samples (**Figure 4E**; **Supplement to Figure 4-Figure 2A**). By IHC, CD3ε protein staining was abundant in lamina propria, epithelium, and T cell areas of Peyer’s patches (**Figure 4F**), indicating CD3ε was present in regions representative of all ileal sections (PP, non-PP, whole ileum). Collectively, *ex vivo* and *in situ* staining for CD79α and CD3ε protein supported scRNA-seq observations: B cells comprised a larger proportion of cells in PP and whole ileum samples, while T cells comprised a larger proportion of cells in non-PP samples. These results are informative in deciding sample preparation for inclusion of cells relevant to biological questions under investigation.

We further validated proportions of CD4 αβ, CD8 αβ, and γδ T cells in various regions of ileum using flow cytometry and RNA *in-situ* hybridization (ISH). T cells recovered via scRNA-seq were regrouped into CD4 αβ, CD8 αβ, and γδ T cells, and percentages of each subset within total T cells was calculated for each ileal scRNA-seq sample. Analysis revealed: (1) increased proportions of CD4 αβ T cells in PP versus non-PP samples; (2) increased proportions of CD8 αβ T cells and γδ T cells in non-PP versus PP samples; and (3) intermediate proportions of all three T cell subsets in whole ileum compared to PP and non-PP samples (**Figure 4G**). By flow cytometry, T cell proportions mirrored patterns obtained from scRNA-seq (**Figure 4H**; **Supplement to Figure 4-Figure 2B**). RNA ISH staining in regions of ileum without Peyer’s patches (**Figure 4I**, right) revealed *TRDC* (γδ T cells) and *CD8B* (CD8 αβ T cells) transcripts were primarily expressed within the epithelial layer, while *CD4* (CD4 αβ T cells) was expressed primarily within the lamina propria, supporting the conclusion that most γδ and CD8 αβ T cells were intraepithelial, and most CD4 αβ T cells resided in the lamina propria. Localization of *CD4, CD8B,* and *TRDC* did not change in epithelium and lamina propria adjacent to Peyer’s patches. However, all three transcripts were also expressed by cells in the T cell zones of Peyer’s patches, which were removed from non-PP samples (**Figure 4I**, left). Flow cytometry staining of epithelium-enriched, sub-epithelium-enriched, and merged cell fractions from PP and non-PP ileal samples was performed to validate ISH findings. The results revealed epithelium-enriched fractions from both PP and non-PP samples had higher percentages of γδ and CD8 αβ T cells and lower percentages of CD4 αβ T cells compared to sub-epithelium-enriched cell fractions (**Supplement to Figure 4-Figure 2C**). In all, *ex vivo* and *in situ* staining to identify location of CD4 αβ, CD8 αβ, and γδ T cells mirrored results from scRNA-seq and provided further locational context of T cells in ileal epithelium, lamina propria, and Peyer’s patches.

PP and non-PP samples were dissected by complete inclusion or exclusion of Peyer’s patches, respectively, and could thus be directly compared to identify annotated cell types enriched in the presence versus absence of Peyer’s patches. To identify cell type enrichment, cell neighborhoods (conglomerates of cells located near each other in the multidimensional space of the dataset) were identified from cells of PP and non-PP samples, and differential abundance analysis was performed on cell neighborhoods (**Figure 4J**; **Supplement to Figure 4-Figure 3B**). Though T/ILC lineage lymphocytes comprised a greater proportion of total cells in non-PP than PP samples (**Figure 4B**), several T/ILC types were more abundant in PP samples, including cycling CD4 αβ T cells, cycling CD8 αβ T cells, and follicular CD4 αβ T cells, with at least 87.5% of cell neighborhoods significantly more abundant in PP samples for each cell type (**Figure 4J**). In contrast, cytotoxic and activated γδ T, CD8 αβ T, and group 1 ILCs, along with *SELL*^hi^ γδ T cells, had the majority of cell neighborhoods (>50% for each cell type) significantly enriched in non-PP samples (**Figure 4J**). Remaining T/ILC types had no or lower percentages of differentially abundant cell neighborhoods: 94.1% of group 3 ILC cell neighborhoods were not significantly differentially abundant; CD2^-^ γδ T cells had 28.0% of cell neighborhoods significantly increased in non-PP samples; naïve CD4/CD8 αβ T cells had 19.2% of cell neighborhoods significantly increased in PP samples; and activated CD4 αβ T cells had 5.1% and 28.2% of cell neighborhoods significantly enriched in PP and non-PP samples, respectively (**Figure 4J**). Cycling γδ T and group 1 ILCs did not have any cell neighborhoods recovered for the analysis; however, no cycling group 1 ILCs were recovered from non-PP samples, and cycling γδ T cells derived from PP samples outnumbered those derived from non-PP samples by sixteen-to-three (**Supplement to Figure 4-Figure 1C**). Similar to results evaluating compositions at the cell lineage level, all cell types from the B lymphocyte lineage (ASCs, transitioning B, resting B, cycling B, activated B) had at least 88.1% of cell neighborhoods significantly more abundant in PP samples, indicating enrichment within Peyer’s patches (**Figure 4J**). Collectively, differential abundance analysis indicated B cells, ASCs, cycling T/ILCs, and follicular CD4 αβ T cells were more abundant in PP samples, likely due to association of these cell types with functions of ileal Peyer’s patches and germinal center responses. Cytotoxic and activated subsets of γδ T, CD8 αβ T, and group 1 ILCs were more abundant in non-PP samples, indicating location and functions within the epithelium and/or lamina propria rather than in Peyer’s patches of ileum.

### Differential location of group 1 ILCs and group 3 ILCs in porcine ileum indicated via *ex vivo* and *in situ detection*

To date, NK cells are the only porcine non-B/non-T lymphocyte subset identified, and typically identified as CD3ε^-^CD8α^+^ lymphocytes using *ex vivo* flow cytometry assessment (reviewed by Gerner et al., 2009). NK cell frequency in porcine intestine has been assessed, but very few NK cells are detected in porcine ileum (Annamalai et al., 2019; Annamalai et al., 2015; Potockova et al., 2015; Sinkora et al., 2011; Wasowicz et al., 2018). In contrast, we identified 2,594 cells (8.1% of total cells) as ILCs in the porcine ileum via scRNA-seq, and few ILCs expressed *CD8A* (**Figure 2B**; **Supplement to Figure 5-Figure 1**). We therefore used scRNA-seq data to inform an approach to identify ILCs (not just CD3ε^-^CD8α^+^ NK cells) by flow cytometry.

Further query of ILC single-cell gene expression profiles revealed ILCs expressed *CD2* but lacked gene expression of canonical T, B, and myeloid lineage leukocyte markers (*CD3E, CD79A,* and *SIRPA*\* [encoding CD172α], respectively; **Figure 2B**; **Supplement to Figure 5-Figure 1**). Since porcine-specific immunoreagents are commercially available for CD2, CD3ε, CD79α, and CD172α, we identified cells corresponding to an ILC phenotype from porcine ileum via flow cytometry as CD2^+^Lin^-^ cells (Lin^-^ = CD3ε^-^CD79α^-^CD172α^-^; **Figure 5A**). Nearly all CD2^+^Lin^-^ cells had forward- and side-scatter properties indicative of lymphocytes (**Figure 5B**). Most ILCs also expressed *PTPRC* (encoding CD45, considered to be a pan-leukocyte marker in pigs [reviewed by Piriou-Guzylack and Salmon, 2008]) in our scRNA-seq data.

However, fewer group 3 ILCs expressed *PTPRC* (67.68%; **Supplement to Figure 5-Figure 1**), and previous work in mice indicates CD45 expression may be lost by ILC3s, rendering it an inconsistent ILC marker (Xu et al., 2019). Congruently, the majority (≥83.1%) but not all CD2^+^Lin^-^ cells in the porcine ileum were CD45^+^ (**Figure 5C**).

To assess spatial context of ILCs in tissue and cautiously infer cell function based on location, we developed *in situ* methods for detection of porcine group 1 and group 3 ILCs in ileum. From initial scRNA-seq analysis, activated, cytotoxic, and cycling group 1 ILCs all had high expression of *ITGAE*, encoding the integrin alpha subunit of CD103 that is highly expressed by intraepithelial lymphocytes in the intestine of other species (**Figure 2B**; **Supplement to Figure 5-Figure 1B-C**; Cepek et al., 1994; Kilshaw and Murant, 1990; Mayassi and Jabri, 2018; Olivares-Villagómez and Van Kaer, 2018). Correspondingly, *ITGAE* was primarily detected in the epithelial layer of porcine ileum by *in situ* analysis (**Supplement to Figure 5-Figure 2A**).

*ITGAE* was also strongly expressed by some T cells in our scRNA-seq dataset (**Figure 2B**; **Supplement to Figure 5-Figure 1B-C**), thus we additionally assessed protein expression of CD3ε to decipher intraepithelial T cells (*ITGAE*^+^CD3ε^+^) from intraepithelial group 1 ILCs (*ITGAE*^+^CD3ε^-^). Dual *ITGAE*/CD3ε *in situ* labeling revealed the majority of *ITGAE^+^* intraepithelial cells were CD3ε^+^ T cells; however, *ITGAE*^+^CD3ε^-^ intraepithelial cells were also noted in porcine ileum and thus assumed to be intraepithelial group 1 ILCs (**Figure 5D**; **Supplement to Figure 5-Figure 2A**).

Group 3 ILCs in porcine ileum had high expression of *IL22* (**Figure 2B, Supplement to Figure 5-Figure 1B-C**), and *IL22* was detected *in situ* primarily in the lamina propria and T cell zones of Peyer’s patches in porcine ileum (**Supplement to Figure 5-Figure 2B**). Furthermore, most but not all *IL22*^+^ cells detected *in situ* lacked CD3ε expression, indicating the majority of *IL22*-expressing cells were not T cells (**Figure 5E**; **Supplement to Figure 5-Figure 2B**).

Therefore, we identified group 3 ILCs as *IL22^+^*CD3ε^-^ cells, though the *in situ* staining combination also identified a minority of *IL22^+^*CD3ε^+^ (presumably T_H22_) cells. Collectively, query of gene expression for ILCs at single-cell resolution allowed us to develop new reagent panels for *ex vivo* and *in situ* analysis of ILCs in the porcine ileum that expanded on previous panels catering to only NK cell identification.

### Transcriptional distinctions between porcine ileal and circulating ILCs

Porcine NK cells were not representative of porcine ileal ILCs; therefore, ileal ILCs required different markers for *ex vivo* and *in situ* detection (**Figure 5**; **Supplement to Figure 5-Figure 1-2**). As porcine peripheral NK cells are readily characterized, we wanted to further explore transcriptional distinctions between porcine ileal ILCs and porcine peripheral ILCs (i.e., NK cells) to infer potential cell functions and identify additional targets for *ex vivo* and *in situ* detection. PBMCs from the same two pigs were processed for scRNA-seq in parallel to ileal samples, and peripheral ILCs were identified by the same criteria used to identify ILCs in the ileum (refer to materials and methods for full details). Porcine peripheral ILC gene expression profiles were largely concordant with NK cells described in our previous work (**Supplement to Figure 6-Figure 1A-E**; **Supplementary Data 1**; Herrera-Uribe & Wiarda et al., 2021). A new merged ILC scRNA-seq dataset containing only single cells annotated as ILCs from both PBMCs and ileal samples was generated (**Figure 6A**). Within the merged ILC dataset, 41.3% of detected cell neighborhoods were differentially abundant based on ileal versus peripheral cell origin (**Figure 6B-C**), suggesting differences between at least some ileal and peripheral ILCs may exist and meriting further exploration.

**Figure 6.**
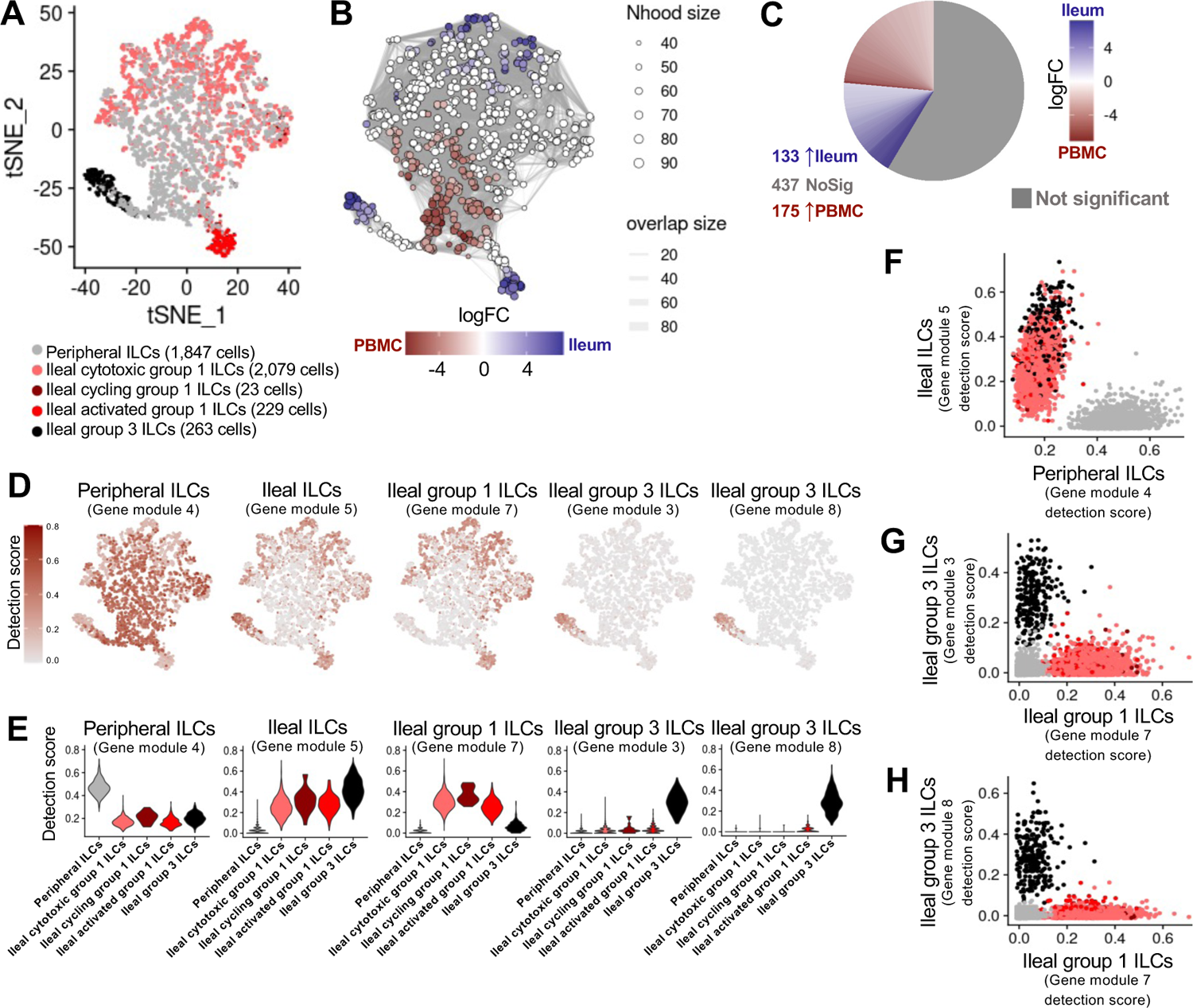
Peripheral ILCs are transcriptionally distinct from ileal ILCs. **(A)** Two-dimensional t-SNE visualization of 4,441 cells recovered from porcine ileum (2,594 cells) and porcine PBMCs (1,847 cells) via scRNA-seq and classified as ILCs in **Supplement to** Figure 1**-**Figure 5C-F & **Supplement to** Figure 6**-Figure D-E**. Each point represents a single cell; color of a point corresponds to one of five ILC annotations. The number of cells belonging to each cell type is listed in the color key below. Derivation from ileum or PBMC scRNA-seq samples for annotated ILC types is indicated in the annotation name. **(B)** Cell neighborhoods identified by differential abundance analysis between cells derived from ileum and PBMCs as shown in **A**, overlaid onto t-SNE coordinates also shown in **A**. Size of a circle indicates the number of cells in a neighborhood (Nhood size); color of a circle indicates magnitude of logFC in abundance in ileum (blue) versus PBMCs (red); width of lines between cell neighborhoods indicates the number of overlapping cells found in each of two neighborhoods (overlap size). **(C)** Pie chart of differential abundance results for cell neighborhoods shown in **B**. Grey indicates the proportion of cell neighborhoods that were not differentially abundant, while cell neighborhoods with significantly increased abundance (p<0.01) in ileum or PBMC samples are shown in blue and red, respectively. The logFC magnitude of differential abundance is also shown by red or blue shading. **(D)** Selected gene module detection scores from multidimensional differential gene expression analysis of cells shown in **A** overlaid onto two-dimensional t-SNE visualization coordinates. Color of a point corresponds to detection score for a gene module within a cell. **(E)** Violin plots summarizing gene module detections scores shown in **D** (y-axis) across annotated ILC types shown in **A** (x-axis). **(F)** Scatter plot of gene module 5 detection scores (y-axis) versus gene module 4 detection scores (x-axis) for all cells shown in **A**. Each point represents a single cell; color of a point corresponds to cell type annotations shown in **A**. **(G)** Scatter plot of gene module 3 detection scores (y-axis) versus gene module 7 detection scores (x-axis) for all cells shown in **A**. Each point represents a single cell; color of a point corresponds to cell type annotations shown in **A**. **(H)** Scatter plot of gene module 8 detection scores (y-axis) versus gene module 7 detection scores (x-axis) for all cells shown in **A**. Each point represents a single cell; color of a point corresponds to cell type annotations shown in **A**. scRNA-seq data shown in **A-H** were derived from ileum and PBMCs of two seven-week-old pigs. Ileum and PBMC samples for scRNA-seq were collected from the same two pigs and processed in parallel. Abbreviations: ILC (innate lymphoid cell); logFC (log fold-change); Nhood (neighborhood); No Sig (no significance); PBMC (peripheral blood mononuclear cell); scRNA-seq (single-cell RNA sequencing); t-SNE (t-distributed stochastic neighbor embedding)

Multidimensional differential gene expression analysis allowed comparison of ILC gene expression profiles independent of annotations assigned within and therefore confounded with ileum or PBMC datasets (**Supplementary Data 10**). Hierarchical clustering of differentially expressed genes with the lowest p-values defined nine gene modules with varying patterns of detection (**Figure 6D-E, Supplement to Figure 6-Figure 2A-C**). Gene module 4 had highest detection in peripheral ILCs, while module 5 had higher detection across all four types of ileal ILCs (**Figure 6D-E**), allowing us to easily segregate ILCs by peripheral or ileal origin based on gene module 4 versus 5 detection scores (**Figure 6F**). Gene module 7 had highest detection in ileal group 1 ILCs (activated, cytotoxic, cycling), while gene module 3 and 8 detection scores correlated positively with one another and were higher in ileal group 3 ILCs (**Figure 6D-E**; **Supplement to Figure 6-Figure 2D**). Collectively, the differences noted allowed for segregation between ileal group 1 and group 3 ILCs on the basis of detection for gene module 7 and 3/8, respectively (**Figure 6G-H**). Therefore, gene module 4 was considered as highly specific to peripheral ILCs, gene module 5 to all ileal ILCs, gene module 7 to ileal group 1 ILCs, and gene modules 3 and 8 to ileal group 3 ILCs. Further refinement focused on genes with the highest specificity for respectively-assigned ILC subsets from gene modules in **Figure 6D-E**, yielding core gene signatures for peripheral ILCs, all ileal ILCs, ileal group 1 ILCs, and ileal group 3 ILCs (**Figure 7A-D**).

**Figure 7.**
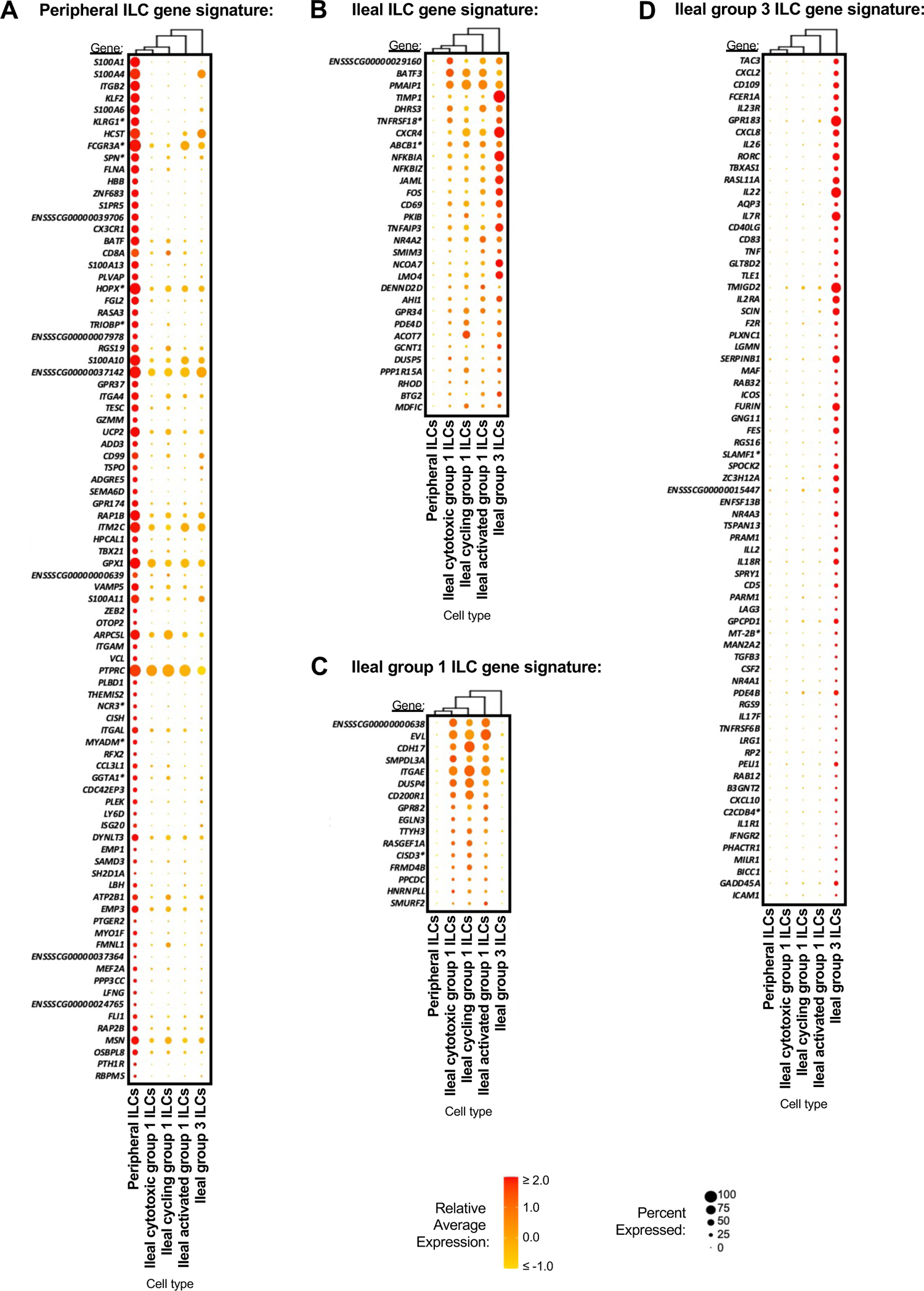
Core gene signatures of ileal and peripheral ILCs. **(A)** Hierarchical relationship of annotated ILC types from porcine ileum and PBMCs shown in a dendrogram (left) and dot plot showing expression patterns within each annotated ILC type for selected genes from gene module 4 used to create a peripheral ILC gene signature (right). In the dot plot, selected genes are listed on the x-axis, and ILC types are listed on the y-axis. Within the dot plot, size of a dot corresponds to the percentage of cells expressing a gene within an annotated cell type; color of a dot corresponds to average expression level of a gene for those cells expressing it within a cell type relative to all other cells in the dataset shown in **A**. **(B)** Hierarchical relationship of annotated ILC types from porcine ileum and PBMCs shown in a dendrogram (left) and dot plot showing expression patterns within each annotated ILC type for selected genes from gene module 5 used to create an ileal ILC gene signature (right). In the dot plot, selected genes are listed on the x-axis, and ILC types are listed on the y-axis. Within the dot plot, size of a dot corresponds to the percentage of cells expressing a gene within an annotated cell type; color of a dot corresponds to average expression level of a gene for those cells expressing it within a cell type relative to all other cells in the dataset shown in **A**. **(C)** Hierarchical relationship of annotated ILC types from porcine ileum and PBMCs shown in a dendrogram (left) and dot plot showing expression patterns within each annotated ILC type for selected genes from gene module 7 used to create an ileal group 1 ILC gene signature (right). In the dot plot, selected genes are listed on the x-axis, and ILC types are listed on the y-axis. Within the dot plot, size of a dot corresponds to the percentage of cells expressing a gene within an annotated cell type; color of a dot corresponds to average expression level of a gene for those cells expressing it within a cell type relative to all other cells in the dataset shown in **A**. **(D)** Hierarchical relationship of annotated ILC types from porcine ileum and PBMCs shown in a dendrogram (left) and dot plot showing expression patterns within each annotated ILC type for selected genes from gene modules 3 and 8 used to create an ileal group 3 ILC gene signature (right). In the dot plot, selected genes are listed on the x-axis, and ILC types are listed on the y-axis. Within the dot plot, size of a dot corresponds to the percentage of cells expressing a gene within an annotated cell type; color of a dot corresponds to average expression level of a gene for those cells expressing it within a cell type relative to all other cells in the dataset shown in **A**. scRNA-seq data shown in **A-D** were derived from ileum and PBMCs of two seven-week-old pigs. Ileum and PBMC samples for scRNA-seq were collected from the same two pigs and processed in parallel. *Ensembl identifiers found in gene annotation were converted to gene symbols; refer to methods section ‘*Gene name modifications*’ for more details Abbreviations: ILC (innate lymphoid cell); PBMC (peripheral blood mononuclear cell); scRNA-seq (single-cell RNA sequencing)

An 86-gene signature derived from gene module 4 was identified for peripheral ILCs (**Figure 7A**) and included several genes recognized as canonical markers for porcine NK cells (*CD8A*, *FCGR3A*, ITGAM, NCR3*, HCST*; Denyer et al., 2006; Gerner et al., 2009; Mair et al., 2013; Piriou-Guzylack and Salmon, 2008; Toka Felix et al., 2009) or recognized as canonical NK cell markers across other species (*KLRG1*, CD99, TBX21, GZMM*; Crinier et al., 2018; Huntington et al., 2007; Knox et al., 2014; Robinette et al., 2015; Townsend et al., 2004).

*FCGR3A\** and *CD8A* encode for CD16 and CD8α, respectively, which are the two primary markers used to identify porcine NK cells at the protein level (reviewed by Gerner et al., 2009; Piriou-Guzylack and Salmon, 2008). Thus, inclusion of *FCGR3A\** and *CD8A* in a gene signature for peripheral ILCs concordant with NK cell descriptions suggest current porcine NK cell identifiers are insufficient for pan-ILC identification in pigs. Additional peripheral ILC signature genes encoded for integrins (*ITGB2, ITGA4, ITGAM, ITGAL*) and other molecules associated with cell receptors/signaling (*FGR, PTPRC, CX3CR1, S1PR5*), consistent with enrichment of biological processes including integrin-mediated signaling pathway (*GO:0007229*), leukocyte cell-cell adhesion (*GO:0007159*), and receptor clustering (*GO:0043113*) in gene module 4.

Thirty genes derived from gene module 5 comprised an ileal ILC gene signature identifying both group 1 and group 3 ILCs in porcine ileum (**Figure 7B**). The gene signature for ileal ILCs included *CD69*, encoding for the cell surface receptor CD69 to identify tissue resident and recently activated cells (Ashouri and Weiss, 2017; Shiow et al., 2006; Szabo et al., 2019b). Enriched biological pathways for module 5 included those associated with cell metabolism, including positive regulation of macromolecule metabolic process (*GO:0010604*), positive regulation of cellular protein catabolic process (*GO:1903364*), and long-chain fatty acid metabolic process (*GO:0001676*), suggesting ileal ILCs were in a highly active metabolic state.

A gene signature for ileal group 1 ILCs included 16 genes derived from gene module 7, as shown in **Figure 7C**. Similar to gene module 5, gene module 7 was enriched for metabolic processes, including catabolic processes nucleoside phosphate catabolic process (*GO:1901292*) and biosynthetic processes such as purine-containing compound biosynthetic process (*GO:0072522*), suggesting an active metabolic state defining group 1 ILCs in ileum. Notably, *ITGAE*, which we used to identify intraepithelial group 1 ILCs in **Figure 5D**, was found in the ileal group 1 ILC gene signature, further supporting its use as an effective group 1 ILC marker in conjunction with other key markers in porcine ileum.

Group 3 ILCs had a core gene signature of 71 genes derived from gene modules 3 and 8 (**Figure 7D**). Many of the signature genes were associated with a type 3 immune response and largely overlapped with differentially expressed genes identified for ileal group 3 ILCs relative to all other ileal T/ILCs and relative to all other ileal ILCs (**Supplementary Data 4, 8**).

Similarly, enriched biological processes for gene modules 3/8 also had high overlap with enriched biological processes found from differentially expressed genes in **Supplementary Data 4, 8**. Notably, *IL22*, used as an *in situ* marker for group 3 ILCs in **Figure 5E**, was detected in the ileal group 3 ILC gene signature, supporting its use as an effective group 3 ILC marker in conjunction with other key markers, not only relative to other cells in ileum but also relative to peripheral ILCs.

## DISCUSSION

Understanding intestinal lymphocyte identity and function to cumulatively promote intestinal health outcomes in pigs has implications for biomedical research, animal health, and global food security. Phenotyping of porcine lymphocytes based on protein expression of a handful of cell surface markers has provided limited information about the biological dynamics of intestinal lymphocytes. Our work provides greater biological insight through further characterization of ileal lymphocytes into inferred functional subsets at transcriptional resolution not previously achieved in pigs, and we also include cross-species/cross-tissue comparisons and spatial context of specific lymphocyte types in different regions of the ileum. In addition to discoveries described herein, the data serves as a resource to be further explored and dissected for information about the heterogeneous landscape of lymphocytes in the porcine ileum.

Pigs readily serve as a biomedical model, with several innovations already applied to study intestinal lymphocytes. For example, germ-free pigs are used to study the dependence of intestinal lymphocyte development on microbial colonization (reviewed by Butler et al., 2009; Pabst, 2020; Sinkora and Butler, 2009, 2016). Germ-free pigs can be colonized with human microbiomes to generate microbially-humanized pigs for understanding the impact of the microbiota on local cellular responses associated with mucosal vaccination strategies and subsequent protection (Kumar et al., 2018; Miyazaki et al., 2018). Moreover, pigs serve as models for studying the impact of stress, nutrition, and infectious disease on intestinal physiology and immune status (reiviewed by Gonzalez et al., 2015; Hryhorowicz et al., 2020; Käser, 2021; Moeser et al., 2017; Roura et al., 2016; Ziegler et al., 2016). Bolstering lymphocyte-mediated intestinal immunity is important for both humans and pigs, but mechanistic approaches rely heavily on first understanding key lymphocytes in the porcine intestine and recognizing shared annotation with humans. To help overcome these obstacles, we identified and described biological implications of lymphocytes in the porcine ileum in comparison to both human and murine ileum. Collectively, our data indicate a general consensus of annotated lymphocyte types across species; however, a few notable differences were described. Therefore, our cross-species comparisons serve as a foundation for deeper-resolved exploration of pigs as models for human medicine. We further explore parallels between pigs and humans as it relates to intestinal lymphocytes throughout the rest of the discussion.

A caveat of scRNA-seq is loss of cellular spatial context when tissues are dissociated, which we partially overcame by differentially dissecting ileum into sections with or without Peyer’s patches prior to cell isolation. Cycling T/ILCs, follicular CD4 αβ T cells, B cells, and ASCs were enriched in Peyer’s patches, suggesting induction of germinal center antibody responses characteristic of GALT. Data support findings from human research, showing an enrichment of cycling cells, CD4 αβ T cells, and B cells in Peyer’s patches (Fujihashi et al., 2013; Junker et al., 2009) and thus strengthening the rationale for applicability of pigs to study Peyer’s patch-associated immune induction in biomedical research. Non-cycling γδ T, CD8 αβ T, and group 1 ILCs were enriched in the absence of Peyer’s patches and detected *in situ* primarily within the ileal epithelium, suggesting these three cell types mostly comprise the intraepithelial lymphocyte (IEL) community in porcine ileum, similar to humans (reviewed by Lutter et al., 2018; Mayassi and Jabri, 2018; Olivares-Villagómez and Van Kaer, 2018). Though porcine intraepithelial group 1 ILCs have not been studied in detail, we have conducted research investigating compositional changes of intraepithelial T lymphocytes (T-IELs) in the porcine intestine after weaning (Wiarda et al., 2020). Cross-species parallels were noted in our previous work, including a predominance of CD8^+^ αβ over γδ T-IELs in small intestine of pigs and humans (Jarry et al., 1990; Lundqvist et al., 1995; Olivares-Villagómez and Van Kaer, 2018; Wiarda et al., 2020). In contrast, γδ T-IELs are more frequent than CD8^+^ αβ T-IELs in murine small intestine (Guy-Grand et al., 1991; Hoytema van Konijnenburg et al., 2017). Thus, our previous work gives initial support of pigs for IEL-centric biomedical research, and scRNA-seq of γδ T, CD8 αβ T, and group 1 ILCs presented herein may help to build on this application. We caution that lamina propria versus epithelial location of single cells cannot be definitively concluded solely from our scRNA-seq data. However, T cells studied via scRNA-seq from lamina propria and epithelium of human ileum had highly overlapping transcriptional profiles (Jaeger et al., 2021), and the same may be true for pigs.

Non-naïve T and B cells in porcine ileum were transcriptionally different from comparable cells in blood, as indicated by lower average mapping scores to porcine PBMCs. Transcriptional differences between ileal- and blood-derived non-naïve T/B cells likely occurred because of inherently activated and/or differentiating states of ileal lymphocytes due to exposure to luminal contents, including the microbiota and microbial-derived metabolites. Our supposition is reinforced by work in germ-free pigs, where small intestinal non-naive T and B cell abundances are reduced in the absence of microbial exposure (Barman et al., 1996; Haverson et al., 2007; Potockova et al., 2015; Rothkötter et al., 1994; Rothkötter and Pabst, 1989; Sinkora et al., 2011). Differences in T/B cells from mucosal tissues versus blood are also observed in humans, where non-naïve lymphocytes increase in abundance, and tissue-specific gene signatures for lymphocyte activation, tissue residency, and/or effector functions are observed for tonsillar B cells (Glass et al., 2020; King et al., 2021), lung-derived T cells (Szabo et al., 2019a), and intestinal T cells (Uniken Venema et al., 2019) compared to similar lymphocyte subsets in the periphery. Collectively, the aforementioned studies and our own work suggest mucosal sites (including the ileum) are locations for congregation of non-naïve T/B cells transcriptionally distinct from T/B cells found in the periphery, likely due in large part to ongoing immune stimulation via microbial exposure and other context-dependent signals at mucosal surfaces.

Similar to intestinal T and B cells, ILCs in the porcine intestine are likely heavily influenced by the microbiota; however, little is known about the influence of microbial colonization on porcine intestinal ILCs due to lack of methods to clearly label and identify intestinal ILCs in pigs. We established gene expression profiles and locational context of group 1 and group 3 ILCs in porcine ileum to cautiously make functional inferences about ILCs in the porcine intestine. Our findings support comparable roles of porcine ileal ILC functions to intestinal ILCs in humans and mice, such as epithelial patrolling behaviors of intraepithelial group 1 ILCs (Fuchs et al., 2013; Talayero et al., 2016; Van Acker et al., 2017) and contribution of lamina propria and GALT-associated group 3 ILCs in immune defense, regulation of the commensal microbiota, GALT development, tissue homeostasis, and antibody production (Aparicio-Domingo et al., 2015; Guo et al., 2014; Kruglov Andrey et al., 2013; Mortha et al., 2014; Satoh-Takayama et al., 2008; Sonnenberg Gregory et al., 2012; Tsuji et al., 2008; von Burg et al., 2014). In mice, intestinal ILC1s proportionally contract and convert to a less polarized ILC3-like regulatory profile in germ-free compared to specific pathogen-free animals, suggesting ILC1 recruitment, differentiation, and/or function may be dampened without microbial stimulation (Gury-BenAri et al., 2016), and it remains plausible that similar phenomena occur in pigs. In support, porcine ileal ILCs gene signatures appeared to be heavily influenced by tissue-specific cell activation when compared to peripheral ILCs, and microbiota-derived signals likely supply tissue-specific transcriptional imprinting. In humans, ILCs from tonsil, lung, and colon have tissue-specific gene expression indicative of tissue residency, cell activation, and modified metabolism compared to peripheral ILCs (Mazzurana et al., 2021), again supporting the contribution of external stimuli in driving activation state. Therefore, our data and newly developed methods for ILC detection should prove useful for further investigation, including defining the impact of the microbiota or intestinal infection on intestinal ILC function in pigs. We could not identify ileal group 2 ILCs in porcine ileum; however, it is possible that group 2 ILCs were not found due to low abundances occurring under steady state conditions. In humans, intestinal group 2 ILCs are similarly rare under steady state conditions but may increase in abundance under conditions that facilitate their recruitment and/or expansion in the intestine, such as with parasitic infection of cancer (Meininger et al., 2020; Qi et al., 2021), and it remains possible that similar phenomena may occur in pigs. In contrast, group 2 ILCs comprise approximately one-quarter of ILCs in the lamina propria of the murine small intestine (Dutton et al., 2017). Thus, predominance of group 1 and group 3 ILCs in the pig ileum provides initial indication that pigs may have intestinal ILC populations more similar in composition to humans than that of mice under steady state.

Direct comparison of porcine ileal ILCs and peripheral NK cells brings into question the current methods for identification of non-T/non-B lymphocytes in pigs, as key markers used to identify NK cells (the only non-T/non-B lymphocyte subset currently identified in pigs) were found in gene signatures for peripheral but not ileal porcine ILCs. Ileal group 3 ILCs were easily distinguishable from peripheral NK cells, while ileal group 1 ILCs shared more similarities to peripheral NK cells. It remains undetermined whether ileal group 1 ILCs are ILC1s or NK cells, as detected differences between ileal group 1 ILCs and circulating NK cells in our data could indicate (1) identification of ILC1s present in the ileum but largely absent from periphery, (2) tissue-specific differences in ileal-derived NK cells that do not fit current phenotypic descriptions used to identify NK cells in pigs, or (3) a combination of both. Unfortunately, differentiation between ILC1s and NK cells remains complicated even in better-characterized humans and mice due to species-to-species and tissue-to-tissue peculiarities within what can still be considered relatively recently discovered immune cell subsets (Meininger et al., 2020; Robinette et al., 2015; Simoni et al., 2017; Simoni and Newell, 2017; Van Acker et al., 2017).

Regardless, we stress the importance of focusing on cell function over technical phenotype classification, as the functional role of cells in the context of immune protection and immunopathology is ultimately what contributes to biological outcome. To this point, gene signatures obtained for ileal and peripheral ILCs may further be used for identifying biomarkers for detection of specific ILC subsets and understanding their biological functions.

To our knowledge, our findings encompass the first single-cell descriptions of the immune cell transcriptomic landscape in porcine intestine, with primary focus on the study of lymphocytes in the ileum, including T cells, ILCs, B cells, and ASCs. In addition, non-lymphocyte cell types in the intestine play important functional roles in intestinal health but fell outside the scope of the current study. The diverse spectrum of biological states for cells captured via scRNA-seq is difficult to holistically describe, and we only scratch the surface of the biological information and complexities contained within our scRNA-seq data. While our data may serve as a starting point for understanding the roles of specific immune cells in a specific biological scenario, such understandings are best fine-tuned using the most appropriate species, tissue, and biological scenario of interest. Consequently, our data serve as an exciting starting point for query to seed future research questions.

## MATERIALS AND METHODS

### Animals and sample collection

Mixed-breed and mixed-gender pigs were obtained from commercial nursery settings for all experiments and assays. All pigs were weaned at approximately three weeks of age. For scRNA-seq, samples were collected from one female (pig 1) and one male (pig 2) at approximately seven weeks of age. Post-mortem intestinal tissues collected to complete *ex vivo* and *in situ* assays were from control animals in unrelated studies to reduce the overall number of animals used. The ages of animals are listed within respective figure captions and ranged from approximately five to nine weeks of age. Humane euthanasia was carried out immediately preceding tissue collections according to procedures approved by the Institutional Animal Care and Use Committee (IACUC) at Iowa State University or the National Animal Disease Center, Agricultural Research Service (ARS), United States Department of Agriculture (USDA).

### Cell isolations

#### Ileum for scRNA-seq

For ileal cell isolations, reagents were equilibrated to room temperature (RT), and samples were stored at RT between steps. Immediately following humane euthanasia, ileal tissue was collected and stored in stabilization buffer (2 mM ethylenediaminetetraacetic acid [EDTA; Invitrogen AM9260G], 2mM L-glutamine [Gibco 25-030], and 0.5% bovine serum albumin [BSA; Sigma-Aldrich A9418] in Hank’s balanced salt solution [HBSS; Gibco 14175]) for transport back to the lab. In the lab, the exterior muscularis layer was peeled off, and tissues were cut open longitudinally to expose the lumen. Tissues were gently rinsed with phosphate buffered saline (PBS), pH 7.2 (made in-house) to remove intestinal contents and carefully dissected into regions of interest (PP, non-PP, whole ileum), as shown in **Supplement to Figure 1-Figure 1B**. Dissected tissues were weighed out to between 1 to 1.5 grams per sample for further use.

Tissues were sequentially transferred to and incubated in the following solutions in a shaking incubator at 37 °C, 200 rpm: 20 minutes in 30 mL mucus dissociation solution (5 mM dithiothretol [DTT; Invitrogen 15508] and 2% heat-inactivated fetal calf serum [FCS; Gibco A38401] in HBSS); 25 minutes in 30 mL epithelial removal solution (5 mM EDTA and 2% FCS in HBSS), repeated with fresh solution for a total of three times; and 10 minutes in 20 mL wash solution (10 mM HEPES [Fisher BioReagents BP299] in HBSS). Epithelial removal and wash solutions were retained for processing of epithelial cells, while mucus dissociation solution was discarded.

Following incubation in wash solution, tissues were minced and placed into gentleMACS C tubes (Miltenyi 130-093-237) containing 15 mL enzyme digestion solution (10 mM HEPES, 0.2 U/mL Liberase TM [Roche 5401127001], and 30 μg/mL DNase I [Sigma D5025] in HBSS). Tissue dissociation was carried out using the intestine C-tube protocol on a gentleMACS Octo Dissociator (Miltenyi 130-095-937) followed by incubation at 37 °C, 200 rpm for 45 minutes and another round of mechanical dissociation using the gentleMACs intestine C-tube protocol. 10 mL stabilization buffer was added to each C-tube, and contents were strained through sterile non-woven surgical gauze sponges (Starryshine GZNW44), followed by a 100-micron nylon mesh cell strainer (BD Falcon 352360).

While tissues were incubating in enzyme digestion solution, epithelial cells were collected by passing epithelial removal and wash solutions through a 100-micron nylon mesh cell strainer, centrifuging at 450 xg for 8 minutes RT, and resuspending in supplemented HBSS (2mM L-glutamine and 2% FCS in HBSS).

Cell fractions from epithelial isolation and tissue digestion for each sample were combined, centrifuged at 450 xg for 8 minutes RT, and resuspended in 24 mL of RT 70% Percoll^TM^ (1.088 g/mL at 22 °C; GE Healthcare Life Sciences 17-0891-01). Aliquots of 8 mL cell/Percoll^TM^ suspension were overlayed with 4 mL HBSS and centrifuged at 400 xg for 30 minutes RT, with slow acceleration and centrifuge break turned off. The density interphase layer of cells was collected, washed with supplemented HBSS, centrifuged at 450 xg for 8 minutes RT, and resuspended again in supplemented HBSS. Quantity and viability of cells was assessed by the Muse Count & Viability Assay Kit (Luminex MCH100102) using a Muse Cell Analyzer (Luminex 0500-3115).

To further enrich for live cells, cell suspensions were passed through another 100-micron nylon filter, centrifuged at 300 xg for 10 minutes RT, and processed using a Dead Cell Removal Kit (Miltenyi 130-090-101). Cells were resuspended in 100 μL kit microbeads per 10^7^ total cells, incubated for 15 minutes, and divided into four equal parts per sample that were each rinsed with 1 mL 1X Binding Buffer. A total of four separate LS columns (Miltenyi 130-042-401) were used for the divided samples to facilitate magnetic sorting using a MultiMACS^TM^ Cell24 Separator Plus (Miltenyi 130-098-637). LS columns were prerinsed with 1X Binding Buffer before applying cells. Columns were rinsed with an additional 3 mL of 1X Binding Buffer twice to facilitate cell pass through. The negative cell pass through was collected, centrifuged 300 xg for 10 minutes RT, resuspended in supplemented HBSS, centrifuged again, and resuspended in a final volume of supplemented HBSS. Muse count and viability was again assessed and deemed adequate for scRNA-seq (>84% live cells per sample). Samples were transported to the sequencing facility (∼15 minute transport), and cell viability was reassessed using a Countess II Automated Cell Counter (ThermoFisher Scientific). Viabilities from the Countess readings were deemed adequate to proceed with partitioning (>76% live cells per sample) for scRNA-seq.

#### PBMCs for scRNA-seq

PBMCs were isolated using Cell Preparation Tubes (CPT^TM^; BD Biosciences 362782) according to manufacturer’s recommendations.

#### Ileum for flow cytometry

Ileal cells collected for flow cytometry experiments were isolated as described for scRNA-seq ileal cells above, with the exception that further enrichment of viable lymphocytes by Percoll^TM^ density gradient centrifugation and magnetic dead cell removal were not performed.

#### Droplet-based scRNA-seq

Single-cell suspensions from ileum and PBMCs were prepared for scRNA-seq by partitioning and preparing libraries for a target of 10,000 cells per sample according to the manufacturer’s protocol for Chromium Single Cell 3’ v2 Chemistry (10X Genomics CG00052). Samples were multiplexed, had equal proportions of cDNA from each sample pooled, and were run across multiple lanes of an Illumina HiSeq 3000 with 2×100 paired-end sequencing as previously described (Herrera-Uribe & Wiarda et al., 2021). Raw data were deposited in .fastq file format for both forward and reverse strands following image analysis, base calling, and demultiplexing.

#### scRNA-seq analysis of ileum data Initial data processing

Initial processing of scRNA-seq data included read alignment/gene quantification, ambient RNA removal, gene/cell filtering, doublet removal, normalization, integration, and dimensionality reduction identical to as previously described (Herrera-Uribe & Wiarda et al., 2021) and as briefly outlined below.

Read alignment, mapping, and gene quantification were carried out with the shell data package, CellRanger v4.0.0 (10X Genomics) and the *Sus scrofa* 11.1 reference genome with 11.1.97 annotation file obtained from Ensembl (Cunningham et al., 2019) and modified as previously described (Herrera-Uribe & Wiarda et al., 2021). Ambient RNA removal was performed using the auto-estimation method from the R package, SoupX v1.4.5 (Young and Behjati, 2020).

Genes without any reads detected in the total of all sequencing reads across all cells and samples were removed from the dataset. Cells with >12.5% of total reads attributed to mitochondrial genes, <550 total genes detected, or <1,250 total unique molecular identifiers (UMIs) detected were removed from the dataset (**Supplementary Data 1**).

The Python package, Scrublet v0.1 (Wolock et al., 2019), was used to remove highly-probable neotypic doublets from the remaining dataset using a doublet rate of 0.07. Cells with corresponding doublet probability scores >0.25 were removed from our dataset (**Supplementary Data 1**).

Normalization, integration, and dimensionality reduction were performed using the R package, Seurat v3.2.2 (Butler et al., 2018; Stuart et al., 2019). SCT-normalized data was utilized to perform anchor-based sample integration with default parameters. Principle component analysis (PCA) was performed to identify the first 100 principle components (PCs) of the data, and a ‘significant’ number of PCs to use for further analyses was determined as the smaller value of (1) the highest PC that had >0.1% change in variation between consecutive PCs or (2) the smallest PC that represented >90% of the cumulative variation and <5% of variation associated with a single PC. t-distributed stochastic neighbor embedding (t-SNE) coordinates were generated for visualization using the significant number of PCs. Log-normalized and scaled data were also calculated for the RNA assay of the Seurat object.

Throughout the workflow, intermediate modified count matrices were generated and converted back to 10X format using the function *write10XCounts()* from the R package, DropletUtils v1.8.0 (Griffiths et al., 2018; Lun et al., 2019).

#### Quality check of ileal sample types

Disparities in the percentage of cells removed between different ileal sample types were noted in **Supplementary Data 1**, with smaller percentages of cells passing all cell filtering steps from non-PP (47.24% and 54.21%) compared to PP (75.31% and 77.13%) and whole ileum (71.56% and 77.40%) samples. Further investigation was performed to ensure disparities were due to differences in sample cell compositions and not differences in the quality of the same cell types between samples. Our query revealed many poor quality cells (>12.5% mitochondrial reads, <550 genes, <1,250 UMIs) that were more abundant in non-PP samples expressed genes characteristic of epithelial cells (*EPCAM, KRT8*) rather than immune cells (*PTPRC* [encoding pan-leukocyte marker CD45]; **Supplement to Figure 1-Figure 2A**). Results were further validated by assessing the ratio of leukocytes (CD45^+^) to epithelial cells (EPCAM^+^) by flow cytometry (see flow cytometry methods) in independently collected ileal samples (**Supplement to Figure 1-Figure 2B-C**). Non-PP samples had smaller leukocyte:epithelial cell ratios than did PP or whole ileum samples (**Supplement to Figure 1-Figure 2D**), indicating a higher occurrence of epithelial cells in non-PP samples. Similar observations were made via IHC, showing that a larger proportion of epithelial cells (stained by pan-cytokeratin protein expression; see IHC methods) were present in regions of ileum lacking Peyer’s patches (**Supplement to Figure 1-Figure 2E**). Thus, smaller percentages of cells passing all cell filtering steps from non-PP samples were largely attributed to a higher occurrence of poor quality epithelial cells originally present in these samples.

#### Cell clustering

The Seurat function *FindNeighbors()* was used to construct a shared nearest neighbor (SNN) graph, specifying to use the significant number of PCs calculated during data dimensionality reduction outlined above. The Seurat function *FindClusters()* using the significant number of PCs was used to identify clusters with clustering resolutions at 0.5 intervals between 0 and 5. Clustering at different resolutions was compared using the function *clustree()* from the R package, clustree v0.4.3 (Zappia and Oshlack, 2018). A clustering resolution of 3 was selected for all downstream work.

#### Hierarchical clustering

Hierarchical clustering was performed with the Seurat function *BuildClusterTree()*, specifying to use only the number of significant PCs calculated during data dimensionality reduction outlined above for a respective dataset. Clustering dendrograms were visualized using the function, *PlotClusterTree()*.

#### Generating data subsets

Some analyses were conducted on only a subset of cells from the original ileum scRNA-seq dataset. To partition out only cells of interest into smaller datasets, we used the Seurat function *subset()* to specify which cells to allocate into smaller datasets. Genes with cumulative zero expression in the new data subsets, scaled data, and dimensionality reduction dimensions were removed using the function *DietSeurat()*. Data in the RNA assay were re-scaled; the first 100 PCs were re-calculated; the number of significant PCs were re-determined; and t-SNE dimensions were re-calculated using methods described above. For some smaller data subsets, integration anchors could not be calculated with default parameter k.filter=200 for the function *FindIntegrationAnchors()*, and the k.filter parameter was adjusted to the largest multiple of 5 at which integration anchors could still be calculated for the dataset.

#### Cell cluster/annotation-based differential gene expression analysis

Differential gene expression (DGE) analyses were performed using functions of the Seurat package and normalized gene counts from the RNA assay. Differentially expressed genes (DEGs) were calculated for each cluster/cell type relative to the average gene expression across an entire dataset using *FindAllMarkers()*. To be considered differentially expressed (DE), a gene was expressed in >10% of cells in one of the populations being compared, had a |logFC| >0.25, and had a corrected p-value <0.05.

#### Multidimensional differential gene expression analysis

Cell cluster/type-independent, multidimensional DGE analysis was performed with the R package, singleCellHaystack v0.3.3 (Vandenbon and Diez, 2020). From log-normalized counts stored in the RNA assay of our Seurat object, median expression levels were calculated for each gene across the entire dataset, followed by determining if expression of each gene was above or below the median expression level within each cell. From this information, DGE was calculated using the function *haystack_highD()*, specifying to use the previously determined number of significant PCs to define the multidimensional space. A gene was considered DE if it had an adjusted p-value <0.05.

Gene modules were created by first selecting only differentially expressed genes with adjusted p-values <1×10^-10^, followed by hierarchical clustering of selected genes with the function *hclust_haystack_highD()*. The number of gene modules to split the gene dendrogram into (*k*) was specified between *k*=3 to *k*=10 and executed with the function *cutree()*. Detection scores of each gene module within each cell were calculated with the function *plot_gene_set_haystack()*. The final value of *k* was selected by examining each model and selecting the one producing the most interpretable and biologically-relevant gene modules.

#### Topic modeling

Topic models were fit with the fastTopics package v0.5-54 (Carbonetto et al., 2021; Dey et al., 2017). Multiple topic models were fit for each cell type subset (*K*=3 to *K*=10). The final value of *K* was selected by examining each model and selecting the one that produced the most interpretable and biologically relevant structure. Genes which were enriched in each topic were determined using the *diff_count_analysis()* function.

#### Biological process enrichment analysis

After identification of DEGs with increased expression in cell clusters, topics, or gene modules, gene sets were subjected to Gene Ontology (GO) term enrichment analysis. GO terms were mapped to Ensembl gene IDs using the biomaRt v2.48.3 (Durinck et al., 2009) R package. Only genes, and therefore GO terms, detected in our data were included in the background set which the enriched sets were compared against. A new GO term background set was created for each data subset. Genes and GO terms which were not detected in each data subset were removed from the respective background list. The R package topGO v2.44.0 (Alexa A and J, 2021; Alexa et al., 2006) was used to carry out GO term enrichment analysis using the ‘elimination’ algorithm and the ‘Fisher’ test statistic. GO terms with p-values <0.05 were considered enriched in the gene set in question relative to the appropriate background GO term set.

#### Ileal cell lineage annotations

Ileal cells were grouped into 54 transcriptionally-distinct clusters as described in *Cell clustering* methods (**Supplement to Figure 1-Figure 4A**). Query of cell lineage canonical gene expression within clusters was utilized to group cells into four major cell lineages (**Figure 1C**; **Supplement to Figure 1-Figure 4B-C**). Since cell clusters did not have highly overlapping expression of canonical genes characteristic of multiple cell lineages, all cells in a cluster were assigned a single cell lineage identity based on the gene expression profiles seen at the cluster level. By this rationale, 14,742 cells in 26 clusters (1, 3, 6, 7, 12, 14, 15, 17, 18, 19, 23, 24, 26, 29, 31, 32, 34, 35, 37, 41, 43, 44, 46, 47, 51, 53) were classified as T/ILC lineage lymphocytes and expressed genes such as *CD3E, CD3G, CD52,* and *ZAP70* involved in T cell receptor (TCR) signaling or ILC intracellular signaling (Bandala-Sanchez et al., 2013; Ginaldi et al., 1998; Straus and Weiss, 1993; Tufa et al., 2020). 16,070 cells in 22 cell clusters (0, 2, 4, 5, 8, 9, 10, 11, 13, 16, 20, 21, 22, 25, 27, 28, 30, 33, 38, 39, 40, 48) were classified as B lineage lymphocytes based on expression of genes such as *CD19, CD79B, MS4A1,* and *JCHAIN* associated with B cell receptor (BCR) signaling/antibody secretion (Herrera-Uribe & Wiarda et al., 2021; Lee et al., 2021). 458 cells in three clusters (42, 49, 52) were classified as myeloid lineage leukocytes and expressed genes such as *SIRPA*, CD68, CSF2RB,* and *ICAM1* that customarily have high expression in macrophages, DCs, and/or mast cells (Akula et al., 2020; Betjes et al., 1991; Valent et al., 1991). 713 cells in three clusters (36, 45, 50) were classified as non-leukocytes based on a lack of pan-leukocyte marker *PTPRC* expression and high expression of genes such as *EPCAM, KRT8, PECAM1,* and *COL18A1* commonly expressed by epithelial/stromal cells (Elmentaite et al., 2021; Han et al., 2018).

#### Ileal cell type annotations

Ileal cells were further annotated into 26 cell types (**Figure 1D**; **Supplement to Figure 1-Figure 3**) with a hybrid, multi-method approach utilizing (1) cell clustering and accompanying cluster-based DGE and hierarchical analyses (discrete cell/non-discrete gene classifications); (2) cell cluster-independent multidimensional DGE analyses and gene module detection (non-discrete cell/discrete gene classifications); and/or (3) grade of membership/topic modeling (non-discrete cell/non-discrete gene classifications) as described below.

#### T/ILC lineage lymphocyte annotations

Cells assigned as T/ILC lineage lymphocytes (**Figure 1C**; **Supplement to Figure 1-Figure 4B-C**) were extracted to form a subset of data for further query of T/ILC identities (as described in *Generating data subsets* methods; **Supplement to Figure 1-Figure 5A**). Twenty-one T cell clusters (3, 6, 7, 12, 14, 15, 17, 19, 23, 24, 26, 29, 31, 32, 34, 35, 37, 41, 46, 47, 51) were identified by expression of porcine pan-T cell marker *CD3E* (expressed by >75% of cells in a cluster; **Supplement to Figure 1-Figure 5B-C**; reviewed by Gerner et al., 2009; Piriou-Guzylack and Salmon, 2008). Meanwhile, the majority of cells in five clusters (1, 18, 43, 44, 53) did not express *CD3E* but, similar to porcine NK cells, did largely express *CD2*, leaving us to identify these cells more broadly as ILCs (**Supplement to Figure 1-Figure 5B-C**; Herrera-Uribe & Wiarda et al., 2021). Within the 21 T cell clusters, clusters were classified as CD4 αβ T cells, CD8 αβ T cells, or γδ T cells if >10% of cells in a cluster expressed *CD4, CD8B,* or *TRDC*, respectively (**Supplement to Figure 1-Figure 5D**). By these criteria, five clusters (12, 15, 26, 41, 46) were identified as CD4 αβ T cells; five clusters (3, 7, 14, 17, 37) were identified as CD8 αβ T cells; and γδ T cells were identified in six clusters (6, 23, 29, 31, 32, 51). Cluster 24 was identified as a mixture of CD4 and CD8 αβ T cells by expression of *CD4* and *CD8B* but not *TRDC* in >10% of cells. Four clusters (19, 34, 35, 47) expressed both *TRDC* and *CD8B* but not *CD4* in >10% of cells and were thus classified as a mixture of γδ and CD8 αβ T cells. Of note, mixed CD4/CD8 αβ and γδ/CD8 αβ T cell clusters appeared to be a mixture of cells expressing either *CD4, CD8B,* or *TRDC* rather than cells co-expressing any two of these markers (**Supplement to Figure 1-Figure 5B**). Thus, T/ILC lineage lymphocytes were further classified as CD4 αβ T cells, mixed CD4/CD8 αβ T cells, CD8 αβ T cells, mixed γδ/CD8 αβ T cells, γδ T cells, or ILCs (**Figure 1-Figure Supplement 5E**), and these classifications were used to generate additional T/ILC data subsets for further annotations.

#### Naïve αβ T cell annotation

Further cell cluster-based DGE analysis of all T/ILC lineage lymphocytes revealed a unique gene expression signature characteristic of circulating and/or naïve T cells in the only mixed CD4/CD8 αβ T cell cluster, cluster 24 (**Supplement to Figure 1-Figure 5F**). Thus, cluster 24 cells were given the designation of naïve CD4/CD8 αβ T cells. Gene expression profiles and GO enrichment for naïve CD4/CD8 αβ T cells relative to all other T/ILC lineage lymphocytes were discussed in results and are synonymous with those also recovered for cluster 24, thus results and rationale for our naïve T cell annotation can be found in **Supplementary Data 8** and are described in preceding results.

#### CD4 αβ T cell annotations

Cells identified as CD4 αβ T cells (clusters 12, 15, 26, 41, 46; **Supplement to Figure 1-Figure 5**) were extracted to create a subset of data (as described in *Generating data subsets* methods). From this data subset, previous cell cluster assignments (as described in *Cell clustering* methods), topic modeling with weighted membership to three topics (as described in *Topic modeling* methods), and multidimensional DGE analysis with detection of three gene modules (as described in *Multidimensional differential gene expression analysis* methods) were assessed to identify DEGs and associated enriched biological processes by each method (**Supplement to Figure 1-Figure 6A-D**; **Supplementary Data 2**).

Topic 3 had significantly greater expression of many genes associated with cellular replication/division, and similar genes were also found within gene module 2 (e.g. *PCLAF, BIRC5, TOP2A, UBE2C*; Dabydeen et al., 2019; Giotti et al., 2019), resulting in enrichment of biological processes such as mitotic sister chromatid segregation (*GO:0000070*), DNA replication (*GO:0006260*), and microtubule cytoskeleton organization involved in mitosis (*GO:1902850*) in both topic 3 and gene module 2 (**Supplementary Data 2**). Therefore, we identified cycling CD4 αβ T cells as CD4 αβ T cells with high topic 3 weights (>0.4) and/or high gene module 2 detection scores (>0.1; **Supplement to Figure 1-Figure 6E**). Topic 2/gene module 3 were both enriched for biological processes including mature B cell differentiation (*GO:0002335*) and positive regulation of humoral immune responses mediated by circulating immunoglobulin (*GO:0002925*) due to contribution from genes with significantly greater expression in both topic 2/gene module 3 that were characteristic of follicle-associated helper T cells, including *CXCR5, PDCD1,* and *IL6R* (**Supplementary Data 2**; Chtanova et al., 2004; Haynes et al., 2007; Schaerli et al., 2000; Shi et al., 2018). Conversely, topic 1 had significantly increased expression of genes indicative of cellular activation and effector functions not necessarily related to follicular T cell help, including *FCGR3A*, IFI6, CCL5* (Szabo et al., 2019a), and many ribosomal genes, and enrichment of biological processes including translation (*GO:0006412*), positive regulation of interferon-gamma production (*GO:0032729*), and T cell differentiation (*GO:0030217*; **Supplementary Data 2**). Consequently, follicular CD4 αβ T cells were identified from remaining non-cycling CD4 αβ T cells as those with higher topic 2 than topic 1 weights and/or high gene module 3 detection scores (>0.3; **Supplement to Figure 1-Figure 6F**). Remaining cells that did not fit criteria for cycling or follicular CD4 αβ T cells were identified as activated CD4 αβ T cells, as they still had high topic 1 weighting (**Supplement to Figure 1-Figure 6F**). In total, we annotated three populations of CD4 αβ τ cells, including cycling CD4 αβ T cells, follicular CD4 αβ T cells, and activated CD4 αβ T cells (**Supplement to Figure 1-Figure 6G-H**).

##### γδ/CD8 αβ T cell annotations

Cells identified as γδ T cells, CD8 αβ T cells, or a mixture of γδ/CD8 αβ T cells (clusters 3, 6, 7, 14, 17, 19, 23, 29, 31, 32, 34, 35, 37, 47, 51; **Supplement to Figure 1-Figure 5**) were extracted to create a subset of data (as described in *Generating data subsets* methods). From this data subset, previous cell cluster assignments (as described in *Cell clustering* methods), topic modeling with weighted membership to three topics (as described in *Topic modeling* methods), and multidimensional DGE analysis with detection of four gene modules (as described in *Multidimensional differential gene expression analysis* methods) were assessed to identify DEGs and associated enriched biological processes by each method (**Supplement to Figure 1-Figure 7A-D**; **Supplementary Data 3**).

Cell cluster-based DGE analysis and hierarchical clustering revealed cluster 51, previously identified as a cluster of γδ T cells (**Supplement to Figure 1-Figure 5**), to be most distantly related from other γδ/CD8 αβ T cell clusters and to have significantly higher and nearly ubiquitous expression of *SELL* (encoding CD62L), which was not expressed at high levels in other γδ/CD8 αβ T cells (**Supplement to Figure 1-Figure 7B, Supplementary Data 3**). Therefore, cells with cluster 51 membership were annotated as *SELL*^hi^ γδ T cells. Another cluster of γδ T cells, cluster 31, had a unique transcriptional profile where cells largely lacked expression of *CD2* (**Supplement to Figure 1-Figure 5B-C**). Therefore, cells in cluster 31 were annotated as CD2^-^ γδ T cells.

Both topic 3 and gene module 3 had significantly increased expression of genes related to cellular replication and/or division, such as *PCLAF, BIRC5, TOP2A, UBE2C* (Dabydeen et al., 2019; Giotti et al., 2019), and were both enriched for biological processes such as regulation of mitotic cell cycle (*GO:0007346*), mitotic sister chromatid separation (*GO:0000070*), and cell cycle DNA replication (*GO:0044786*; **Supplementary Data 3**). From the remaining γδ/CD8 αβ T cells (excluding *SELL*^hi^ γδ T cells and CD2^-^ γδ T cells), we identified cells with high weighting and/or high detection scores for topic 3 (>0.41) and gene module 3 (>0.11), respectively, as cycling cells (**Supplement to Figure 1-Figure 7E**). Topic 1 and gene module 1 were both enriched for genes characteristic of cell activation, such as *CTSW, CCR9, SLA-DQB1, XCL1,* and many ribosomal genes (Boismenu et al., 1996; Gerner et al., 2009; Iwata et al., 2004; Kelner Gregory et al., 1994; Ondr and Pham, 2004; Stoeckle et al., 2009; Svensson et al., 2002; Uehara et al., 2002), and biological processes including antigen processing and presentation of exogenous peptide antigen via MHC class II (*GO:0019886*), ribosomal large subunit assembly (*GO:0000027*), and immune effector process (*GO:0002252*; **Supplementary Data 3**).

Meanwhile, topic 2 had significantly increased expression of genes related to cytotoxic effector functions, such as *GZMA*, GZMB,* and GNLY (Hidalgo et al., 2008), and biological processes including natural killer cell activation (*GO:0030101*), natural killer cell differentiation (*GO:0001779*), and positive regulation of natural killer cell mediated cytotoxicity (*GO:0045954*; **Supplementary Data 3**). Therefore, of the remaining cells (excluding *SELL*^hi^ γδ T cells, CD2^-^ γδ T cells, and cycling cells), cells with high topic 1 weights (topic 1 weight at least 4x greater than topic 2 weight) and/or high gene module 1 detection scores (>0.25) were called activated T cells, while remaining cells were classified as cytotoxic T cells due to high topic 2 weighting (**Supplement to Figure 1-Figure 7F**).

Furthermore, cycling, cytotoxic, and activated T cells were classified as γδ or CD8 αβ T cells by assessing expression ratios of *TRDC:CD8B*. Cells with a ratio >1 were classified as γδ T cells, while remaining cells with a ratio <1 were classified as CD8 αβ T cells. Cells in which both *TRDC* and *CD8B* expression were zero were classified as CD8 αβ T cells, as *TRDC* gene expression was observed to be less sparse than *CD8B* expression, decreasing the likelihood of a technical zero being detected for *TRDC* over *CD8B* in our sparse data matrix (**Supplement to Figure 1-Figure 7G-H**). Collectively, we annotated eight populations of γδ/CD8 αβ T cells, including *SELL*^hi^ γδ T cells, CD2^-^ γδ T cells, cycling γδ T cells, activated γδ T cells, cytotoxic γδ T cells, cycling CD8 αβ T cells, activated CD8 αβ T cells, and cytotoxic CD8 αβ T cells (**Supplement to Figure 1-Figure 7I-J**).

#### ILC annotations

Cells identified as ILCs (clusters 1, 18, 43, 44, 51; **Supplement to Figure 1-Figure 5**) were extracted to create a subset of data (as described in *Generating data subsets* methods). From this data subset, previous cell cluster assignments (as described in *Cell clustering* methods), topic modeling with weighted membership to three topics (as described in *Topic modeling* methods), and multidimensional DGE analysis with detection of three gene modules (as described in *Multidimensional differential gene expression analysis* methods) were assessed to identify DEGs and associated enriched biological processes by each method (**Supplement to Figure 1-Figure 8A-D**; **Supplementary Data 4**).

We noted cluster 43 to be the most distantly-related from other ILC clusters (**Supplement to Figure 1-Figure 8B**) and also had high topic 1 weights, low gene module 3 detection, and high gene module 1 detection (**Supplement to Figure 1-Figure 8E**). Topic 1 and gene module 1 were enriched for genes characteristic of type 3 immunity, including *IL22, CXCL8, RORC, IL17F, IL23R* (Qi et al., 2021; Robinette et al., 2015), which were similarly found to have significantly increased expression in cluster 43 by cluster-based DGE analysis (**Supplementary Data 4**). In contrast, gene module 3 was enriched for genes characteristic of cytotoxic/type 1 immunity, including *CCL5, GZMA*, GNLY,* and *GZMB* (Hidalgo et al., 2008; Szabo et al., 2019a), and biological processes including positive regulation of interferon-gamma production (*GO:*0032729) and positive regulation of leukocyte mediated cytotoxicity (*GO:0001912*; **Supplementary Data 4**). Therefore, we classified cells in cluster 43 as group 3 ILCs and all remaining cells as group 1 ILCs (**Supplement to Figure 1-Figure 8E**).

Amongst group 1 ILCs, cluster 53 cells were noted to have nearly ubiquitous and significantly increased expression of many genes related to cell replication and/or division (e.g. *PCLAF, BIRC5, TOP2A, UBE2C*; Dabydeen et al., 2019; Giotti et al., 2019) and were enriched for similarly associated biological processes (e.g. cell cycle [*GO:0007049*], chromosome organization [*GO:0051276*], mitotic nuclear division [*GO:0140014*]; **Supplementary Data 4**). Therefore, we classified cluster 53 cells as cycling group 1 ILCs (**Supplement to Figure 1-Figure 8B**). Both topic 2 and gene module 2 had significantly increased expression of genes (e.g. *FCGR3A*, SLA-DRA*,* ribosomal genes; Cooper et al., 2001; Robinette and Colonna, 2016) and enrichment of biological processes (e.g. rRNA processing [*GO:0006364*], cellular response to interleukin-4 [*GO:0071353*], cellular response to toxic substance [*GO:0097237*]) related to cellular activation, while topic 3 had significantly increased expression of genes (e.g. *GNLY, GZMB, GZMA\**]) and enrichment of biological processes (e.g. natural killer cell activation [*GO:0030101*], natural killer cell mediated immunity [*GO:0002228*], leukocyte mediated cytotoxicity [*GO:0001909*]) related to cell cytotoxicity (**Supplementary Data 4**). Therefore, we classified activated group 1 ILCs as cells with low topic 3 weights (<0.05) and high gene module 2 detection scores (>0.4) or high topic 2 weights (>0.9; **Supplement to Figure 1-Figure 8F**). Remaining cells were classified as cytotoxic group 1 ILCs due to high topic 3 weighting (**Supplement to Figure 1-Figure 8F**). Collectively, we annotated four populations of ILCs, including activated group 1 ILCs, cytotoxic group 1 ILCs, cycling group 1 ILCs, and group 3 ILCs (**Supplement to Figure 1-Figure 8G-H**).

#### B lineage lymphocyte annotations

Cells identified as B lineage lymphocytes (clusters 0, 2, 4, 5, 8, 9, 10, 11, 13, 16, 20, 21, 22, 25, 27, 28, 30, 33, 38, 39, 40, 48; **Supplement to Figure 1-Figure 4**) were extracted to create a subset of data (as described in *Generating data subsets* methods). From this data subset, previous cell cluster assignments (as described in *Cell clustering* methods), topic modeling with weighted membership to three topics (as described in *Topic modeling* methods), and multidimensional DGE analysis with detection of four gene modules (as described in *Multidimensional differential gene expression analysis* methods) were assessed to identify DEGs and associated enriched biological processes by each method (**Supplement to Figure 1-Figure 9A-D**; **Supplementary Data 5**).

Clusters 25 and 33 belonged to a node most distantly related from other cells by hierarchical clustering (**Supplement to Figure 1-Figure 9B**). Cluster 25 had high weighting of topic 1, containing genes with significantly increased expression related to antibody secretion (e.g. *JCHAIN*, *CCR10, XBP1, PRDM1*; Kunkel et al., 2003; Reimold et al., 2001; Rinkenberger et al., 1996; Shapiro-Shelef et al., 2003), and similar genes related to antibody secretion were also found in cluster 25 by cluster-based DGE analysis (**Supplementary Data 5**). Similarly, both cluster 25 and topic 1 were enriched for biological processes including humoral immune response (*GO:0006959*), positive regulation of B cell activation (*GO:0050871*), positive regulation of protein exit from endoplasmic reticulum (*GO:0070863*), and protein glycosylation (*GO:0006486*; **Supplementary Data 5**), allowing us to classify cluster 25 as ASCs. Cluster 33 was closely related to cluster 25 by hierarchical clustering (**Supplement to Figure 1-Figure 9B**), but had lower weighting of topic 1 and expression of several genes related to antibody secretion and higher expression of several activation-associated and B cell canonical genes as stated in results, indicating cluster 33 to be B cells transitioning into ASCs, termed transitioning B cells.

Clusters 9, 13, and 30 also created a node of cells more distantly related from other B clusters by hierarchical clustering (**Supplement to Figure 1-Figure 9B**). Clusters 9, 13, and 30 all had significantly increased expression of genes associated with a resting phenotype, including *CCR7, SELL*, *CD40* (Bhattacharya et al., 2007; Förster et al., 1999; Zhang et al., 2021a), yet they all had significantly lower expression of activation markers such as *AICDA* and *CD86* (**Supplementary Data 5**; Engel et al., 1994; Muramatsu et al., 1999). Furthermore, all three clusters were also enriched for biological processes such as leukocyte homeostasis (*GO:0001776*), negative regulation of cellular protein metabolic processes (*GO:0032269*), homeostatic process (*GO:0042592*), and negative regulation of B cell activation (*GO:0050869*; **Supplementary Data 5**). Consequently, all cells belonging to clusters 9, 13, and 30 were annotated as resting B cells.

Topic 2 and gene module 3 both had significantly increased expression of genes related to cell division/replication (e.g. *PCLAF, BIRC5, TOP2A, UBE2C*; Dabydeen et al., 2019; Giotti et al., 2019) and enrichment of similarly-associated biological processes (e.g. cell division [*GO:0051301*], DNA-dependent DNA replication [*GO:0006261*], mitotic cell cycle [*GO:0000278*]; **Supplementary Data 5**). Consequently, remaining cells (excluding clusters 25, 33, 9, 13, 30) were defined as cycling B cells if they had high topic 2 weights (>0.32) or high gene module 3 detection scores (>0.06; **Supplement to Figure 1-Figure 9E**). Remaining cells were classified as activated B cells, as they did not fit the phenotype of cells classified as resting B cells, transitioning B cells, cycling B cells, or ASCs. Moreover, activated B cells still had higher expression of several activation associated genes and biological processes, as discussed in results. In total, we annotated five populations of B lineage lymphocytes, including ASCs, transitioning B cells, resting B cells, cycling B cells, and activated B cells (**Supplement to Figure 1-Figure 9F-G**).

#### Non-lymphocyte annotations

A data subset was created for myeloid lineage leukocytes (clusters 42, 49, 52; **Supplement to Figure 1-Figure 4**) as described in *Generating data subsets* methods. Since non-lymphocytes were not cells targeted for sequencing, cells identified as myeloid lineage leukocytes were annotated based only on cell cluster assignments (as described in *Cell clustering* methods) and associated hierarchical clustering, DGE, and biological process enrichment analyses (**Supplement to Figure 1-Figure 10A-B**; **Supplementary Data 6).** Cluster 49 was classified as macrophages based on expression of genes such as *CXCL2, CD68,* and complement component-encoding genes (e.g. *C1QA, C1QB, C1QC*; Betjes et al., 1991; De Filippo et al., 2008; Satpathy et al., 2012) and enrichment of biological processes including macrophage migration (*GO:1905517*), macrophage differentiation (*GO:0030225*), positive regulation of phagocytosis (*GO:0050766*), and toll-like receptor signaling pathway (*GO:0002224*; **Supplementary Data 6**). Cluster 42 was classified as DCs based on significantly elevated expression of *FLT3* and MHC-II encoding genes (e.g. *SLA-DRA*, SLA-DQB1*; Auray et al., 2016), and enrichment of biological processes including antigen processing and presentation of peptide antigen via MHC class II (*GO:0002495*), dendritic cell differentiation (*GO:0097028*), myeloid dendritic cell activation (*GO:*0001773), and positive regulation of protein processing (*GO:0010954*; **Supplementary Data 6**). Cluster 52 was annotated as mast cells based on expression of genes such as those encoding IgE receptors (e.g. *MS4A2, FCER1A*; Walczak-Drzewiecka et al., 2013) and enrichment of biological processes such as mast cell activation (*GO:0045576*), mast cell degranulation (*GO:0043303*), prostaglandin metabolic process (*GO:0006693*), and leukotriene metabolic process (*GO:0006691*; **Supplementary Data 6**). Collectively, the three myeloid lineage leukocyte cell clusters were annotated as DCs, macrophages, and mast cells (**Supplement to Figure 1-Figure 10C-D**).

From non-leukocytes (clusters 36, 45, 50; **Supplement to Figure 1-Figure 4**), a data subset was created as described in *Generating data subsets* methods. Since non-lymphocytes were not cells targeted for sequencing, cells identified as non-leukocytes were annotated based only on cell cluster assignments (as described in *Cell clustering* methods) and associated hierarchical clustering, DGE, and biological process enrichment analyses (**Supplement to Figure 1-Figure 10E-F**; **Supplementary Data 7**). Clusters 36 and 45 were most closely related by hierarchical clustering (**Supplement to Figure 1-Figure 10F**) and both had significantly elevated expression of genes characteristic of epithelial cells (e.g. encoding cytokeratins, solute carriers; **Supplementary Data 7**; Elmentaite et al., 2021; Han et al., 2018) and were thus merged and annotated as epithelial cells. Cluster 50 had significantly increased expression of genes characteristic of endothelial cells or fibroblasts, including *COL18A1, CCL19, CCL21, PECAM1, VCAM1* (Elmentaite et al., 2021; Han et al., 2018), and enrichment of biological processes including endothelial cell migration (*GO:0043542*), establishment of endothelial barrier (*GO:0061028*), regulation of extracellular matrix assembly (*GO:1901201*), and positive regulation of fibroblast migration (*GO:0010763*; **Supplementary Data 7**) and were consequently annotated as stromal cells. In all, the three non-leukocyte clusters were annotated as either epithelial or stromal cells (**Supplement to Figure 1-Figure 10G-H**).

#### Reference-based label transfer and mapping

Mapping and cell label prediction of porcine ileal cells to previously annotated cells from scRNA-seq datasets was performed using Seurat v3.9.9.9026 (Butler et al., 2018; Hao et al., 2021; Stuart et al., 2019). Each porcine ileal scRNA-seq sample (n=6) was treated as an individual query dataset, and scRNA-seq datasets of porcine PBMCs (Herrera-Uribe & Wiarda et al., 2021), human ileum (Elmentaite & Ross et al., 2020), and murine ileum (Xu et al., 2019) were treated as reference datasets. Post-quality control UMI count matrices, cell barcodes, and corresponding cell annotations were obtained for each reference dataset, and all samples from porcine PBMCs (n=7), control samples (pediatric, non-Crohn’s) from human ileum (n=8), and control samples (non-allergy) from murine ileum (n=14) were further used to create reference datasets. Cells with unknown identities (‘unknown’ annotation in porcine PBMCs and ‘unresolved’ annotation in murine ileum) or low UMIs (‘low UMI count’ designation in murine ileum) were further removed, leaving a total of 28,684; 11,302; and 27,159 cells from porcine PBMCs, human ileum, and murine ileum, respectively. For comparison between porcine and human or murine data, genes were filtered to contain only one-to-one human-to-pig or mouse-to-pig gene orthologs, as determined with BioMart (Smedley et al., 2015) and as previously described (Herrera-Uribe & Wiarda et al., 2021), within both reference and query datasets. For comparison between porcine ileal and PBMC datasets, genes were not filtered further.

Each reference dataset was next converted into a Seurat object and processed with SCT normalization, integration, scaling of the integrated assay, and PCA calculations as described in preceding methods. Annotation of cell lineage was also added to cells of each reference dataset, and assignment of each annotated cell type to a cell lineage can be seen in **Supplement to Figure 1-Figure 12**.

Each query dataset was processed using porcine genes for comparison to porcine PBMCs, only one-to-one human-to-pig gene orthologs for comparison to human ileum (termed humanized query data), or only one-to-one mouse-to-pig gene orthologs for comparison to murine ileum (termed murinized query data), resulting in a porcine, humanized, and murinized query dataset for each of the original six ileal scRNA-seq samples (18 query datasets total). Gene names from humanized and murinized query datasets were converted to human and murine gene names, respectively. Each query dataset was then converted into a Seurat object and processed with log normalization/scaling of the RNA assay and PCA calculations as described in preceding methods. Samples of query data were processed individually and not integrated.

Prediction and mapping scores to reference datasets were next generated for query samples using integrated assay data from reference datasets and RNA assay data from query datasets. Transfer anchors between each query with a respective reference dataset were calculated using the function *FindTransferAnchors()*, with canonical correlation analysis (CCA) reduction (recommended for cross-species comparisons), log normalization, and 30 dimensions specified. Transfer anchors were used as input for the function *TransferData()* to calculate predictions probabilities (range 0 to 1) for each query cell at the level of cell type and cell lineage annotations, again using CCA reduction and 30 dimensions. Mapping scores (range 0 to 1) were also calculated with the function *MappingScore()* using the calculated transfer anchors, reference dataset cell embeddings, and query dataset neighbors, weights matrix, and embeddings as inputs.

Query data from all six samples were merged back together, and prediction probabilities and mapping scores were summarized for each respective combination of reference identity assignment and query cluster ID. Mapping scores indicated poor (score=0) versus good (score=1) representation of query data in a reference dataset. Prediction probabilities indicated most highly probable similar cells in the reference dataset (0=not highly probable, 1=highly probable).

#### Pseudobulk analysis

Pseudobulk RNA-seq datasets were generated from ileum scRNA-seq samples using the Seurat function *AverageExpression()* to store mean-normalized counts from the RNA assay of Seurat objects. Pseudobulk samples were generated for each of the six ileal scRNA-seq samples either using reads after ambient RNA removal but before gene/cell filtering or using reads from the final dataset of 31,983 cells following gene/cell filtering. Using the R package, edgeR v3.30.3 (McCarthy et al., 2012; Robinson et al., 2010), pseudobulk counts were incorporated into a DGEList object with function *DGEList()*, and multidimensional scaling (MDS) plots to visualize sample-specific pseudobulk profiles were created with the function, *plotMDS()*.

#### Differential abundance analysis

Differential abundance (DA) analysis of cells originating from PP versus non-PP samples of ileum was performed using the R package, miloR v1.0.0 (Dann et al., 2021). All cells derived from PP and non-PP samples were taken as a data subset and reprocessed as described in preceding methods. In addition, SNNs were recalculated for the data subset with the Seurat function *FindNeighbors()*. The Seurat object was converted into a Milo object with incorporation of the SNN data graph calculated in Seurat with nearest neighbors (k) = 20 specified. Cell neighborhoods were created with the miloR function *makeNhoods()*, using 20% of cells in the dataset, k=20, and the significant number of PCs calculated for the data subset. Cell neighborhoods were visualized by overlay onto original t-SNE coordinates calculated for the original total dataset (including whole ileum samples) in order to promote cohesiveness with previous data visualizations. Within each cell neighborhood, the proportions of cells coming from each PP or non-PP sample was calculated. PP and non-PP samples were specified as treatment variables to compare in an experimental matrix, each with two replicates (pig 1 and 2). Distances between cell neighborhoods were calucalted with *calcNhoodDistance()* and the significant number of PCs for the data subset. DA testing was then performed within each cell neighborhood with the function *testNhoods()*. Cell neighborhoods were further annotated to correspond to the 26 cell types shown in **Figure 1D**, where a neighborhood was assigned as a cell type if >70% of cells in the neighborhood belonged to a single respective cell type annotation. If <70% of cells in a neighborhood were annotated as a single cell type, the neighborhood was classified as mixed. A neighborhood was considered differentially abundant at a significance level of 0.1 for corrected p-values.

#### scRNA-seq analysis of PBMC data

PBMC scRNA-seq data were processed independently but in parallel to ileal samples, with analyses mirroring those described for ileal scRNA-seq data analysis. A clustering resolution of 1.5 was used to cluster PBMCs, and cluster numbers were given a ‘p’ prefix (e.g. cluster p1 instead of cluster 1) to differentiate from ileal cell clusters. From PBMC clusters, gene expression was assessed to identify clusters of ILCs, using the same criteria as described for identification of porcine ileal ILCs and with additional query of NK cell genes described previously (Herrera-Uribe & Wiarda et al., 2021). By these criteria, we identified ILCs in PBMCs as cells belonging to clusters p0, p4, p26, p28, and p30 (**Supplement to Figure 6-Figure 1**).

The two PBMC samples derived from the same pigs used for ileal scRNA-seq have been published previously in the reference dataset used for porcine PBMC mapping and prediction comparisons described in above methods; however, reads were not corrected and trimmed further as was done in the previously published work (Herrera-Uribe & Wiarda et al., 2021).

#### scRNA-seq analysis of merged ileum/PBMC ILC data

Total datasets of porcine ileum and PBMCs were reduced using *DietSeurat()* to retain only the RNA assay and to remove dimensionality reductions, graphs, and scaled data. The two datasets were then combined with the Seurat function *merge()*, and clusters identified as ILCs from each dataset (clusters 1, 18, 43, 44, 53 in ileum and clusters p0, p4, p26, p28, and p30 in PBMCs) were specified within the function *subset()* to generate a data subset of only ILCs as described in *Generating data subsets* for ileal scRNA-seq analysis methods. In addition, SNNs were recalculated for the data subset with the Seurat function *FindNeighbors()*.

DA, multidimensional DGE, and biological process enrichment analyses were performed as described for ileum scRNA-seq data. DA analysis was performed between ILCs originating from ileum versus PBMC samples. Further annotation of cell neighborhoods into specific cell types (e.g. ileal cytotoxic group 1 ILCs, peripheral ILCs) was not performed.

Gene signatures for peripheral ILCs, ileal ILCs, ileal group 1 ILCs, and group 3 ILCs were created by further filtering of genes in modules created through multidimensional DGE analysis (**Supplementary Data 10**) that had high detection scores in ILC subsets specified in **Figure 6D-E**. To be included in a final gene signature, a gene from a specified module had to be expressed by >10% of cells in all target ILC subsets and for each target ILC subset compared pairwise to each non-target ILC subset have either (1) at least 2x the percentage of cells in the target ILC subset expressing the gene compared to the non-target ILC subset or (2) have a larger percentage of cells (but less than 2x as many) expressing the gene and have at least 2x the average log-normalized gene expression count for the target ILC subset compared to the non-target ILC subset. Summary statistics used for this filtering process are found in **Supplementary Data 10**.

From this rationale, genes in gene module 4 were further filtered to identify a peripheral ILC gene signature, where peripheral ILCs were the target subset and ileal cytotoxic group 1 ILCs, ileal cycling group 1 ILCs, ileal activated group 1 ILCs, and ileal group 3 ILCs were non-target subsets. Gene module 5 genes were filtered to identify an ileal ILC gene signature, where ileal cytotoxic group 1 ILCs, ileal cycling group 1 ILCs, ileal activated group 1 ILCs, and ileal group 3 ILCs were the target ILC subsets and peripheral ILCs were the non-target ILC subset. Genes in gene module 7 were filtered to identify an ileal group 1ILC gene signature, where ileal cytotoxic group 1 ILCs, ileal cycling group 1 ILCs, and ileal activated group 1 ILCs were target ILC subsets, and peripheral ILCs and group 3 ILCs were non-target subsets. The ileal group 3 ILC gene signature was created by filtering genes in gene modules 3 and 8, using ileal group 3 ILCs as the target subset and ileal cytotoxic group 1 ILCs, ileal cycling group 1 ILCs, ileal activated group 1 ILCs, and peripheral ILCs as non-target ILC subsets.

#### Gene name modifications

Several porcine genes did not have a gene symbol assigned by Ensembl in the genome annotation file used but did have a gene symbol in the NCBI database corresponding to the Ensembl gene identifier. In these cases, we substituted the Ensembl gene identifier with the NCBI gene symbol in our figures and text and have indicated this change with an asterix (*) at the end of the gene symbol used. Genes with duplicated gene symbols were also converted to Ensembl IDs when creating our gene annotation file, and these were converted back to gene symbols in our figures and text and denoted by an asterix (*) as well. The gene annotated as *HLA-DRA* was also converted to the updated pig-specific gene symbol, *SLA-DRA*. A comprehensive list of all such substitutions used in figures and text are in **Supplementary Data 11**.

#### Flow cytometry

Flow cytometry staining with cell viability dye and antibodies reactive to extracellular epitopes was performed as previously described (Wiarda et al., 2020). For antibody panels with intracellular CD79α detection, intracellular staining was completed after staining for extracellular markers, using the True-Nuclear Transcription Factor Buffer Set (BioLegend 424401) according to manufacturer’s instructions and as previously described (Boettcher et al., 2020a; Boettcher et al., 2020b). Cell events were acquired on a FACSymphony A5 flow cytometer (BD Biosciences), and data were analyzed with FlowJo v10.6.1 software (FlowJo, LLC) as previously described (Wiarda et al., 2020).

A panel to identify general populations of leukocytes, epithelial cells, T cells, and B cells (gating strategy shown in **Supplement to Figure 1-Figure 2B** and **Supplement to Figure 4-Figure 2A**) included Fixable Viability Dye-eFluor^TM^780 (ThermoFisher 65-0865-14); mouse α-pig CD45 (BioRad MCA1222GA) detected with rat α-mouse IgG1-BUV395 (BD 740234); rat α-mouse CD326-BV605 (CD326 aka EPCAM; BioLegend 118227); mouse α-pig CD3ε-PE-Cy7 (BD 561477); and mouse α-human CD79α-BV421 (BD 566225). A panel to identify γδ, CD4 αβ, and CD8 αβ T cells (gating strategy shown in **Supplement to Figure 4-Figure 2B**) included Fixable Viability Dye-eFluor^TM^780 (ThermoFisher 65-0865-14); mouse α-pig CD3ε-PE-Cy7 (BD 561477); mouse α-pig γδTCR-iFluor594 (primary antibody Washington State University PG2032; custom conjugation to iFluor594 performed by Caprico Biotechnologies); mouse α-pig CD4-PerCP-Cy5.5 (BD 561474); and mouse α-pig CD8β-PE (BioRad MCA5954PE). A panel to identify ileal ILCs (gating strategy shown in **Figure 5A**) included Fixable Viability Dye-eFluor^TM^780 (ThermoFisher 65-0865-14); mouse α-pig CD45 (BioRad MCA1222GA) detected with rat α-mouse IgG1-BUV395 (BD 740234); mouse α-pig CD2 (WSU PG2009) detected with rat α-mouse IgG3-BV650 (BD 744136); mouse α-pig CD3ε-PE-Cy7 (BD 561477); mouse α-human CD79α-BV421 (BD 566225); and mouse α-pig CD172α-FITC (BioRad MCA2312F).

#### Chromogenic immunohistochemistry

IHC staining for CD3ε protein was completed on formalin-fixed, paraffin-embedded (FFPE) tissues fixed for 24-30 hours in 10% neutral-buffered formalin (NBF; 3.7% formaldehyde) as previously described (Wiarda et al., 2020) with polyclonal rabbit α-human CD3ε antibody (Dako A0452) and polyclonal goat α-rabbit horseradish peroxidase (HRP)-conjugated antibody (Dako K4003). The same protocol was used for CD79α protein staining but replacing the primary antibody incubation with diluted monoclonal mouse α-human CD79α antibody (LifeSpan BioSciences LS-B4504) for 1 hour RT and replacing the secondary antibody incubation with HRP-labeled polyclonal goat α-mouse antibody (Dako K4000) for 30 minutes RT. IHC staining for pan-cytokeratin protein was carried out similar to CD3ε and CD79α IHC, but antigen retrieval was performed by incubation in Tris-EDTA buffer (10 mM Tris, 1 mM EDTA, pH 9.0; made in-house) for 20 minutes at 95 °*C*. Primary antibody incubation was performed by incubating slides with diluted monoclonal mouse α-human pan-cytokeratin antibody (Bio-Rad MCA1907T) overnight at 4 °*C* followed by secondary antibody incubation with HRP-labeled α-mouse antibody for 30 minutes RT.

#### Chromogenic RNA in-situ hybridization

FFPE tissues were obtained as described in IHC methods. Single-color chromogenic RNA ISH with *Sus scrofa*-specific channel 1 *TRDC* probe (Advanced Cell Diagnostics [ACD] 553141) was completed with the RNAscope 2.5 HD Reagent Kit-RED (ACD 322350) as previously described (Palmer et al., 2019) with the following modifications: (1) protease was applied for only 15 minutes at 40 °C and (2) slides were mounted and coverslipped as described elsewhere (Boettcher et al., 2020a). Dual chromogenic RNA ISH labeling with *Sus scrofa*-specific channel 1 (*CD8B*; ACD 815781) and channel 2 (*CD4*; ACD 491891-C2) probes was completed with the RNAscope 2.5 HD Duplex Reagent Kit (ACD 322430) as previously described (Boettcher et al., 2020a).

#### Dual protein immunofluorescence/fluorescent RNA in-situ hybridization

FFPE tissues were obtained as described in IHC methods. Dual immunofluorescence (IF) labelling of CD3ε protein and fluorescent RNA ISH of *ITGAE* or *IL22* transcripts was completed according to the RNAscope Multiplex Fluorescent v2 Assay Combined with Immunofluorescence – Integrated Co-Detection Workflow (ACD technical note MK 51-150), with some modifications. The RNAscope Multiplex Fluorescent Reagent Kit v2 (ACD 323100) was primarily used for RNA detection, with additional reagents outside of this kit pointed out in subsequent methods descriptions. Slides were baked, deparaffinized, rehydrated, and hydrogen peroxide incubations were completed as described in RNA ISH methods. Target retrieval was completed by submerging slides in 1X Co-Detection Target Retrieval solution (ACD 323165) for 15 minutes 95 °C, followed by rinsing with distilled water, 0.1% Tween in PBS (PBST) pH 7.2, and drawing of a hydrophobic barrier. CD3ε antibody (as used in IHC) was diluted in Co-detection Antibody Diluent (ACD 323160) and applied to slides for 1 hour at RT, followed by washing 2×2 minutes in PBST. Tissues were fixed in 10% NBF (3.7% formaldehyde) for 30 minutes at RT, then washed 3×2 minutes in PBST. Protease Plus was applied to tissue sections for 15 minutes at 40 °C, followed by rinsing with distilled water. The following steps were next completed sequentially at 40 °C, with 2×2 minute washes in 1X wash buffer between incubations: RNAscope custom channel 1 probe (*ITGAE* [ACD 590581] or *IL22* [ACD 590611]) 2 hours, AMP1 30 min, AMP2 30 min, AMP3 15min, HRP-C1 15 min, Opal 570 (Akoya Biosciences FP1488001KT) diluted in RNAscope Multiplex TSA Buffer (ACD 322809) 30 min, HRP blocker 15 min, goat α-rabbit IgG (H+L) F(ab’)-Alexa Fluor 488 (Invitrogen A11070) diluted in Co-detection Antibody Diluent 1 hour, DAPI 30 seconds. Following DAPI incubation, DAPI was removed and slides were mounted with ProLong Gold antifade reagent (Invitrogen P36930) and #1.5 thickness cover glass (Thermo Scientific 152450). Slides were dried in the dark at RT, placed in the dark at 4 °C overnight, and imaged within one week.

Fluorescent images were acquired with a Nikon A1R Confocal Microscope using a 60X Plan Apo oil objective with numerical aperture 1.40 via hybrid galvano/resonant scanner. Single solid lasers 405, 488 and 561 were used with standard and highly sensitive GaAsP detectors with center wavelength/bandwidth 450/50 (DAPI), 525/50 (Alexa Fluor 488) and 595/50 (Opal 570). 60X large image acquisition was used to generate a single high magnification, wide field-of-view image by automatically stitching multiple adjacent frames from a multipoint acquisition using the motorized stage on a fully automated inverted Ti2 Nikon microscope. Images were acquired and processed using NIS-Elements software (Nikon).

## Supporting information

Wiarda_SupplementaryFigures

Wiarda_SupplementaryData11

Wiarda_SupplementaryData09

Wiarda_SupplementaryData08

Wiarda_SupplementaryData10

Wiarda_SupplementaryData07

Wiarda_SupplementaryData06

Wiarda_SupplementaryData05

Wiarda_SupplementaryData02

Wiarda_SupplementaryData03

Wiarda_SupplementaryData04

Wiarda_SupplementaryData01

## DATA AVAILABILITY STATEMENT

Data and scripts will be available upon publication.

## AUTHOR CONTRIBUTIONS

JEW: Conceptualization, Methodology, Software, Validation, Formal analysis, Investigation, Data curation, Writing – original draft, Visualization JMT: Software, Data curation, Writing – review & editing SKS: Software, Data curation, Writing – review & editing CKT: Conceptualization, Writing – review & editing CLL: Conceptualization, Methodology, Investigation, Resources, Writing – review & editing, Supervision, Project administration, Funding acquisition

## ACKNOWLEDGEMENTS

We thank the following for their excellent contributions to this work: Dr. Kristen Byrne, Zahra Bond, and Sage Becker for technical and laboratory assistance; Dr. David Alt, Dr. Mike Baker, and the Iowa State University DNA Facility for library preparation and sequencing; Drs. Daniel Nielsen and Darrell Bayles for technologic assistance; Samuel Humphrey for flow cytometry expertise and services; Judith Stasko for histology expertise and services; Adrienne Shircliff for confocal microscopy expertise and services; Drs. Nicholas Gabler and Amy Vincent for collection of postmortem samples.

## ETHICS

Animal studies were reviewed and approved by the Institutional Animal Care and Use Committee (IACUC) at Iowa State University or the National Animal Disease Center, ARS, USDA.

## ABBREVIATIONS

ACD: Advanced Cell Diagnostics

ARS: Agricultural Research Service

ASC: antibody-secreting cell

BCR: B cell receptor

BSA: bovine serum albumin

c1: cluster 1

c2: cluster 2

CCA: canonical correlation analysis

cDC: conventional dendritic cell

CPT: Cell Preparation Tube

DA: differential abundance

DC: dendritic cell

DE: differentially expressed

DEG: differentially expressed gene

DGE: differential gene expression

DN: double-negative

DOE: Department of Energy

DTT: dithiothretol

DZ: dark zone

EDTA: ethylenediamine tetraacetic acid

FCS: fetal calf serum

FFPE: formalin-fixed, paraffin-embedded

FMO: fluorescence-minus-one

FSC-A: forward scatter area

FSC-H: forward scatter height

GALT: gut-associated lymphoid tissue

GC: germinal center

GO: Gene Ontology

HBSS: Hank’s balanced salt solution

HRP: horseradish peroxidase

IACUC: institutional animal care and use committee

IEL: intraepithelial lymphocyte

IF: immunofluorescence

IHC: immunohistochemistry

ILC: innate lymphoid cell

IQR: interquartile range

ISH: *in situ* hybridization

logFC: log fold-change

LTi: lymphoid tissue inducer

LZ: light zone

MDS: multidimensional scaling

NBF: neutral-buffered formalin

No: Sig no significance

pDC: plasmacytoid dendritic cell

Nhood: neighborhood

NK: natural killer

NKT: natural killer T

ORAU: Oak Ridge Associated Universities

ORISE: Oak Ridge Institute for Science and Education

PBMC: peripheral blood mononuclear cell

PBS: phosphate buffered saline

PBST: phosphate buffered saline with Tween

PC: principle component

PCA: principle component analysis

PP: Peyer’s patch

RNA-seq: RNA sequencing

RT: room temperature

scRNA-seq: single-cell RNA sequencing

SNN: shared nearest neighbor

SSC-A: side scatter height

T-IEL: intraepithelial T lymphocyte

t-SNE: t-distributed stochastic neighbor embedding

TA: transit amplifying

TCR: T cell receptor

T_FH_: T follicular helper

T_FR_: T follicular regulatory

T_reg_: T regulatory

UMI: unique molecular identifier

USDA: United States Department of Agriculture

## Footnotes

* Ensembl identifiers found in gene annotation were converted to gene symbols; refer to methods section ‘*Gene name modifications*’ for more details

