## Supplementary material for "Porcine intestinal innate lymphoid cells and lymphocyte spatial context revealed through single-cell RNA sequencing": Wiarda_SupplementaryFigures

### Supplement to Figure 1-Figure 1

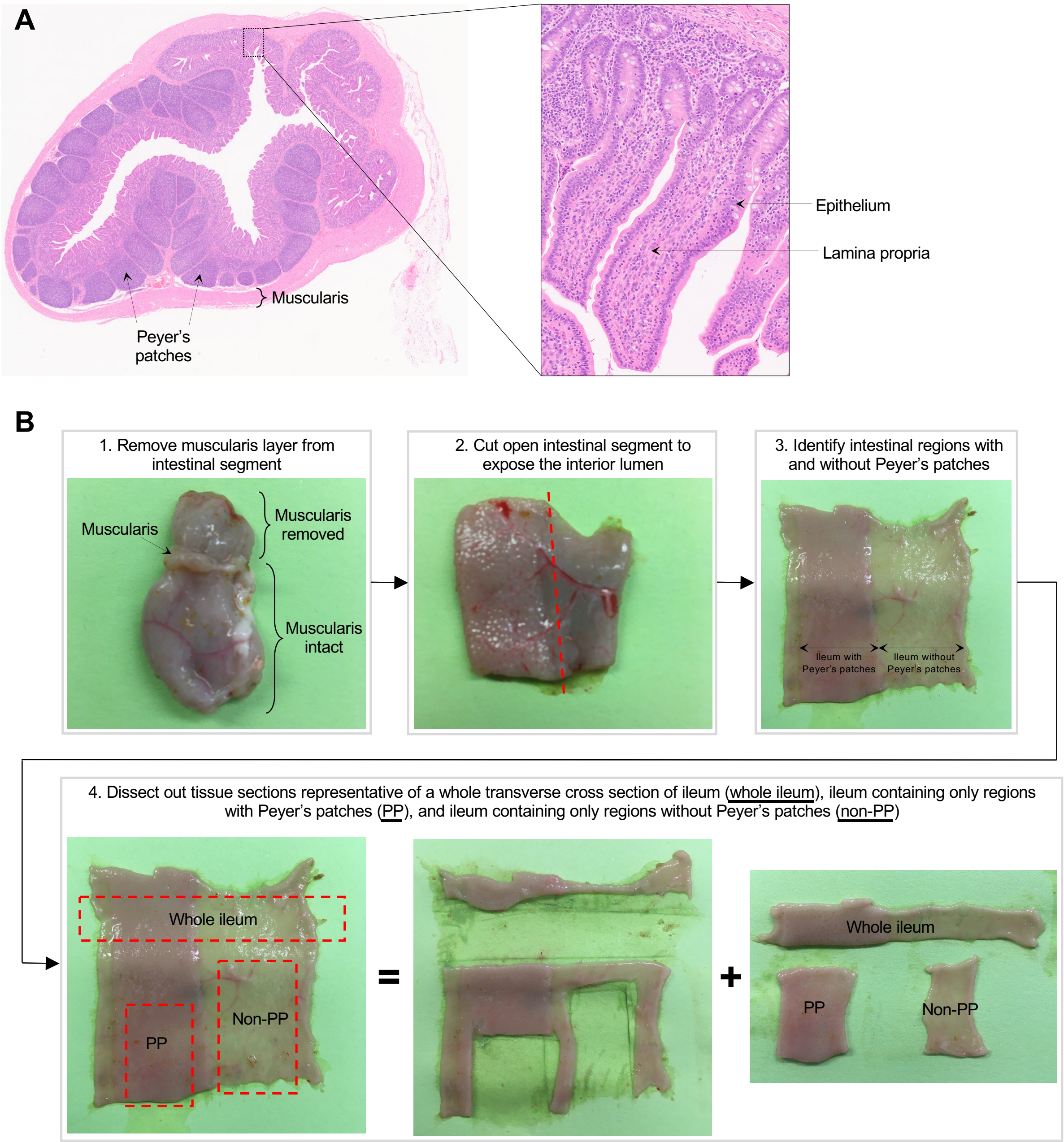

**Supplement to Figure 1–Figure 1. Histology and dissection of porcine ileum.**

**(A)** Transverse cross section of ileum collected from distal small intestine of a seven-week-old pig used for scRNA-seq as shown in **Figure 1A** and stained with hemotoxylin (purple) and eosin (pink). Histological structures corresponding to tissue muscularis, Peyer's patches, epithelium, and lamina propria are indicated.

**(B)** Representative images of tissue dissections performed on ileum to obtain a whole transverse cross section of ileum (whole ileum), ileum containing only regions with Peyer's patches (PP), and ileum containing only regions without Peyer's patches (non-PP). Images shown in **B** were from a nine-week-old pig and were not from animals used for scRNA-seq.

Abbreviations: PP (Peyer's patch); scRNA-seq (single-cell RNA sequencing)

### Supplement to Figure 1-Figure 2

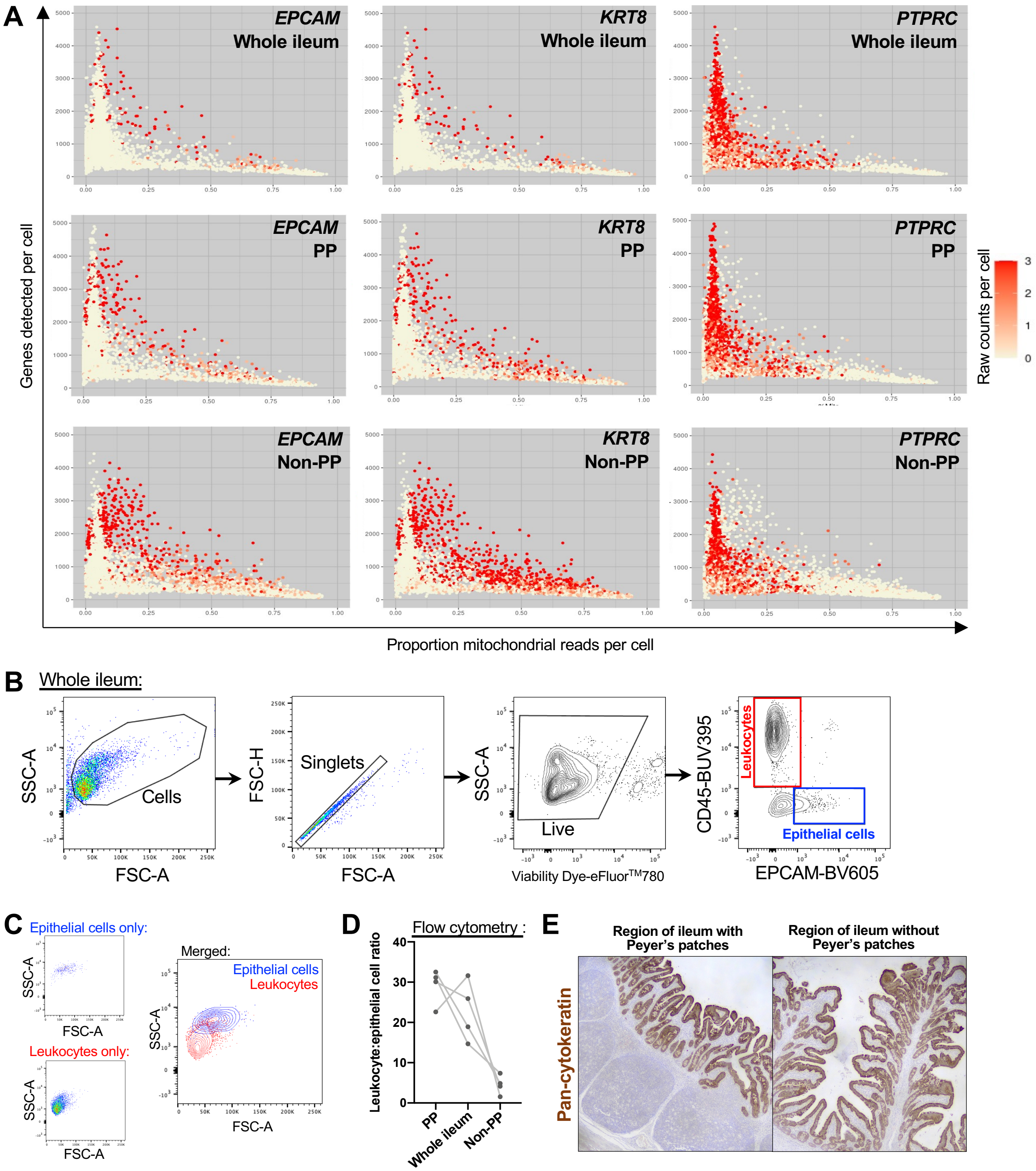

**Supplement to Figure 1–Figure 2. Enrichment of poor-quality epithelial cells in ileum without Peyer's patches.**

**(A)** Plots of two quality control metrics (genes detected per cell [y-axis] and proportion mitochondrial reads per cell [x-axis]) used to identify and filter out poor quality cells from scRNA-seq data. Each point represents a single cell. Point fill color corresponds to raw gene counts for epithelial genes *EPCAM* (left) and *KRT8* (center) and pan-leukocyte gene *PTPRC* (right). Plots are shown from whole ileum (top), PP (middle), and non-PP (bottom) samples collected from one seven-week-old pig used for scRNA-seq.

**(B)** Flow cytometry gating strategy to identify leukocytes ($CD45^+$) and epithelial cells ($EPCAM^+$) from total live cells isolated from porcine ileum. Gating is shown for a whole ileum sample (containing both regions with and without Peyer's patches).

**(C)** Overlay of gated leukocytes and epithelial cells from **B** onto original forward- and side-scatter coordinates to infer parameters of cell size and complexity, respectively, that are consistent with leukocytes and epithelial cells.

**(D)** Ratio of the number of leukocytes to the number of epithelial cells (y-axis) identified by flow cytometry gating shown in **B**. Cells were isolated from three types of ileal dissections (x-axis); samples derived from different ileal dissections of the same pig are connected with a grey line.

**(E)** IHC staining for epithelial pan-cytokeratin protein (brown) in a region of ileum with Peyer's patches (left) or without Peyer's patches (right).

Flow cytometry and IHC experiments were not performed on animals used for scRNA-seq. Flow cytometry experiments shown in **B-D** were conducted using four six-week-old pigs. IHC staining in **E** was completed on a five-week-old pig.

Abbreviations: FSC-A (forward scatter area); FSC-H (forward scatter height); IHC (immunohistochemistry); PP (Peyer's patch); scRNA-seq (single-cell RNA sequencing); SSC-A (side scatter area)

### Supplement to Figure 1-Figure 3

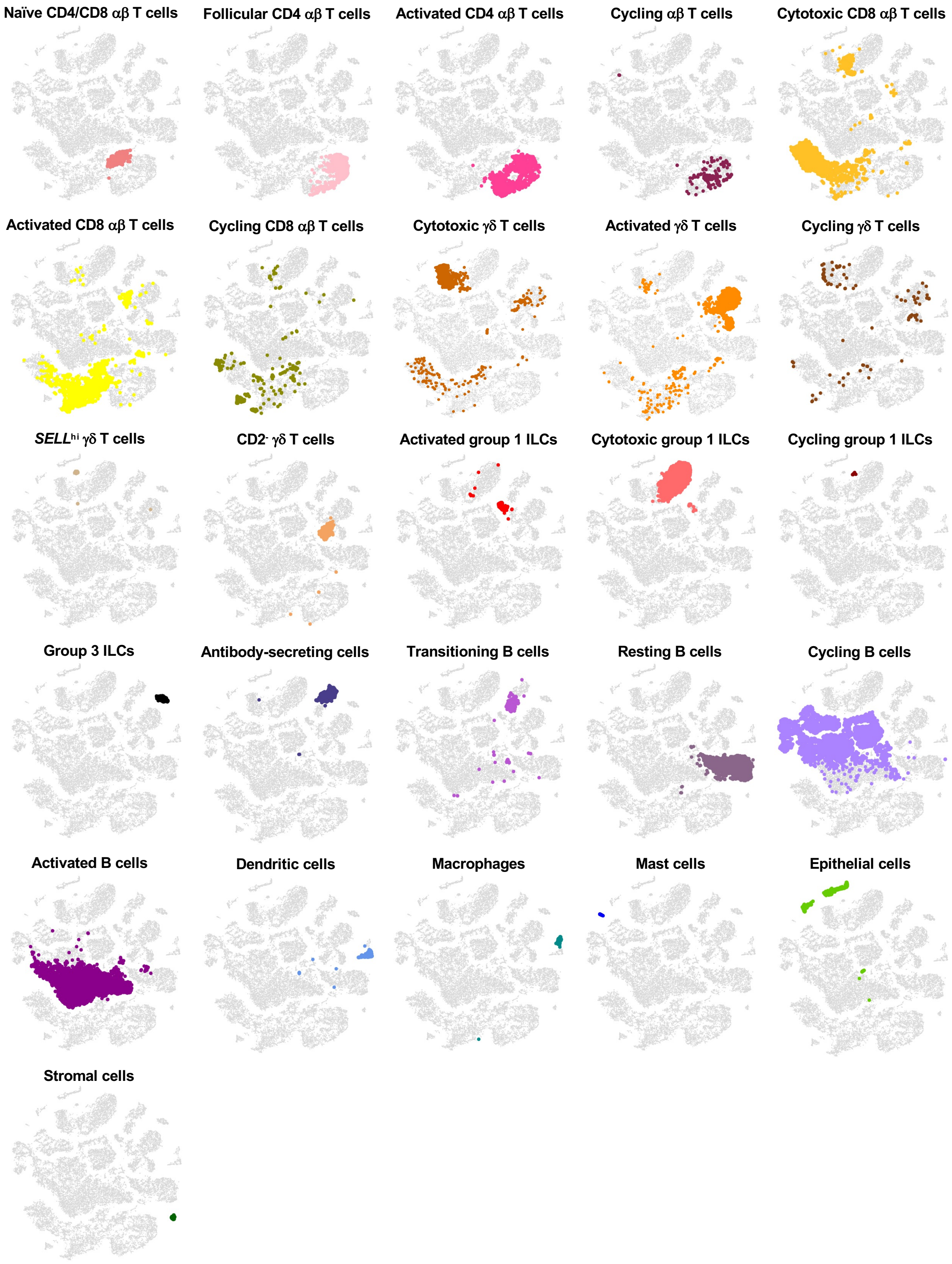

**Supplement to Figure 1-Figure 3. Overlay of cell type annotations onto t-SNE visualization of cells from porcine ileum scRNA-seq data.** Overlay of 26 annotated cell types onto two-dimensional t-SNE visualization of 31,983 cells recovered from ileum of two seven-week-old pigs via scRNA-seq. Each point represents a single cell. Cell type is indicated in a respective panel by one of 26 colors corresponding to cell types shown in **Figure 1D**, while all other cells not corresponding to a specified cell type are shown in light grey.
Abbreviations: ILC (innate lymphoid cell); scRNA-seq (single-cell RNA sequencing); t-SNE (t-distributed stochastic neighbor embedding)

### Supplement to Figure 1-Figure 4

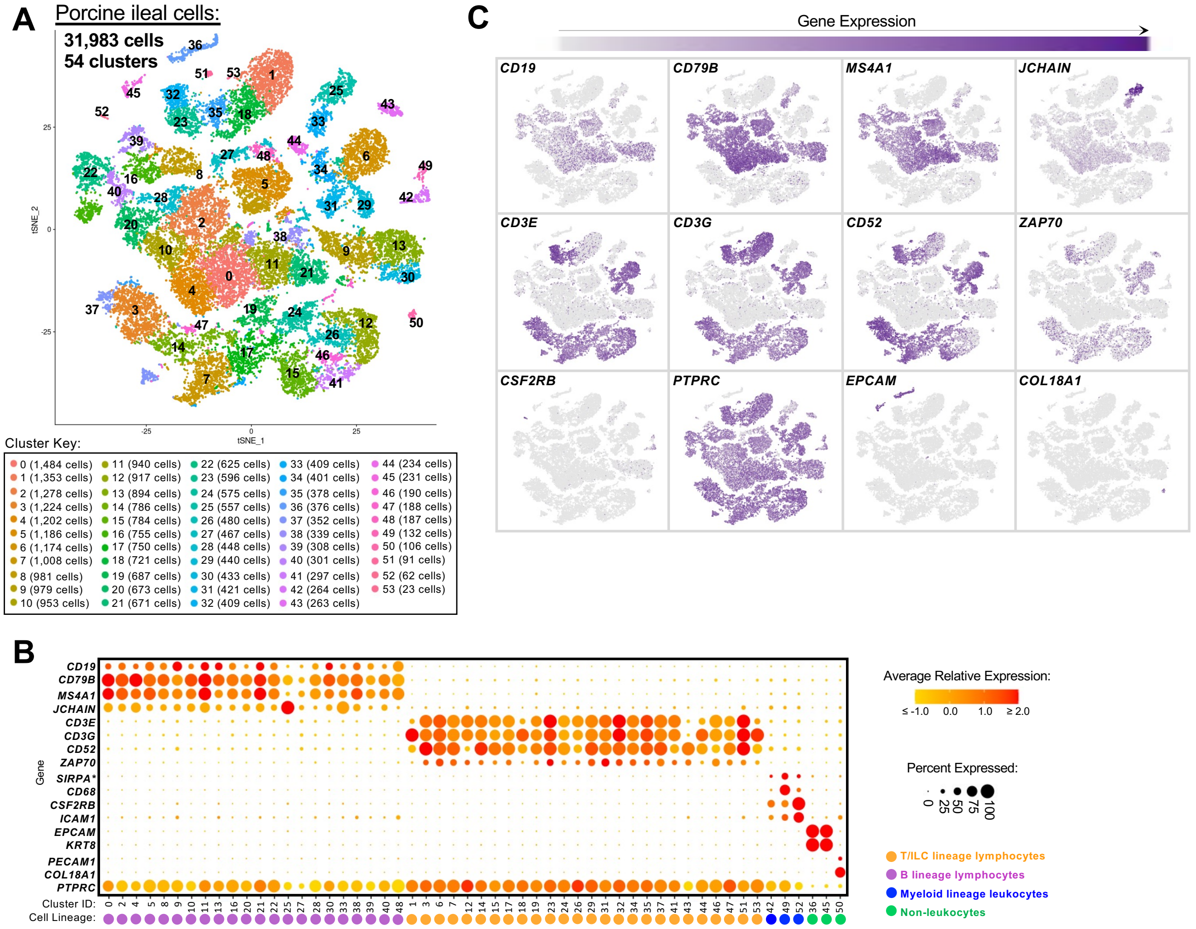

**Supplement to Figure 1-Figure 4. Cell lineage annotation of cells from porcine ileum scRNA-seq data.**

**(A)** Two-dimensional t-SNE visualization of 31,983 cells recovered from porcine ileum via scRNA-seq. Each point represents a single cell; color of a point corresponds to one of 54 cell clusters a cell belonged to, with more transcriptionally similar cells belonging to the same cluster. The number of cells belonging to each cluster is listed in the cluster key.

**(B)** Gene expression patterns of selected canonical genes (y-axis) across cell clusters shown in **A** (x-axis). Within the plot, size of a dot corresponds to the percentage of cells expressing a gene within a cell cluster; color of a dot corresponds to average expression level of a gene for those cells expressing it within a cell cluster relative to all other cells in the dataset shown in **A**. Below cluster ID on the x-axis, the color of a circle corresponds to cell lineage annotation given to each cluster.

**(C)** Expression of a subset of canonical genes from **B** overlaid onto two-dimensional t-SNE visualization coordinates of cells shown in **A**. Color of a point corresponds to expression level of a specified gene within a cell relative to all other cells in the dataset shown in **A**.

scRNA-seq data shown in **A-C** were derived from ileum of two seven-week-old pigs.

*Ensembl identifiers found in gene annotation were converted to gene symbols; refer to methods section '*Gene name modifications*' for more details

Abbreviations: ILC (innate lymphoid cell); scRNA-seq (single-cell RNA sequencing); t-SNE (t-distributed stochastic neighbor embedding)

### Supplement to Figure 1-Figure 5

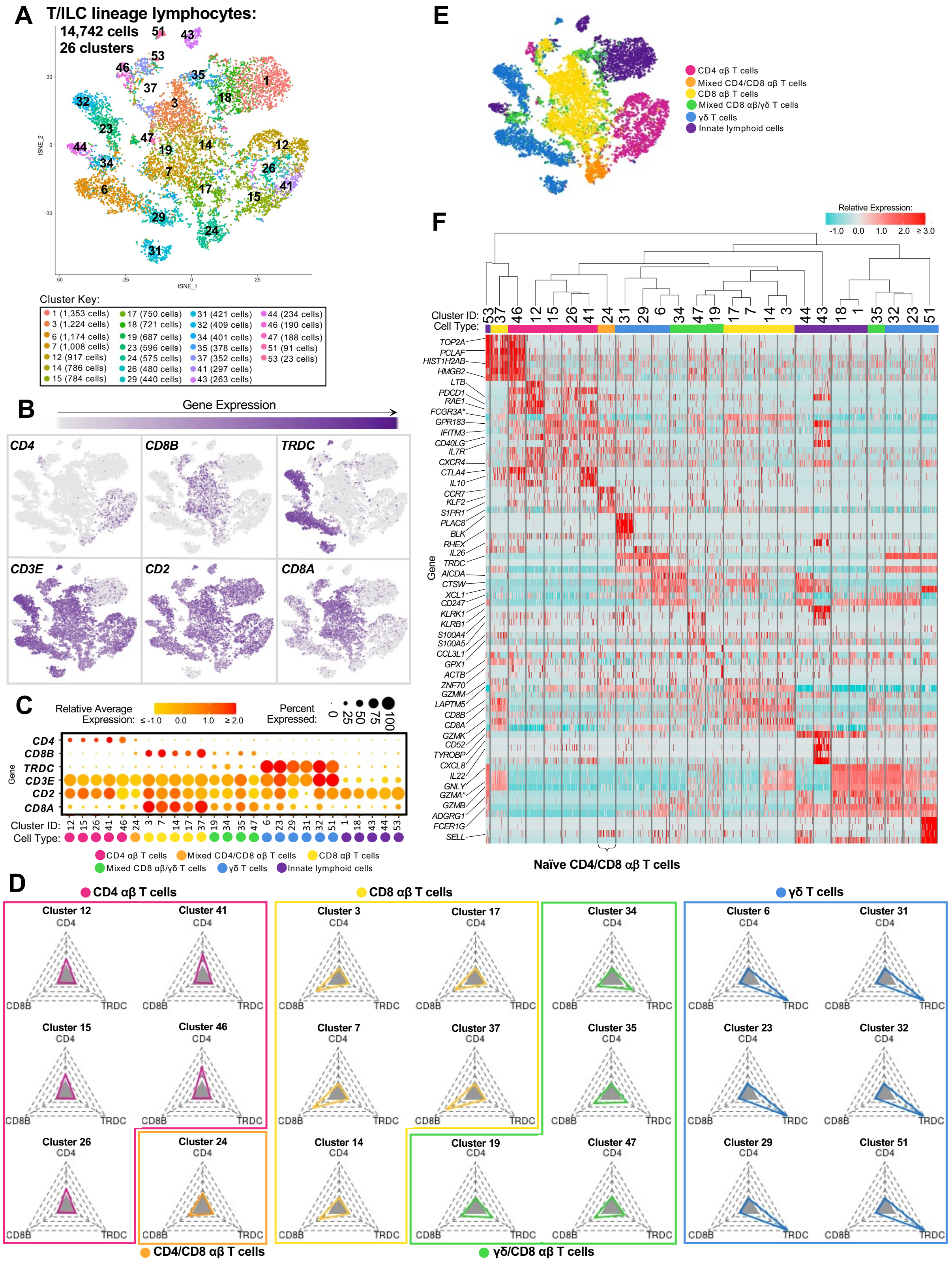

**Supplement to Figure 1-Figure 5. Annotation of T/ILC lineage lymphocytes from porcine ileum scRNA-seq data.**
**(A)** Two-dimensional t-SNE visualization of 14,742 cells recovered from porcine ileum via scRNA-seq and classified as T/ILC lineage lymphocytes in **Figure 1C** & **Supplement to Figure 1-Figure 4B**. Each point represents a single cell; color of a point corresponds to one of 26 cell clusters a cell belonged to, with more transcriptionally similar cells belonging to the same cell cluster. The number of cells belonging to each cell cluster is listed in the cluster key.
**(B)** Gene expression of selected canonical genes overlaid onto two-dimensional t-SNE visualization coordinates of cells shown in **A**. Color of a point corresponds to expression level of a specified gene within a cell relative to all other cells in the dataset shown in **A**.
**(C)** Gene expression patterns of selected canonical genes (y-axis) across cell clusters shown in **A** (x-axis). Within the plot, size of a dot corresponds to the percentage of cells expressing a gene within a cell cluster. Color of a dot corresponds to average expression level of a gene for those cells expressing it within a cell cluster relative to other cells in the dataset shown in **A**. Below cluster ID on the x-axis, the color of a circle corresponds to a further T/ILC classification given to each cell cluster.
**(D)** Radial plots showing the percentage of cells expressing *CD4, CD8B,* or *TRDC* within each respective cell cluster shown in **A**. Shaded grey triangles at the center of each plot indicate the minimum limit of positive detection (10%), while the outer limits of each plot are equivalent to 100%. T/ILC classification given to each cluster based on the percentage of cells positive for each gene are shown by plot color and outline.
**(E)** Two-dimensional t-SNE visualization of cells shown in **A**, where color of a point now corresponds to T/ILC classification given to a cell cluster in **C-D**.
**(F)** Heatmap of top differentially expressed genes within each cell cluster shown in **A**. Up to five differentially expressed genes with the highest positive logFC values were selected for each cell cluster. Genes were differentially expressed in a specified cluster relative to the average of all other cells in the dataset shown in **A**. Gene expression profiles from up to 100 cells of each cluster are shown in the heatmap. Each column represents a single cell. Selected gene names are shown on the y-axis, and cell cluster IDs are shown on the x-axis. T/ILC classification given to each cluster in **C-E** is indicated below each cell cluster ID on the x-axis. Hierarchical relationships of cell clusters are shown using a phylogenetic tree at the top of the heatmap.
scRNA-seq data shown in **A-F** were derived from ileum of two seven-week-old pigs.
*Ensembl identifiers found in gene annotation were converted to gene symbols; refer to methods section '*Gene name modifications*' for more details
Abbreviations: ILC (innate lymphoid cell); logFC (log fold-change); scRNA-seq (single-cell RNA sequencing); t-SNE (t-distributed stochastic neighbor embedding)

### Supplement to Figure 1-Figure 6

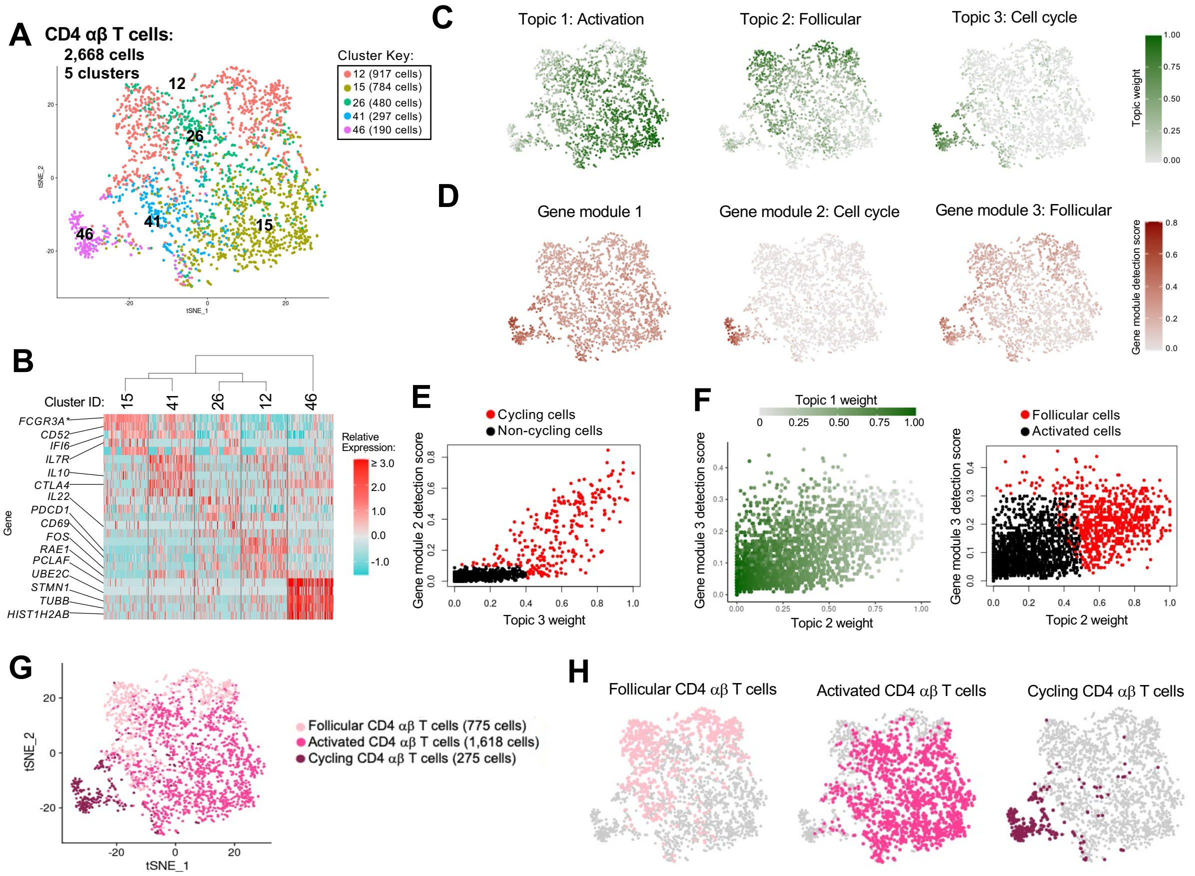

**Supplement to Figure 1-Figure 6. Annotation of CD4 αβ T cells from porcine ileum scRNA-seq data.**

**(A)** Two-dimensional t-SNE visualization of 2,668 cells recovered from porcine ileum via scRNA-seq and classified as CD4 αβ T cells in **Supplement to Figure 1-Figure 5C-F**. Each point represents a single cell; color of a point corresponds to one of five cell clusters a cell belonged to, with more transcriptionally similar cells belonging to the same cell cluster. The number of cells belonging to each cell cluster is listed in the cluster key.
**(B)** Heatmap of top differentially expressed genes within each cell cluster shown in **A**. Up to five differentially expressed genes with the highest positive logFC values were selected for each cell cluster. Genes were differentially expressed in a specified cluster relative to the average of all other cells in the dataset shown in **A**. Gene expression profiles from up to 100 cells of each cell cluster are shown in the heatmap, with each column representing a single cell. Selected gene names are shown on the y-axis, and cell cluster IDs are shown on the x-axis. Hierarchical relationships of cell clusters are shown using a phylogenetic tree at the top of the heatmap.
**(C)** Topic weights from topic modeling of cells shown in **A** overlaid onto two-dimensional t-SNE visualization coordinates. Color of a point corresponds to proportional weighting of a topic within a cell, where total weighting across all topics in each cell is equal to one.
**(D)** Gene module detection scores from multidimensional differential gene expression analysis of cells shown in **A** overlaid onto two-dimensional t-SNE visualization coordinates. Color of a point corresponds to detection score for a gene module within a cell.
**(E)** Scatter plot of gene module 2 detection scores (y-axis) versus topic 3 weights (x-axis) for all cells shown in **A**. Each point represents a single cell. Cells with a gene module 2 detection score >0.1 and/or topic 3 weight >0.4 are shown in red and were annotated as cycling cells. Remaining cells are shown in black and were classified as non-cycling cells.
**(F)** Scatter plots of gene module 3 detection scores (y-axis) versus topic 2 weights (x-axis) for all non-cycling cells shown in **E**. Each point represents a single cell. Left: point fill corresponds to topic 1 weights. Right: cells with a gene module 3 detection score >0.3 and/or topic 2 weight > topic 1 weight are shown in red and annotated as follicular cells. Remaining cells are shown in black and annotated as activated cells.
**(G)** CD4 αβ T cell annotations established in **B-F** overlaid onto two-dimensional t-SNE visualization coordinates of cells shown in **A**. Color of a point corresponds to cell type annotation. The number of cells belonging to each cell type is listed in the key on the right.
**(H)** Overlay of individual cell types onto two-dimensional t-SNE visualization shown in **G**. Cell type is indicated in a respective panel by one of three colors corresponding to cell types shown in **G**, while all other cells not corresponding to a specified cell type are shown in light grey.
scRNA-seq data shown in **A-H** were derived from ileum of two seven-week-old pigs.
*Ensembl identifiers found in gene annotation were converted to gene symbols; refer to methods section '*Gene name modifications*' for more details
Abbreviations: logFC (log fold-change); scRNA-seq (single-cell RNA sequencing); t-SNE (t-distributed stochastic neighbor embedding)

### Supplement to Figure 1-Figure 7

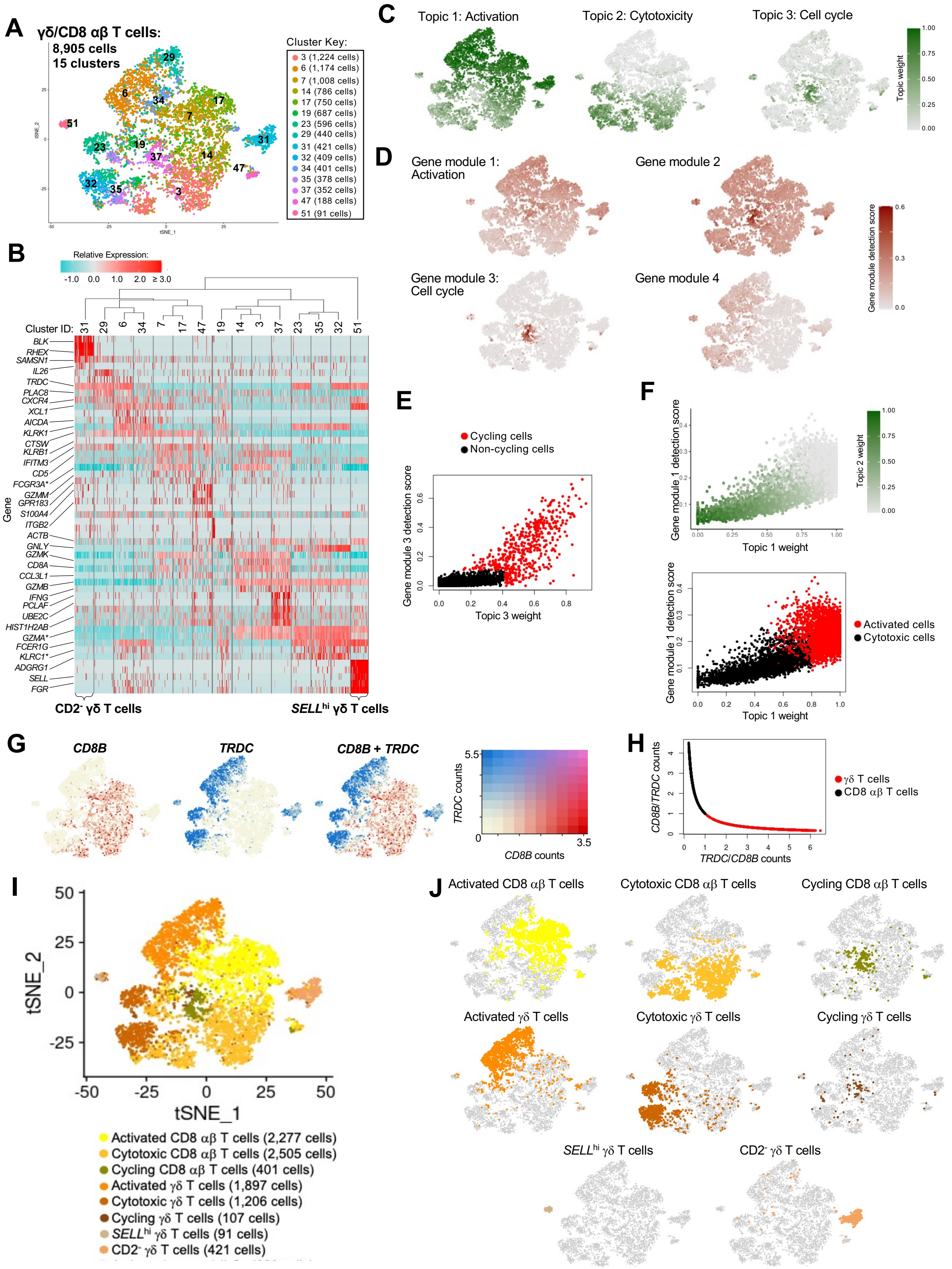

**Supplement to Figure 1-Figure 7. Annotation of γδ and CD8 αβ T cells from porcine ileum scRNA-seq data.**

**(A)** Two-dimensional t-SNE visualization of 8,905 cells recovered from porcine ileum via scRNA-seq and classified as γδ T cells, CD8 αβ T cells, or mixed γδ/CD8 αβ T cells in **Supplement to Figure 1-Figure 5C-F**. Each point represents a single cell; color of a point corresponds to one of 15 cell clusters a cell belongs to, with more transcriptionally similar cells belonging to the same cell cluster. The number of cells belonging to each cell cluster is listed in the cluster key.

**(B)** Heatmap of top differentially expressed genes within each cell cluster shown in **A**. Up to five differentially expressed genes with the highest positive logFC values were selected for each cell cluster. Genes were differentially expressed in a specified cell cluster relative to the average of all other cells in the dataset shown in **A**. Gene expression profiles from up to 100 cells of each cluster are shown in the heatmap, with each column representing a single cell. Selected gene names are shown on the y-axis, and cell cluster IDs are shown on the x-axis. Hierarchical relationships of cell clusters are shown using a phylogenetic tree at the top of the heatmap. At the bottom of the heatmap, cells in cluster 31 were annotated as $CD2^-$ γδ T cells, and cells in cluster 51 were annotated as *SELL*$^{hi}$ γδ T cells.

**(C)** Topic weights from topic modeling of cells shown in **A** overlaid onto two-dimensional t-SNE visualization coordinates. Color of a point corresponds to proportional weighting of a topic within a cell, where total weighting across all topics in each cell is equal to one.

**(D)** Gene module detection scores from multidimensional differential gene expression analysis of cells shown in **A** overlaid onto two-dimensional t-SNE visualization coordinates. Color of a point corresponds to detection score for a gene module within a cell.

**(E)** Scatter plot of gene module 3 detection scores (y-axis) versus topic 3 weights (x-axis) for all cells shown in **A**, excluding $CD2^-$ γδ T cells (cell cluster 31) and *SELL*$^{hi}$ γδ T cells (cell cluster 51). Each point represents a single cell. Cells with a gene module 3 detection score >0.11 and/or topic 3 weight >0.41 are shown in red and annotated as cycling cells. Remaining cells are shown in black and classified as non-cycling cells.

**(F)** Scatter plots of gene module 1 detection scores (y-axis) versus topic 1 weights (x-axis) for all non-cycling cells shown in **E**. Each point represents a single cell. Upper: point fill corresponds to topic 2 weights. Lower: cells with a gene module 1 detection score >0.25 and/or topic 1 weight at least 4x greater than topic 2 weight are shown in red and classified as activated cells. Remaining cells are shown in black and classified as cytotoxic cells.

**(G)** Relative gene expression levels of *CD8B* (left), *TRDC* (right), and merged *CD8B* and *TRDC* overlaid onto two-dimensional t-SNE visualization coordinates shown in **A**.

**(H)** Scatter plot of ratios of log-normalized *CD8B/TRDC* (y-axis) and *TRDC/CD8B* (x-axis) counts for all cells shown in **E**. Each point represents a single cell. Cells with a *TRDC/CD8B* ratio >1 are shown in red and classified as γδ T cells. Remaining cells are shown in black and classified as CD8 αβ T cells.

**(I)** γδ and CD8 αβ T cell annotations established from combined classifications in **A-H** overlaid onto two-dimensional t-SNE visualization coordinates of cells shown in **A**. Color of a point corresponds to cell type annotation. The number of cells belonging to each cell type is listed in the color key on the bottom.

**(J)** Overlay of individual cell types onto two-dimensional t-SNE visualization shown in **I**. Cell type is indicated in a respective panel by one of eight colors corresponding to cell types shown in **I**, while all other cells not corresponding to a specified cell type are shown in light grey.

scRNA-seq data shown in **A-J** were derived from ileum of two seven-week-old pigs.

*Ensembl identifiers found in gene annotation were converted to gene symbols; refer to methods section ‘*Gene name modifications*’ for more details

Abbreviations: logFC (log fold-change); scRNA-seq (single-cell RNA sequencing); t-SNE (t-distributed stochastic neighbor embedding)

### Supplement to Figure 1-Figure 8

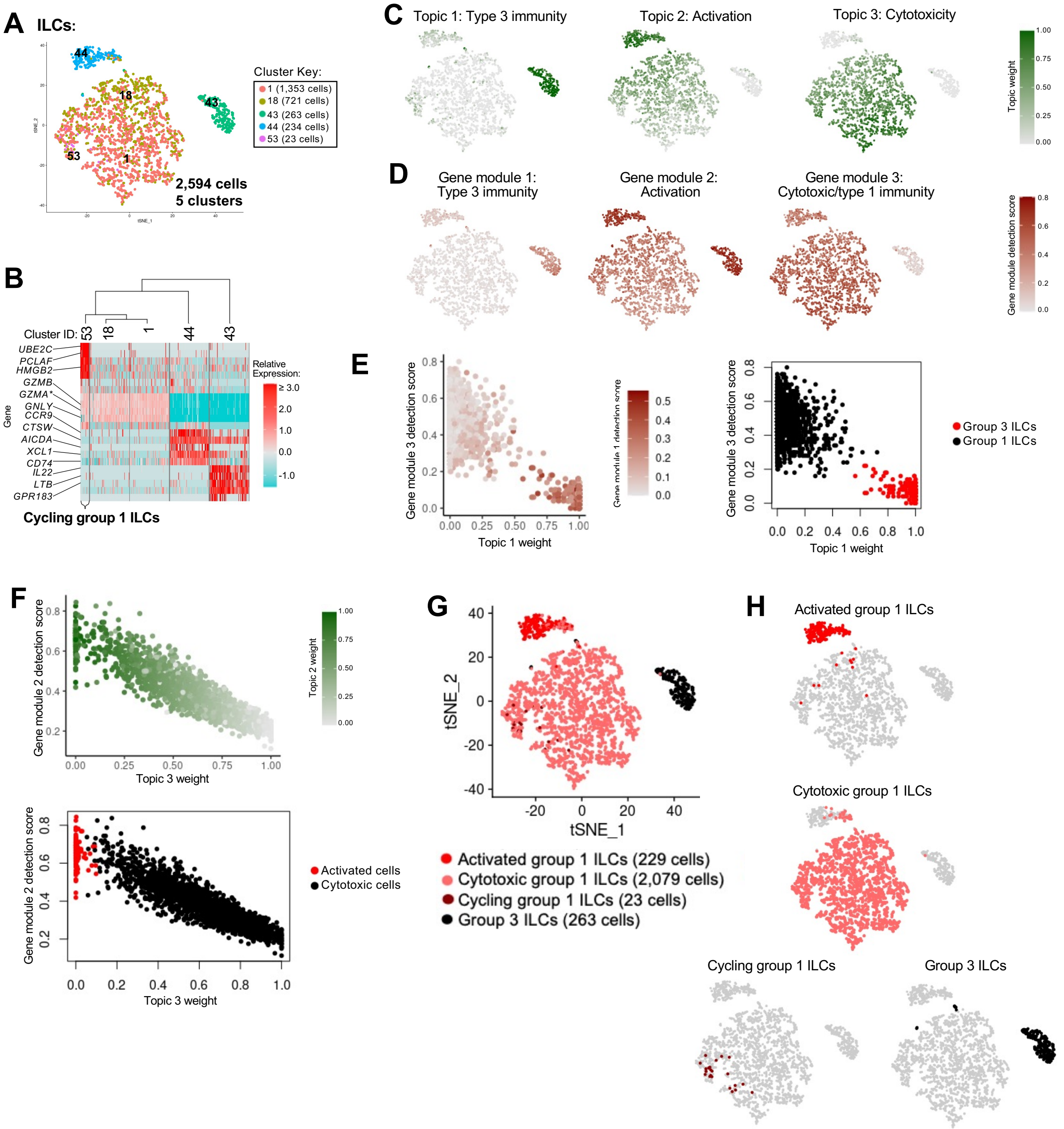

**Supplement to Figure 1-Figure 8. Annotation of ILCs from porcine ileum scRNA-seq data.** **(A)** Two-dimensional t-SNE visualization of 2,594 cells recovered from porcine ileum via scRNA-seq and classified as ILCs in **Supplement to Figure 1-Figure 5C-F**. Each point represents a single cell; color of a point corresponds to one of five cell clusters a cell belongs to, with more transcriptionally similar cells belonging to the same cell cluster. The number of cells belonging to each cell cluster is listed in the cluster key.
**(B)** Heatmap of top differentially expressed genes within each cell cluster shown in **A**. Up to five differentially expressed genes with the highest positive logFC values were selected for each cell cluster. Genes were differentially expressed in a specified cell cluster relative to the average of all other cells in the dataset shown in **A**. Gene expression profiles from up to 100 cells of each cell cluster are shown in the heatmap, with each column representing a single cell. Selected gene names are shown on the y-axis, and cell cluster IDs are shown on the x-axis. Hierarchical relationships of cell clusters are shown using a phylogenetic tree at the top of the heatmap. At the bottom of the heatmap, cluster 53 was annotated as cycling group 1 ILCs.
**(C)** Topic weights from topic modeling of cells shown in **A** overlaid onto two-dimensional t-SNE visualization coordinates. Color of a point corresponds to proportional weighting of a topic within a cell, where total weighting across all topics in each cell is equal to one.
**(D)** Gene module detection scores from multidimensional differential gene expression analysis of cells shown in **A** overlaid onto two-dimensional t-SNE visualization coordinates. Color of a point corresponds to detection score for a gene module within a cell.
**(E)** Scatter plots of gene module 3 detection scores (y-axis) versus topic 1 weights (x-axis) for all cells shown in **A**. Each point represents a single cell. Left: point fill corresponds to gene module 1 detection score. Right: cells belonging to cluster 43 are shown in red and were annotated as group 3 ILCs. Remaining cells are shown in black and classified as group 1 ILCs.
**(F)** Scatter plots of gene module 2 detection scores (y-axis) versus topic 3 weights (x-axis) for cells belonging to cell clusters 1, 18, or 44 in **A-B**. Each point represents a single cell. Upper: point fill corresponds to topic 2 weights. Lower: cells with a topic 3 weight <0.05 and gene module 2 detection scores >0.4 or topic 2 weights >0.9 are shown in red and annotated as activated cells. Remaining cells are shown in black and annotated as cytotoxic cells.
**(G)** ILC annotations established in **B-F** overlaid onto two-dimensional t-SNE visualization coordinates of cells shown in **A**. Color of a point corresponds to cell type annotation. The number of cells belonging to each cell type is listed in the color key on the bottom.
**(H)** Overlay of individual cell types onto two-dimensional t-SNE visualization shown in **G**. Cell type is indicated in a respective panel by one of four colors corresponding to cell types shown in **G**, while all other cells not corresponding to a specified cell type are shown in light grey.
scRNA-seq data shown in **A-H** were derived from ileum of two seven-week-old pigs.
*Ensembl identifiers found in gene annotation were converted to gene symbols; refer to methods section '*Gene name modifications*' for more details
Abbreviations: ILC (innate lymphoid cell); logFC (log fold-change); scRNA-seq (single-cell RNA sequencing); t-SNE (t-distributed stochastic neighbor embedding)

### Supplement to Figure 1-Figure 9

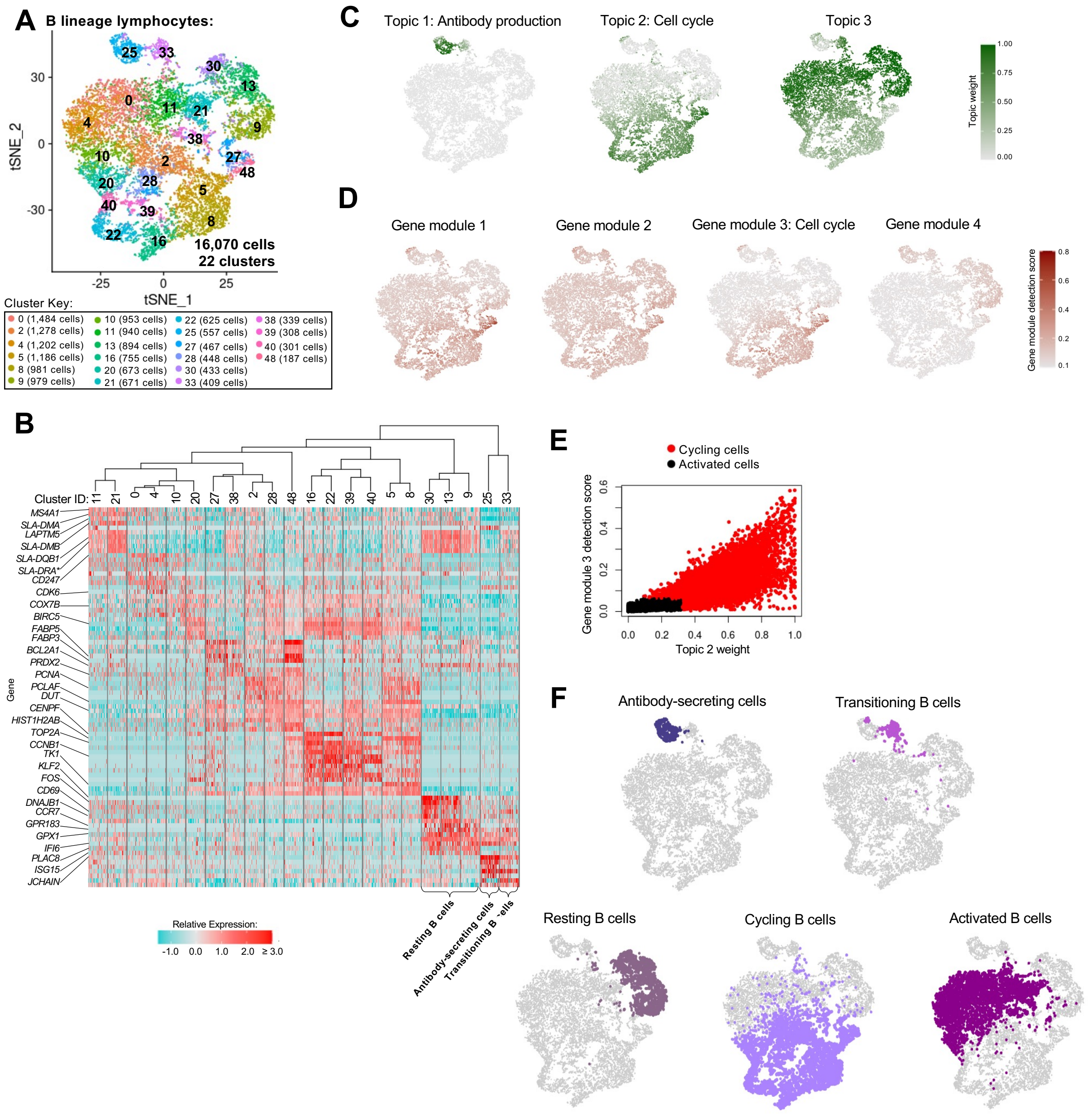

**Supplement to Figure 1-Figure 9. Annotation of B lineage lymphocytes from porcine ileum scRNA-seq data.**

**(A)** Two-dimensional t-SNE visualization of 16,070 cells recovered from porcine ileum via scRNA-seq and classified as B lineage lymphocytes in **Figure 1C** & **Supplement to Figure 1-Figure 4B**. Each point represents a single cell; color of a point corresponds to one of 22 cell clusters a cell belongs to, with more transcriptionally similar cells belonging to the same cell cluster. The number of cells belonging to each cell cluster is listed in the cluster key.

**(B)** Heatmap of top differentially expressed genes within each cell cluster shown in **A**. Up to five differentially expressed genes with the highest positive logFC values were selected for each cell cluster. Genes were differentially expressed in a specified cell cluster relative to the average of all other cells in the dataset shown in **A**. Gene expression profiles from up to 100 cells of each cell cluster are shown in the heatmap, with each column representing a single cell. Selected gene names are shown on the y-axis, and cell cluster IDs are shown on the x-axis. Hierarchical relationships of clusters are shown using a phylogenetic tree at the top of the heatmap. At the bottom of the heatmap, cluster 33 was annotated as transitioning B cells, cluster 25 as antibody-secreting cells, and clusters 9, 13, and 30 as resting B cells.

**(C)** Topic weights from topic modeling of cells shown in **A** overlaid onto two-dimensional t-SNE visualization coordinates. Color of a point corresponds to proportional weighting of a topic within a cell, where total weighting across all topics in each cell is equal to one.

**(D)** Gene module detection scores from multidimensional differential gene expression analysis of cells shown in **A** overlaid onto two-dimensional t-SNE visualization coordinates. Color of a point corresponds to detection score for a gene module within a cell.

**(E)** Scatter plots of gene module 3 detection scores (y-axis) versus topic 2 weights (x-axis) for all cells shown in **A,** excluding resting B cells (clusters 9, 13, 20), transitioning B cells (cluster 33) and antibody-secreting cells (cluster 25). Each point represents a single cell. Cells with gene module 3 detection scores >0.06 and/or topic 2 weights >0.32 are shown in red and annotated as cycling cells. Remaining cells are shown in black and annotated as activated cells.

**(F)** B lineage lymphocyte annotations established in **B-E** overlaid onto two-dimensional t-SNE visualization coordinates of cells shown in **A**. Cell type is indicated in a respective panel by one of five colors corresponding to annotated cell types, while all other cells not corresponding to a specified cell type are shown in light grey.

scRNA-seq data shown in **A-F** were derived from ileum of two seven-week-old pigs.

*Ensembl identifiers found in gene annotation were converted to gene symbols; refer to methods section '*Gene name modifications*' for more details

Abbreviations: logFC (log fold-change); scRNA-seq (single-cell RNA sequencing); t-SNE (t-distributed stochastic neighbor embedding)

### Supplement to Figure 1-Figure10

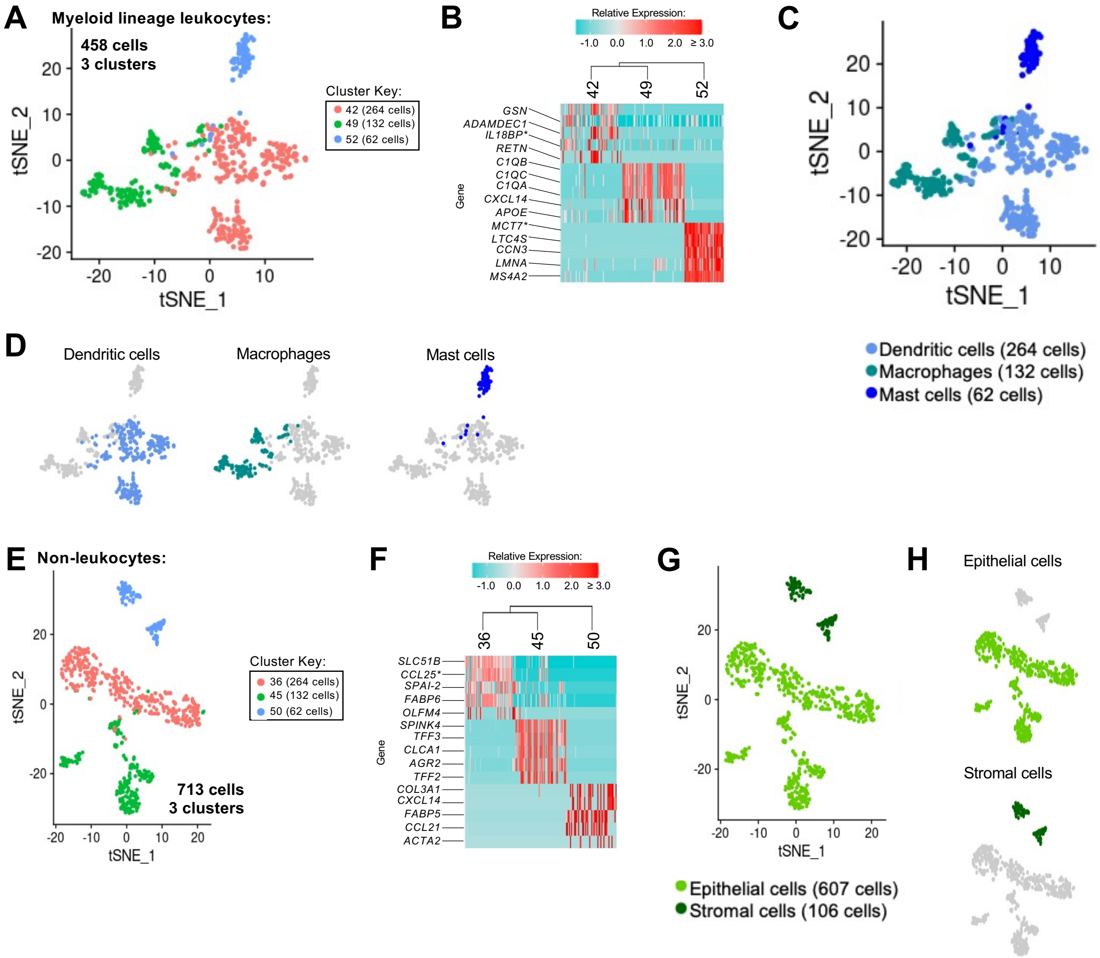

**Supplement to Figure 1-Figure 10. Annotation of non-lymphocytes from porcine ileum scRNA-seq data.**

**(A)** Two-dimensional t-SNE visualization of 458 cells recovered from porcine ileum via scRNA-seq and classified as myeloid lineage leukocytes in **Figure 1C** & **Supplement to Figure 1-Figure 4B**. Each point represents a single cell; color of a point corresponds to one of three cell clusters a cell belongs to, with more transcriptionally similar cells belonging to the same cell cluster. The number of cells belonging to each cell cluster is listed in the cluster key.
**(B)** Heatmap of top differentially expressed genes within each cell cluster shown in **A**. Up to five differentially expressed genes with the highest positive logFC values were selected for each cell cluster. Genes were differentially expressed in a specified cell cluster relative to the average of all other cells in the dataset shown in **A**. Gene expression profiles from up to 100 cells of each cell cluster are shown in the heatmap, with each column representing a single cell. Selected gene names are shown on the y-axis, and cell cluster IDs are shown on the x-axis. Hierarchical relationships of cell clusters are shown using a phylogenetic tree at the top of the heatmap.
**(C)** Myeloid lineage leukocyte annotations established in **A-B** overlaid onto two-dimensional t-SNE visualization coordinates of cells shown in **A**. Color of a point corresponds to cell type annotation. The number of cells belonging to each cell type is listed in the color key below.
**(D)** Overlay of individual cell types onto two-dimensional t-SNE visualization shown in **C**. Cell type is indicated in a respective panel by one of three colors corresponding to cell types shown in **C**, while all other cells not corresponding to a specified cell type are shown in light grey.
**(E)** Two-dimensional t-SNE visualization of 713 cells recovered from porcine ileum via scRNA-seq and classified as non-leukocytes in **Figure 1C** & **Supplement to Figure 1-Figure 4B**. Each point represents a single cell; color of a point corresponds to one of three cell clusters a cell belonged to, with more transcriptionally similar cells belonging to the same cell cluster. The number of cells belonging to each cell cluster is listed in the cluster key.
**(F)** Heatmap of top differentially expressed genes within each cell cluster shown in **E**. Up to five differentially expressed genes with the highest positive logFC values were selected for each cell cluster. Genes were differentially expressed in a specified cell cluster relative to the average of all other cells in the dataset shown in **E**. Gene expression profiles from up to 100 cells of each cell cluster are shown in the heatmap, with each column representing a single cell. Gene names are shown on the y-axis, and cell cluster IDs are shown on the x-axis. Hierarchical relationships of cell clusters are shown using a phylogenetic tree at the top of the heatmap.
**(G)** Myeloid lineage leukocyte annotations established in **E-F** overlaid onto two-dimensional t-SNE visualization coordinates of cells shown in **E**. Color of a point corresponds to cell type annotation. The number of cells belonging to each cell type is listed in the color key below.
**(H)** Overlay of individual cell types onto two-dimensional t-SNE visualization shown in **G**. Cell type is indicated in a respective panel by one of three colors corresponding to cell types shown in **G**, while all other cells not corresponding to a specified cell type are shown in light grey.
scRNA-seq data shown in **A-H** were derived from ileum of two seven-week-old pigs.
*Ensembl identifiers found in gene annotation were converted to gene symbols; refer to methods section '*Gene name modifications*' for more details
Abbreviations: logFC (log fold-change); scRNA-seq (single-cell RNA sequencing); t-SNE (t-distributed stochastic neighbor embedding)

#### Supplement to Figure 1-Figure 11

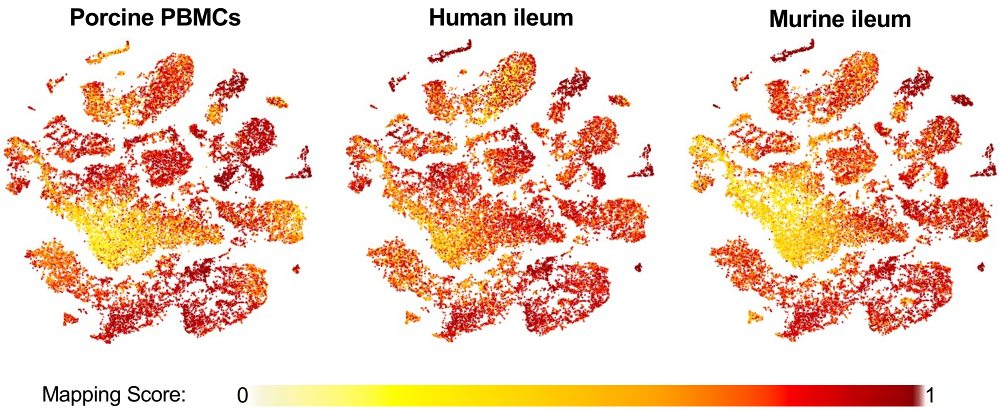

**Supplement to Figure 1-Figure 11. Mapping scores of porcine ileal cells to reference scRNA-seq datasets.**

Mapping scores from mapping of porcine ileum scRNA-seq query data to reference scRNA-seq datasets of porcine PBMCs (left), human ileum (center), and murine ileum (right) overlaid onto two-dimensional t-SNE visualization of porcine ileum scRNA-seq data shown in **Figure 1C-D**. Each point represents a single cell; the color of each point indicates mapping score to a corresponding reference dataset. Higher mapping scores indicate better representation of a cell from porcine ileum in a specified reference dataset.

Query scRNA-seq data were derived from ileum of two seven-week-old pigs.

Abbreviations: PBMC (peripheral blood mononuclear cell); scRNA-seq (single-cell RNA sequencing); t-SNE (t-distributed stochastic neighbor embedding)

### Supplement to Figure 1-Figure 12

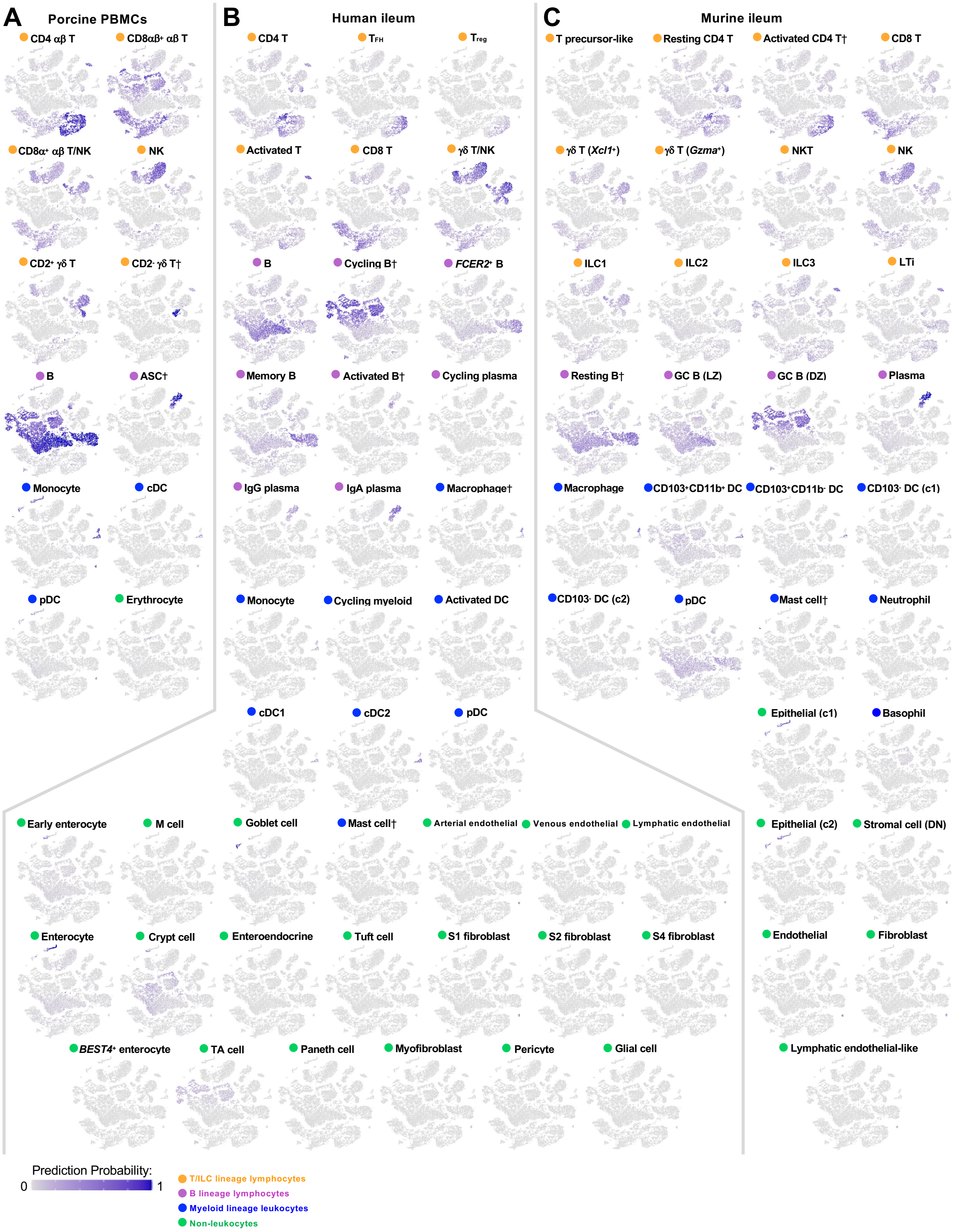

**Supplement to Figure 1-Figure 12. Prediction scores of porcine ileal cells to annotated cell types in reference scRNA-seq datasets.**

Prediction probabilities for porcine ileum scRNA-seq query data from label transfer of annotated cell types in reference scRNA-seq datasets of **(A)** porcine PBMCs, **(B)** human ileum, and **(C)** murine ileum overlaid onto two-dimensional t-SNE visualization of porcine ileum scRNA-seq data shown in **Figure 1C-D**. Each point represents a single cell; the color of each point indicates prediction probability to a corresponding cell type annotation from a specified reference dataset. Cell lineage of each annotated reference cell type is indicated by a circle next to each respective annotated cell type name. Within each of **A, B,** or **C**, cumulative prediction probabilities for each cell across all annotated reference cell types are equal to one.

Query scRNA-seq data were derived from ileum of two seven-week-old pigs.

† Identical cell type annotations were given to cells in both porcine ileum and a reference scRNA-seq dataset. Cell type annotations were given to each dataset by independent rationales, and identical annotations do not necessarily indicate identical cell types were recovered from both porcine ileum and reference data.

Abbreviations: ASC (antibody-secreting cell); c1 (cluster 1); c2 (cluster 2); cDC (convnentional dendritic cell); DC (dendritic cell); DN (double-negative); DZ (dark zone); GC (germinal center); ILC (innate lymphoid cell); LTi (lymphoid tissue inducer); LZ (light zone); PBMC (peripheral blood mononuclear cell); pDC (plasmacytoid dendritic cell); NK (natural killer); NKT (natural killer T); scRNA-seq (single-cell RNA sequencing); t-SNE (t-distributed stochastic neighbor embedding); TA (transit amplifying); $T_{FH}$ (T follicular helper); $T_{reg}$ (T regulatory)

### Supplement to Figure 2-Figure 1

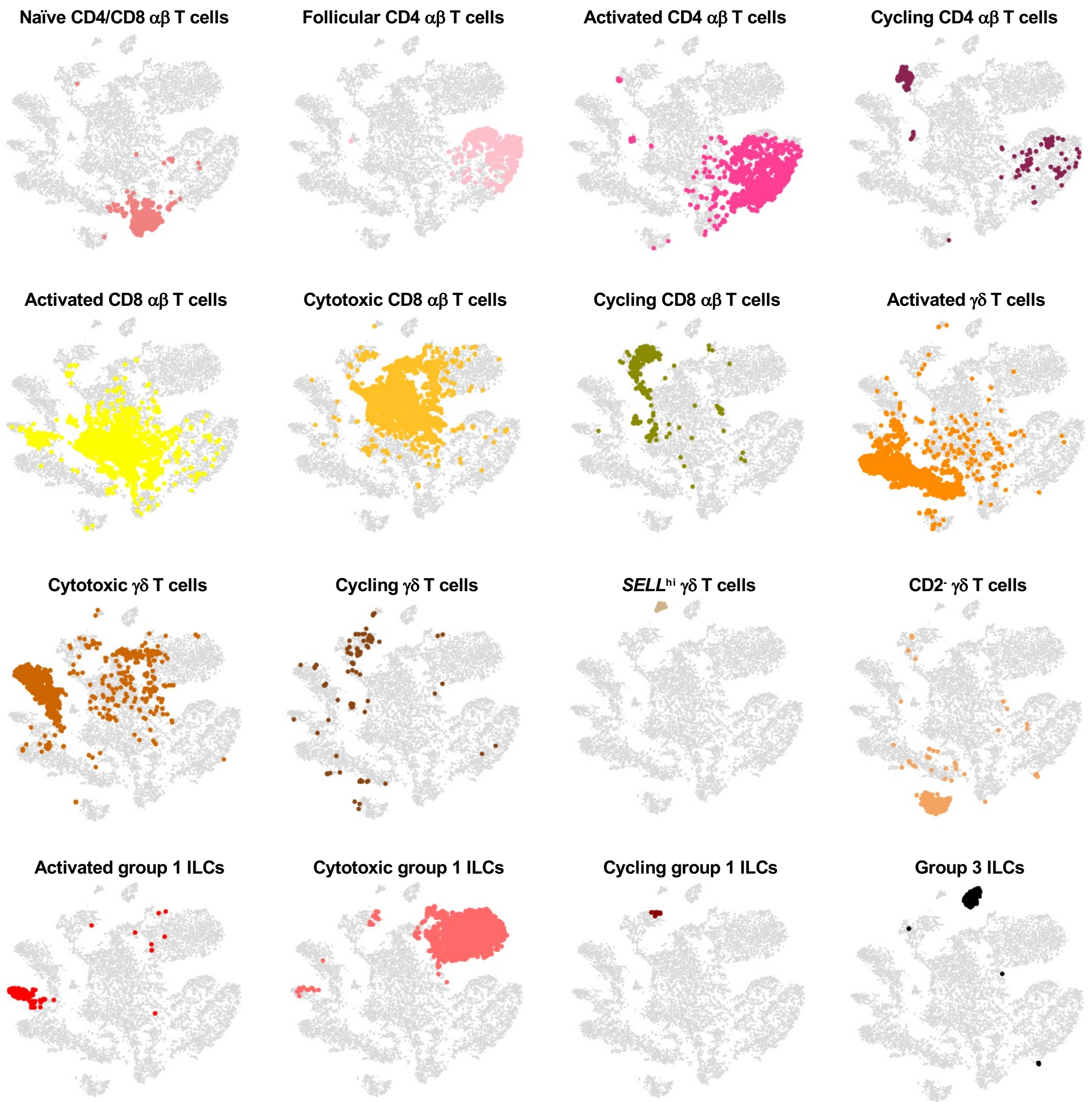

**Supplement to Figure 2-Figure 1. Overlay of T/ILC annotations onto t-SNE visualization of cells from porcine ileum scRNA-seq data.** Overlay of 16 annotated T/ILC types onto two-dimensional t-SNE visualization of 14,742 cells recovered from ileum of two seven-week-old pigs via scRNA-seq and classified as T/ILC lineage lymphocytes in **Figure 1C** & **Supplement to Figure 1-Figure 4B**. Each point represents a single cell. Cell type is indicated in a respective panel by one of 16 colors corresponding to cell types shown in **Figure 2A**, while all other cells not corresponding to a specified cell type are shown in light grey.
Abbreviations: ILC (innate lymphoid cell); scRNA-seq (single-cell RNA sequencing); t-SNE (t-distributed stochastic neighbor embedding)

#### Supplement to Figure 2-Figure 2

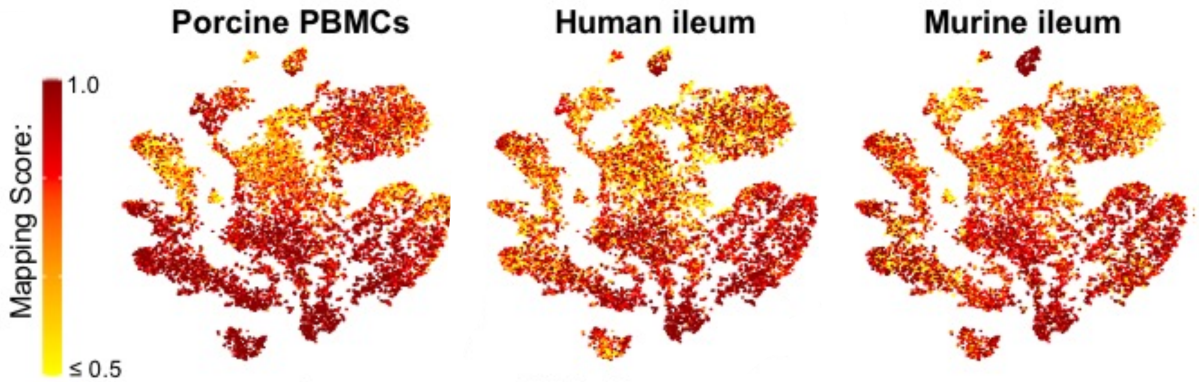

**Supplement to Figure 2-Figure 2. Overlay of mapping scores onto t-SNE reduction of T/ILC lineage lymphocytes from porcine ileum scRNA-seq data.**
Mapping scores from mapping of porcine ileum scRNA-seq query data to reference scRNA-seq datasets of porcine PBMCs (left), human ileum (center), and murine ileum (right). Mapping scores are the same as shown in **Supplement to Figure 1-Figure 11** but are now shown only for T/ILC lineage lymphocytes overlaid onto two-dimensional t-SNE visualization of porcine ileum scRNA-seq data shown in **Figure 2A**. Each point represents a single cell; the color of each point indicates mapping score to a corresponding reference dataset. Higher mapping scores indicate better representation of a cell from porcine ileum in a specified reference dataset.
Query scRNA-seq data were derived from ileum of two seven-week-old pigs.
Abbreviations: ILC (innate lymphoid cell); PBMC (peripheral blood mononuclear cell); scRNA-seq (single-cell RNA sequencing); t-SNE (t-distributed stochastic neighbor embedding)

### Supplement to Figure 2-Figure 3

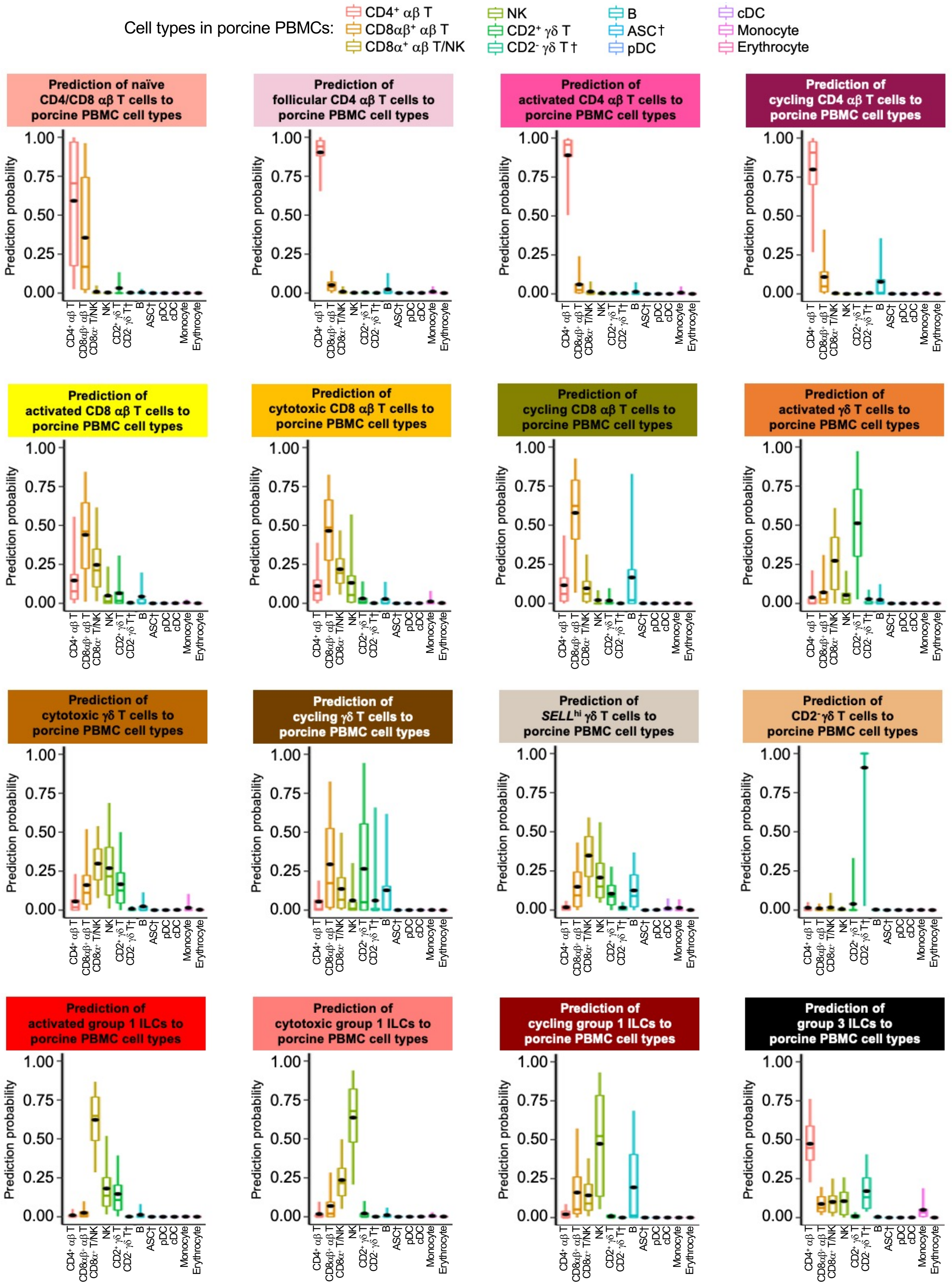

**Supplement to Figure 2-Figure 3. Prediction of porcine ileal T/ILC lineage lymphocytes to annotated cell types in porcine PBMCs.**

Box plots of the distribution of prediction probabilities (y-axes) for T/ILC lineage lymphocyte cell types from porcine ileum (represented by individual box plots) with labels transferred to annotated cell types of a porcine PBMC scRNA-seq reference dataset (x-axes and box plot color). Boxes span the interquartile range (IQR) of the data ($25^{th}$ and $75^{th}$ percentiles), with the median ($50^{th}$ percentile) indicated by a horizontal line. Whiskers span the $5^{th}$ and $95^{th}$ percentiles of the data. A red dot represents the data mean.

Query scRNA-seq data were derived from ileum of two seven-week-old pigs.

† Identical cell type annotations were given to cells in both porcine ileum and a reference scRNA-seq dataset. Cell type annotations were given to each dataset by independent rationales, and identical annotations do not necessarily indicate identical cell types were recovered from both porcine ileum and reference data.

Abbreviations: ASC (antibody-secreting cell); cDC (conventional dendritic cell); ILC (innate lymphoid cell); IQR (interquartile range); NK (natural killer); PBMC (peripheral blood mononuclear cell); pDC (plasmacytoid dendritic cell); scRNA-seq (single-cell RNA sequencing)

### Supplement to Figure 2-Figure 4

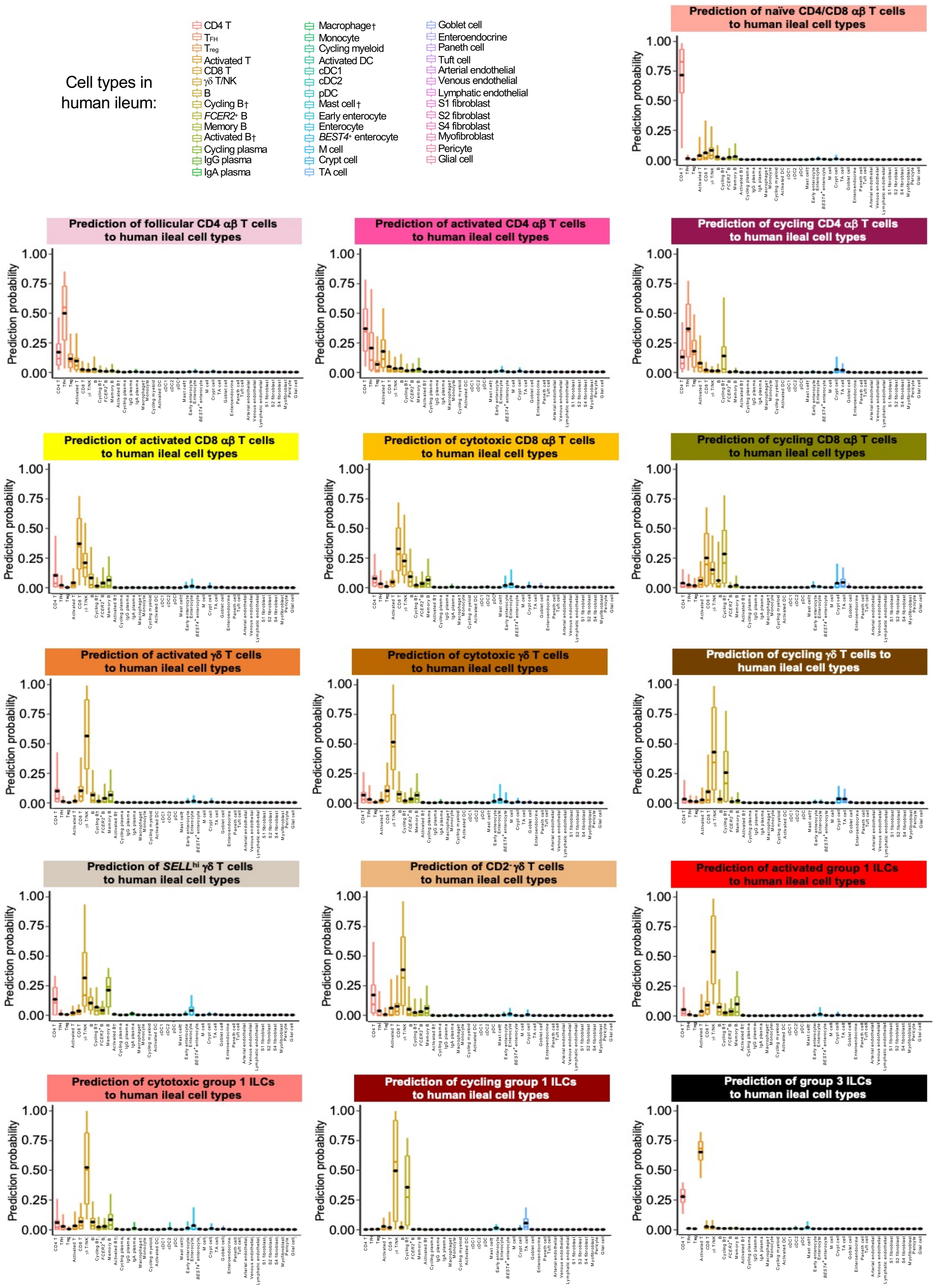

**Supplement to Figure 2-Figure 4. Prediction of porcine ileal T/ILC lineage lymphocytes to annotated cell types in human ileum.**

Box plots of the distribution of prediction probabilities (y-axes) for T/ILC lineage lymphocyte cell types from porcine ileum (represented by individual box plots) with labels transferred to annotated cell types of a human ileum scRNA-seq reference dataset (x-axes and box plot color). Boxes span the interquartile range (IQR) of the data (25$^{th}$ and 75$^{th}$ percentiles), with the median (50$^{th}$ percentile) indicated by a horizontal line. Whiskers span the 5$^{th}$ and 95$^{th}$ percentiles of the data. A red dot represents the data mean.

Query scRNA-seq data were derived from ileum of two seven-week-old pigs.

† Identical cell type annotations were given to cells in both porcine ileum and a reference scRNA-seq dataset. Cell type annotations were given to each dataset by independent rationales, and identical annotations do not necessarily indicate identical cell types were recovered from both porcine ileum and reference data.

Abbreviations: cDC (convnentional dendritic cell); DC (dendritic cell); ILC (innate lymphoid cell); IQR (interquartile range); pDC (plasmacytoid dendritic cell); NK (natural killer); scRNA-seq (single-cell RNA sequencing); TA (transit amplifying); $T_{FH}$ (T follicular helper); $T_{reg}$ (T regulatory)

### Supplement to Figure 2-Figure 5

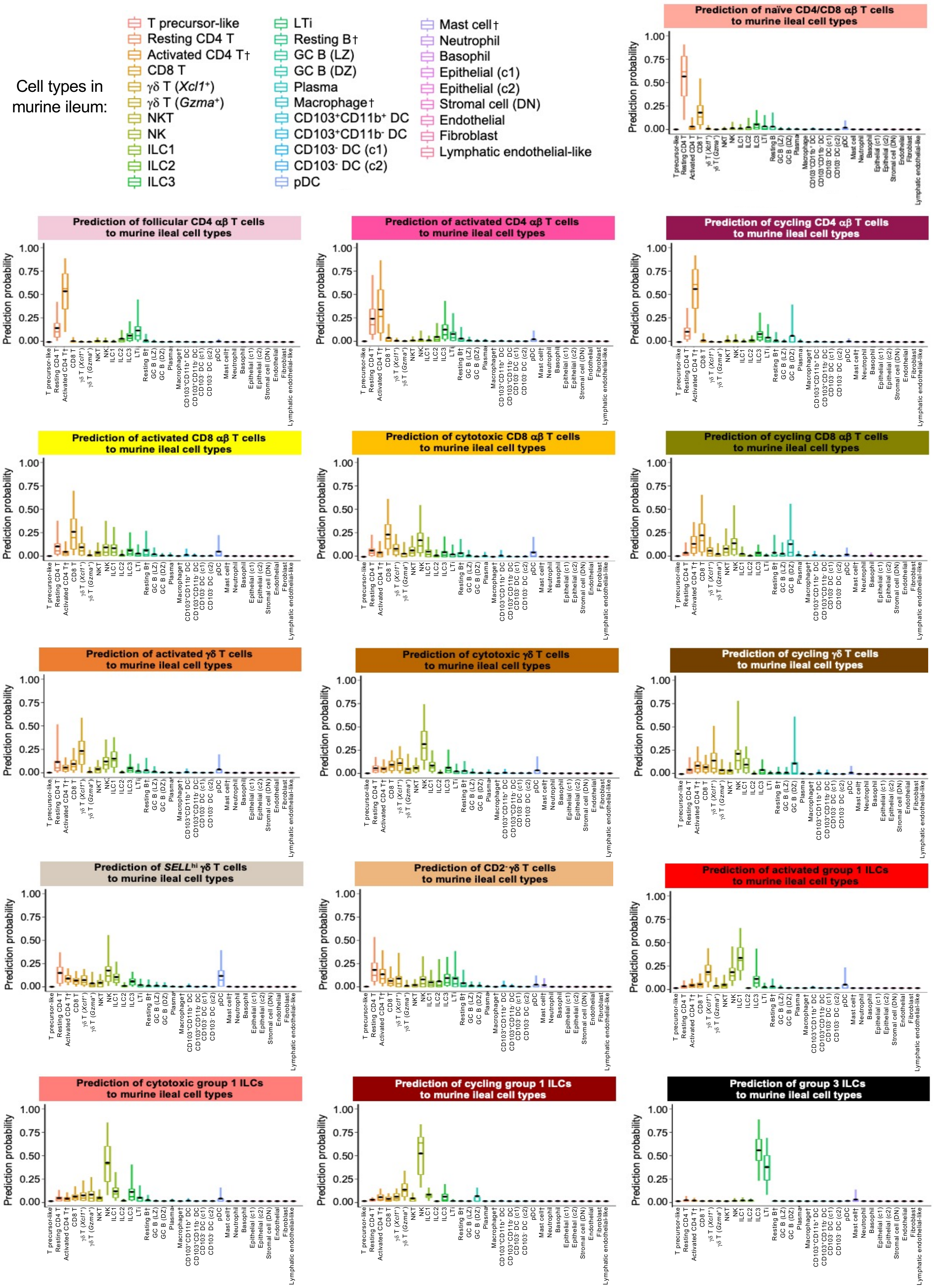

**Supplement to Figure 2-Figure 5. Prediction of porcine ileal T/ILC lineage lymphocytes to annotated cell types in murine ileum.**
Box plots of the distribution of prediction probabilities (y-axes) for T/ILC lineage lymphocyte cell types from porcine ileum (represented by individual box plots) with labels transferred to annotated cell types of a murine ileum scRNA-seq reference dataset (x-axes and box plot color). Boxes span the interquartile range (IQR) of the data ($25^{th}$ and $75^{th}$ percentiles), with the median ($50^{th}$ percentile) indicated by a horizontal line. Whiskers span the $5^{th}$ and $95^{th}$ percentiles of the data. A red dot represents the data mean.
Query scRNA-seq data were derived from ileum of two seven-week-old pigs.
† Identical cell type annotations were given to cells in both porcine ileum and a reference scRNA-seq dataset. Cell type annotations were given to each dataset by independent rationales, and identical annotations do not necessarily indicate identical cell types were recovered from both porcine ileum and reference data.
Abbreviations: c1 (cluster 1); c2 (cluster 2); DC (dendritic cell); DN (double-negative); DZ (dark zone); GC (germinal center); ILC (innate lymphoid cell); IQR (interquartile range); LTi (lymphoid tissue inducer); LZ (light zone); pDC (plasmacytoid dendritic cell); NK (natural killer); NKT (natural killer T); scRNA-seq (single-cell RNA sequencing); TA (transit amplifying); $T_{FH}$ (T follicular helper); $T_{reg}$ (T regulatory)

### Supplement to Figure 3-Figure 1

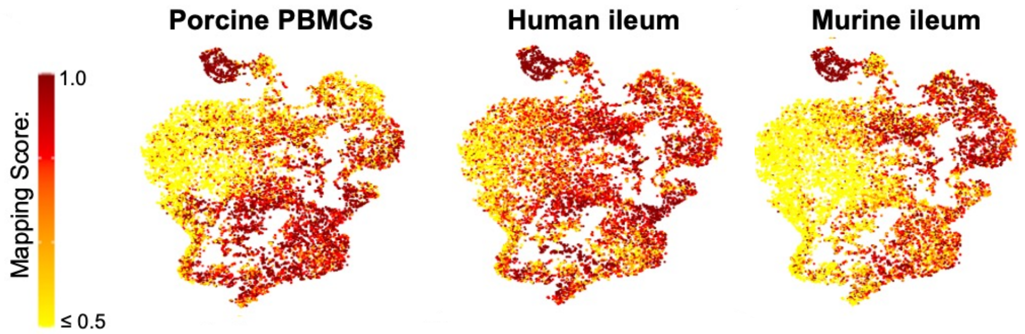

**Supplement to Figure 3-Figure 1. Overlay of mapping scores onto t-SNE reduction of B lineage lymphocytes from porcine ileum scRNA-seq data.**
Mapping scores from mapping of porcine ileum scRNA-seq query data to reference scRNA-seq datasets of porcine PBMCs (left), human ileum (center), and murine ileum (right). Mapping scores are the same as shown in **Supplement to Figure 1-Figure 11** but are now shown only for B lineage lymphocytes overlaid onto two-dimensional t-SNE visualization of porcine ileum scRNA-seq data shown in **Figure 3A**. Each point represents a single cell; the color of each point indicates mapping score to a corresponding reference dataset. Higher mapping scores indicate better representation of a cell from porcine ileum in a specified reference dataset.
Query scRNA-seq data were derived from ileum of two seven-week-old pigs.
Abbreviations: PBMC (peripheral blood mononuclear cell); scRNA-seq (single-cell RNA sequencing); t-SNE (t-distributed stochastic neighbor embedding)

#### Supplement to Figure 3-Figure 2

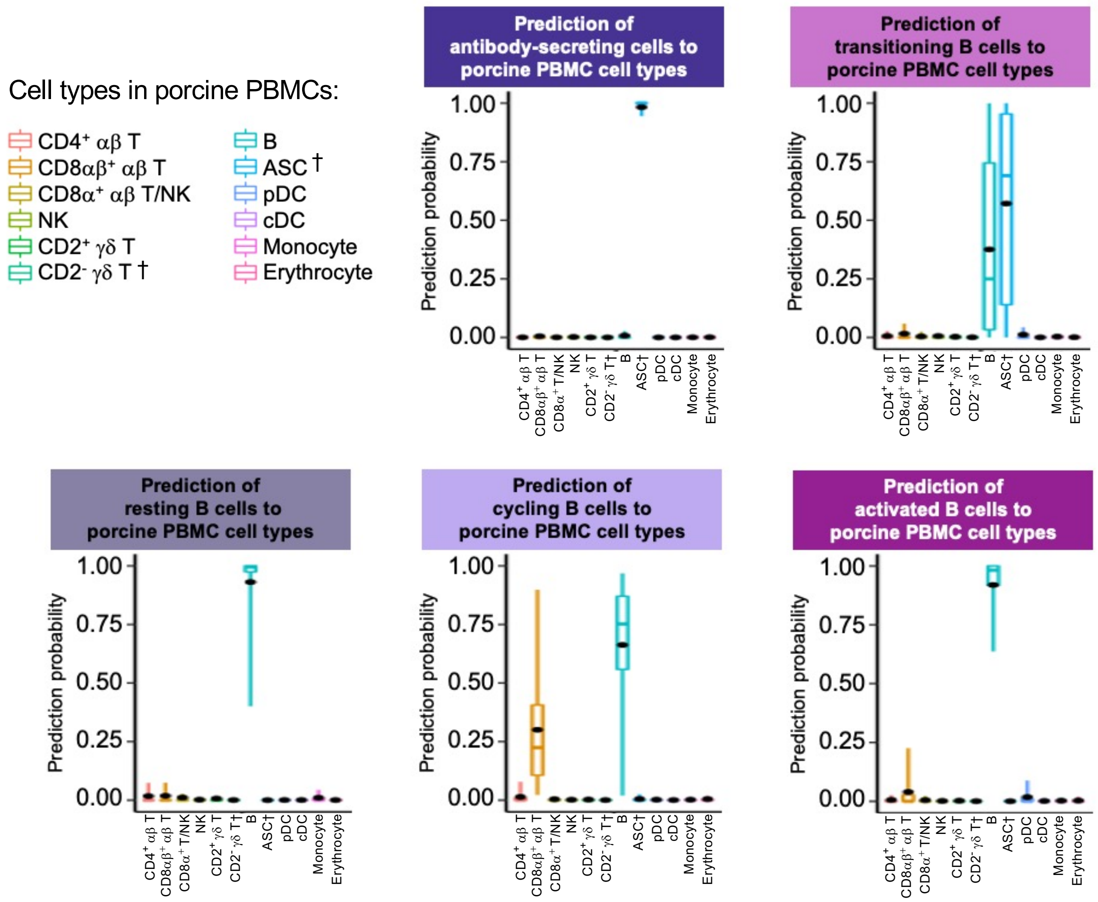

**Supplement to Figure 3-Figure 2. Prediction of porcine ileal B lineage lymphocytes to annotated cell types in porcine PBMCs.**

Box plots of the distribution of prediction probabilities (y-axes) for B lineage lymphocyte cell types from porcine ileum (represented by individual box plots) with labels transferred to annotated cell types of a porcine PBMC scRNA-seq reference dataset (x-axes and box plot color). Boxes span the interquartile range (IQR) of the data ($25^{th}$ and $75^{th}$ percentiles), with the median ($50^{th}$ percentile) indicated by a horizontal line. Whiskers span the $5^{th}$ and $95^{th}$ percentiles of the data. A red dot represents the data mean.

Query scRNA-seq data were derived from ileum of two seven-week-old pigs.

† Identical cell type annotations were given to cells in both porcine ileum and a reference scRNA-seq dataset. Cell type annotations were given to each dataset by independent rationales, and identical annotations do not necessarily indicate identical cell types were recovered from both porcine ileum and reference data.

Abbreviations: ASC (antibody-secreting cell); cDC (conventional dendritic cell); IQR (interquartile range); NK (natural killer); PBMC (peripheral blood mononuclear cell); pDC (plasmacytoid dendritic cell); scRNA-seq (single-cell RNA sequencing)

### Supplement to Figure 3-Figure 3

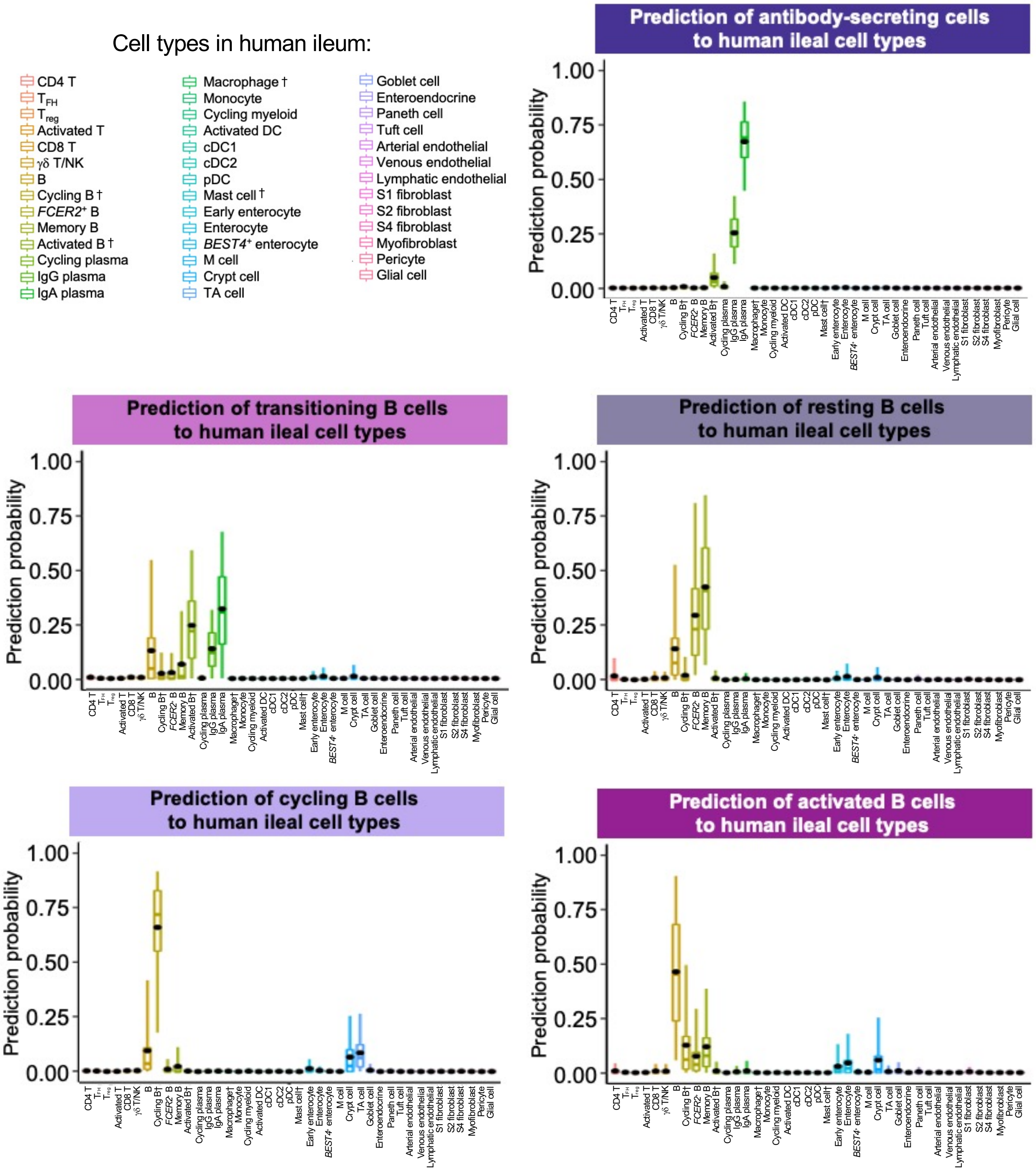

**Supplement to Figure 3-Figure 3. Prediction of porcine ileal B lineage lymphocytes to annotated cell types in human ileum.**

Box plots of the distribution of prediction probabilities (y-axes) for B lineage lymphocyte cell types from porcine ileum (represented by individual box plots) with labels transferred to annotated cell types of a human ileum scRNA-seq reference dataset (x-axes and box plot color). Boxes span the interquartile range (IQR) of the data ($25^{th}$ and $75^{th}$ percentiles), with the median ($50^{th}$ percentile) indicated by a horizontal line. Whiskers span the $5^{th}$ and $95^{th}$ percentiles of the data. A red dot represents the data mean.

Query scRNA-seq data were derived from ileum of two seven-week-old pigs.

† Identical cell type annotations were given to cells in both porcine ileum and a reference scRNA-seq dataset. Cell type annotations were given to each dataset by independent rationales, and identical annotations do not necessarily indicate identical cell types were recovered from both porcine ileum and reference data.

Abbreviations: cDC (convnentional dendritic cell); DC (dendritic cell); IQR (interquartile range); pDC (plasmacytoid dendritic cell); NK (natural killer); scRNA-seq (single-cell RNA sequencing); TA (transit amplifying); $T_{FH}$ (T follicular helper); $T_{reg}$ (T regulatory)

### Supplement to Figure 3-Figure 4

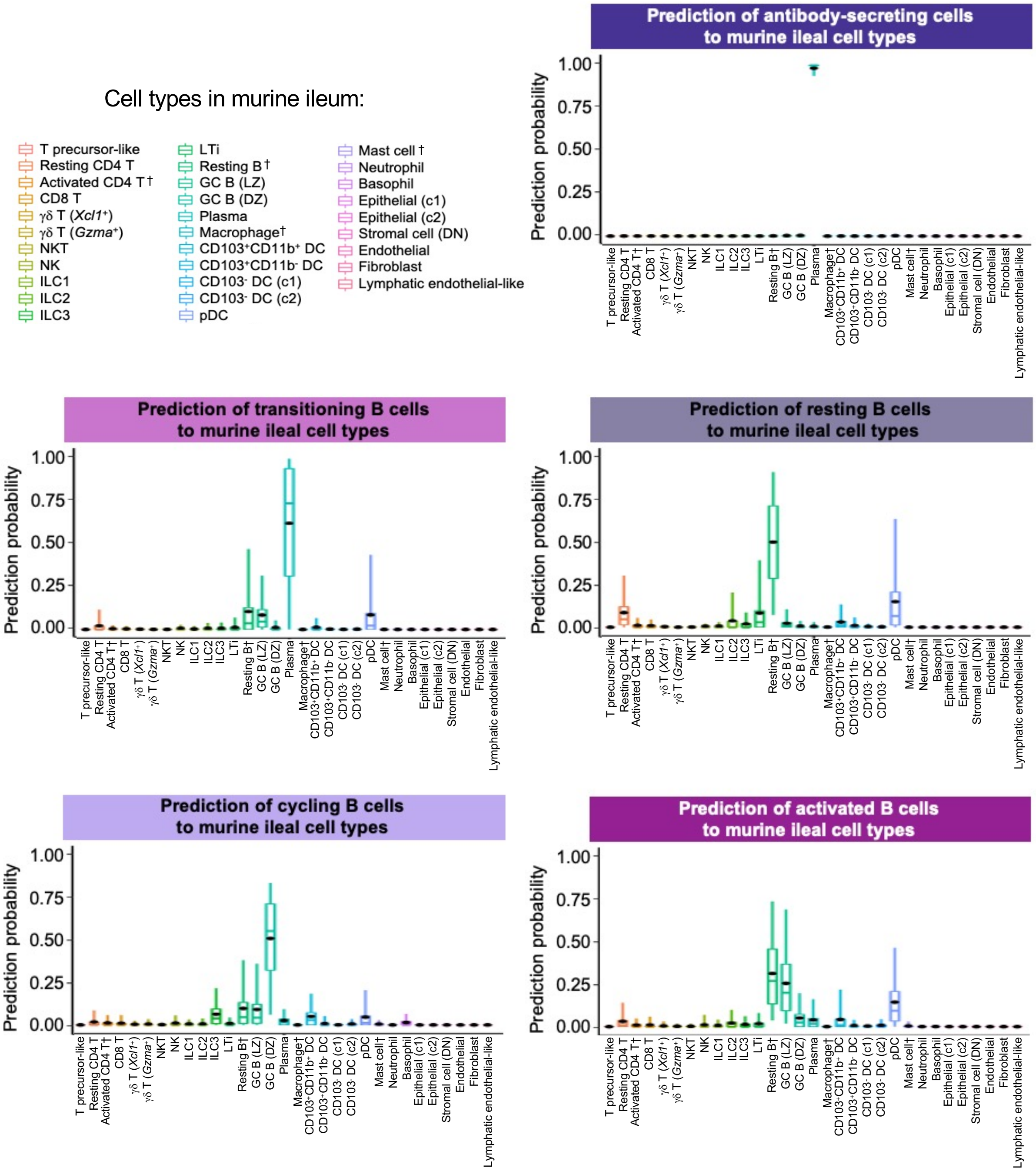

**Supplement to Figure 3-Figure 4. Prediction of porcine ileal B lineage lymphocytes to annotated cell types in murine ileum.**

Box plots of the distribution of prediction probabilities (y-axes) for B lineage lymphocyte cell types from porcine ileum (represented by individual box plots) with labels transferred to annotated cell types of a murine ileum scRNA-seq reference dataset (x-axes and box plot color). Boxes span the interquartile range (IQR) of the data (25$^{th}$ and 75$^{th}$ percentiles), with the median (50$^{th}$ percentile) indicated by a horizontal line. Whiskers span the 5$^{th}$ and 95$^{th}$ percentiles of the data. A red dot represents the data mean.

Query scRNA-seq data were derived from ileum of two seven-week-old pigs.

† Identical cell type annotations were given to cells in both porcine ileum and a reference scRNA-seq dataset. Cell type annotations were given to each dataset by independent rationales, and identical annotations do not necessarily indicate identical cell types were recovered from both porcine ileum and reference data.

Abbreviations: c1 (cluster 1); c2 (cluster 2); DC (dendritic cell); DN (double-negative); DZ (dark zone); GC (germinal center); ILC (innate lymphoid cell); IQR (interquartile range); LTi (lymphoid tissue inducer); LZ (light zone); pDC (plasmacytoid dendritic cell); NK (natural killer); NKT (natural killer T); scRNA-seq (single-cell RNA sequencing); TA (transit amplifying); $T_{FH}$ (T follicular helper); $T_{reg}$ (T regulatory)

### Supplement to Figure 4-Figure 1

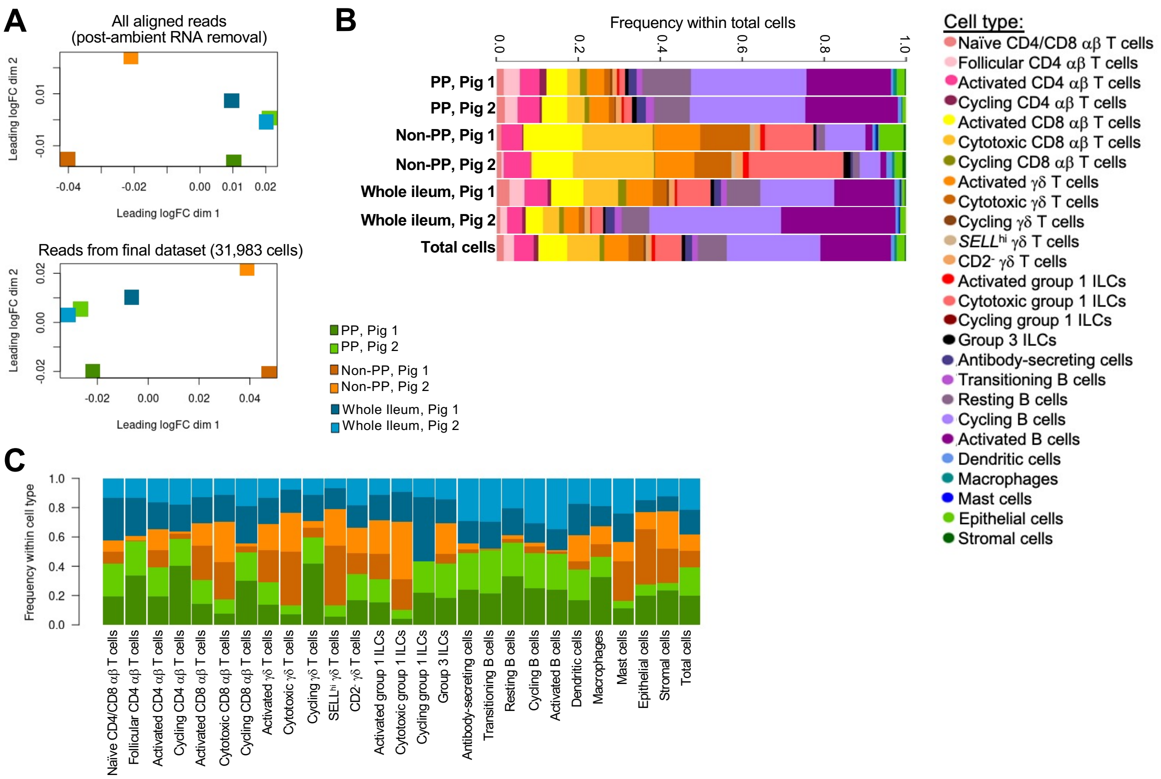

**Supplement to Figure 4-Figure 1. Comparison of sample types from scRNA-seq of porcine ileum.**

**(A)** Multidimensional scaling (MDS) plot of pseudobulk samples from six porcine ileal samples subjected to scRNA-seq. Pseudobulk samples are comprised of the cumulative gene counts from all reads/cells of each sample prior to quality control filtering (top) and in the final filtered dataset (bottom).

**(B)** Stacked bar plot of annotated cell type frequencies (x-axis) within each porcine ileal sample and total cells (y-axis) subjected to scRNA-seq. Bar size is indicative of total frequency (1) within each sample and is not indicative of the number of cells in each sample.

**(C)** Stacked bar plot of sample frequencies (y-axis) within each annotated porcine ileal cell type and total cells (x-axis) recovered via scRNA-seq. Bar size is indicative of total frequency (1) within each cell type and is not indicative of the number of cells in each cell type.

scRNA-seq data shown in **A-C** were derived from ileum of two seven-week-old pigs.

Abbreviations: dim (dimension); ILC (innate lymphoid cell); logFC (log fold-change); MDS (multidimensional scaling); PP (Peyer's patch); scRNA-seq (single-cell RNA sequencing)

### Supplement to Figure 4-Figure 2

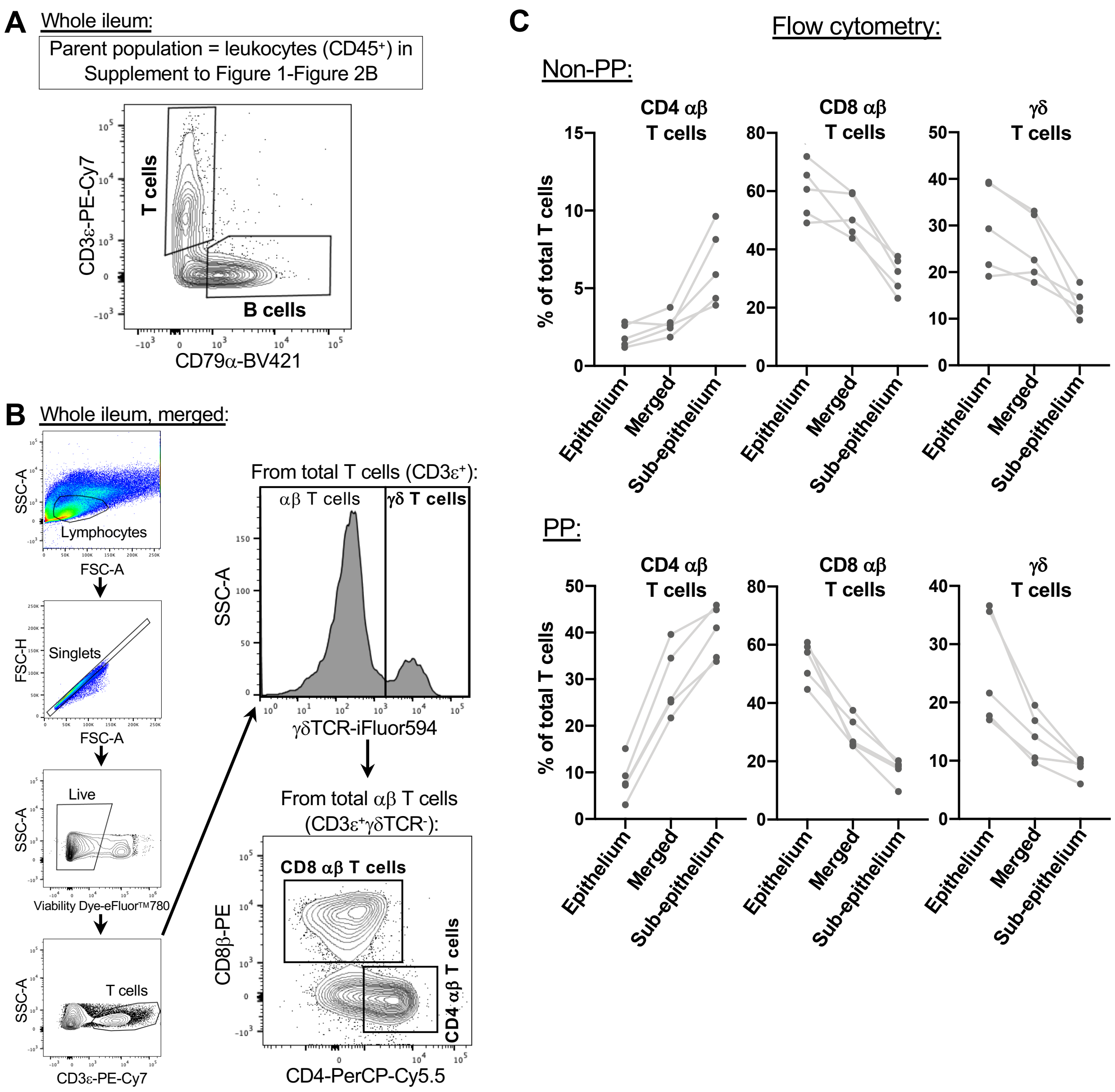

**Supplement to Figure 4-Figure 2. Validation of scRNA-seq lymphocyte compositions via flow cytometry.**

**(A)** Flow cytometry gating strategy used to identify percentages of T cells ($CD3\varepsilon^+$) and B cells ($CD79\alpha^+$) from total viable $CD45^+$ leukocytes within porcine ileal samples. Gating is shown for the same whole ileum sample (containing both regions with and without Peyer's patches) shown in **Supplement to Figure 1-Figure 2B**, starting from the parent population of cells captured and gated as leukocytes in **Supplement to Figure 1-Figure 2B**.

**(B)** Flow cytometry gating strategy used to identify percentages of CD4 $\alpha\beta$ T cells ($\gamma\delta TCR^-CD4^+$), CD8 $\alpha\beta$ T cells ($\gamma\delta TCR^-CD8\beta^+$), or $\gamma\delta$ T cells ($\gamma\delta TCR^+$) within total viable $CD3\varepsilon^+$ T cells of porcine ileal samples. Gating is shown for a whole ileum sample (containing both regions with and without Peyer's patches).

**(C)** Plot of the percentage of CD4 $\alpha\beta$ T cells ($\gamma\delta TCR^-CD4^+$; left), CD8 $\alpha\beta$ T cells ($\gamma\delta TCR^-CD8\beta^+$; center), or $\gamma\delta$ T cells ($\gamma\delta TCR^+$; right) within total T cells ($CD3\varepsilon^+$; y-axis) from PP (upper) and non-PP (lower) samples. Within each sample, cells were collected from epithelial, sub-epithelial, and merged (containing epithelial & sub-epithelial cell fractions; same as shown in **Figure 4H**) cell fractions (x-axes) and assessed by the flow cytometry gating strategy in **B**. Measurements from different cell fractions derived from the same animal are connected by a light grey line.

Flow cytometry experiments were not performed on animals used for scRNA-seq and were instead performed on four six-week-old pigs in **A** and five nine-week-old pigs in **B-C**.

Abbreviations: FSC-A (forward scatter area); FSC-H (forward scatter height); PP (Peyer's patch); scRNA-seq (single-cell RNA sequencing); SSC-A (side scatter area); TCR (T cell receptor)

### Supplement to Figure 4-Figure 3

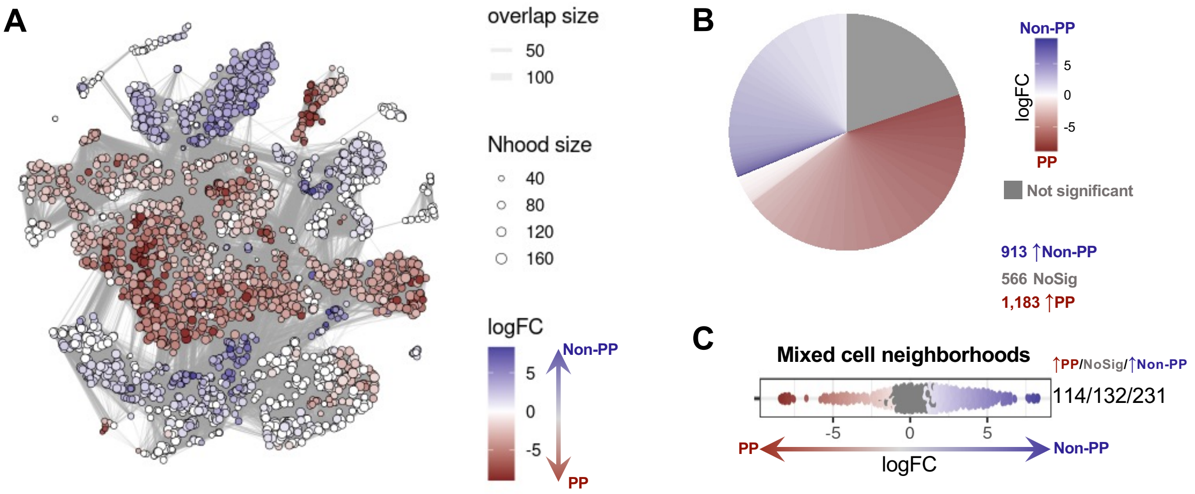

**Supplement to Figure 4-Figure 3. Differential abundance analysis of porcine ileum with versus without Peyer's patches.**

**(A)** Cell neighborhoods identified by differential abundance analysis. Only cells derived from PP and non-PP samples were included in differential abundance analysis but were overlaid back onto their original t-SNE coordinates of the full dataset that included whole ileum samples, shown in **Figure 1C-D**. Size of a circle indicates the number of cells in a neighborhood (Nhood size); color of a circle indicates magnitude of logFC in abundance in non-PP (blue) versus PP (red) samples; width of lines between cell neighborhoods indicates the number of overlapping cells found in each of two neighborhoods (overlap size).

**(B)** Pie chart of differential abundance results for cell neighborhoods shown in **A**. Grey indicates the proportion of cell neighborhood that were not differentially abundant, while cell neighborhoods with significantly increased abundance ($p<0.01$) in non-PP or PP samples are shown in blue and red, respectively. The logFC magnitude of differential abundance is also shown by red or blue shading.

**(C)** Plot similar to that shown for differential abundance analysis in **Figure 5J** but for all mixed cell neighborhoods that were not assigned as a specific cell type due to having $<70\%$ of cells belonging to a single cell type annotation. Each point represents an individual cell neighborhood. Grey points indicate cell neighborhoods that were not significantly more abundant in a specific sample type. Non-grey points indicate cell neighborhoods exhibiting differential abundance ($p<0.1$). Red/blue fill of differentially abundant points corresponds to the magnitude and direction of logFC. Red indicates increased abundance in PP samples, while blue indicates increased abundance in non-PP samples. On the far right, counts of cell neighborhoods with increased abundance in PP samples/no differential abundance/increased abundance in non-PP samples are shown for each cell type.

scRNA-seq data shown in **A-C** were derived from ileum of two seven-week-old pigs.

Abbreviations: logFC (log fold-change); Nhood (neighborhood); NoPP (Peyer's patch); No Sig (no significance); t-SNE (t-distributed stochastic neighbor embedding)

### Supplement to Figure 5-Figure 1

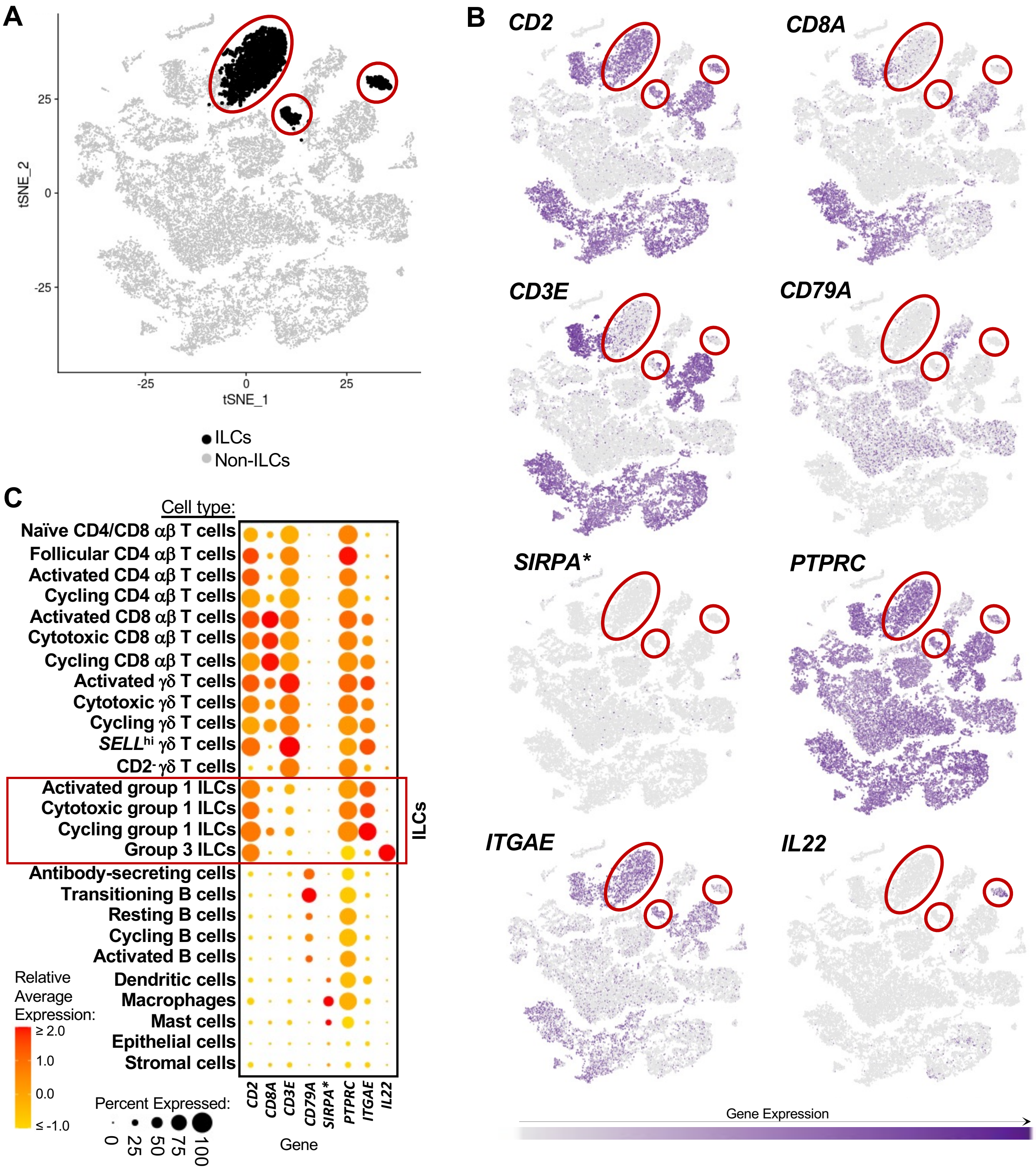

**Supplement to Figure 5-Figure 1. Gene expression profiles of group 1 and group 3 ILCs in porcine ileum.**

**(A)** Overlay onto two-dimensional t-SNE visualization shown in **Figure 1C-D** of cells annotated as ILCs (activated group 1 ILCs, cytotoxic group 1 ILCs, cycling group 1 ILCs, or group 3 ILCs; shown in black) in porcine ileum scRNA-seq data. All cells not annotated as ILCs (non-ILCs) are shown in light grey. Each point represents a single cell.

**(B)** Expression of a subset of canonical genes used to identify group 1 and group 3 ILCs in porcine ileum scRNA-seq data, overlaid onto two-dimensional t-SNE visualization coordinates of cells shown in **A**. Color of a point corresponds to expression level of a specified gene within a cell relative to all other cells in the dataset shown in **A**. Regions on the t-SNE plot with concentrations of ILCs are indicated by red circles.

**(C)** Gene expression patterns of canonical genes shown in **B** (x-axis) across annotated cell types in porcine ileum scRNA-seq data (x-axis). Within the plot, size of a dot corresponds to the percentage of cells expressing a gene within an annotated cell type; color of a dot corresponds to average expression level of a gene for those cells expressing it within an annotated cell type relative to all other cells in the dataset shown in **A**.

scRNA-seq data shown in **A-C** were derived from ileum of two seven-week-old pigs.

*Ensembl identifiers found in gene annotation were converted to gene symbols; refer to methods section '*Gene name modifications*' for more details

Abbreviations: ILC (innate lymphoid cell); scRNA-seq (singe-cell RNA sequencing); t-SNE (t-distributed stochastic neighbor embedding)

### Supplement to Figure 5-Figure 2

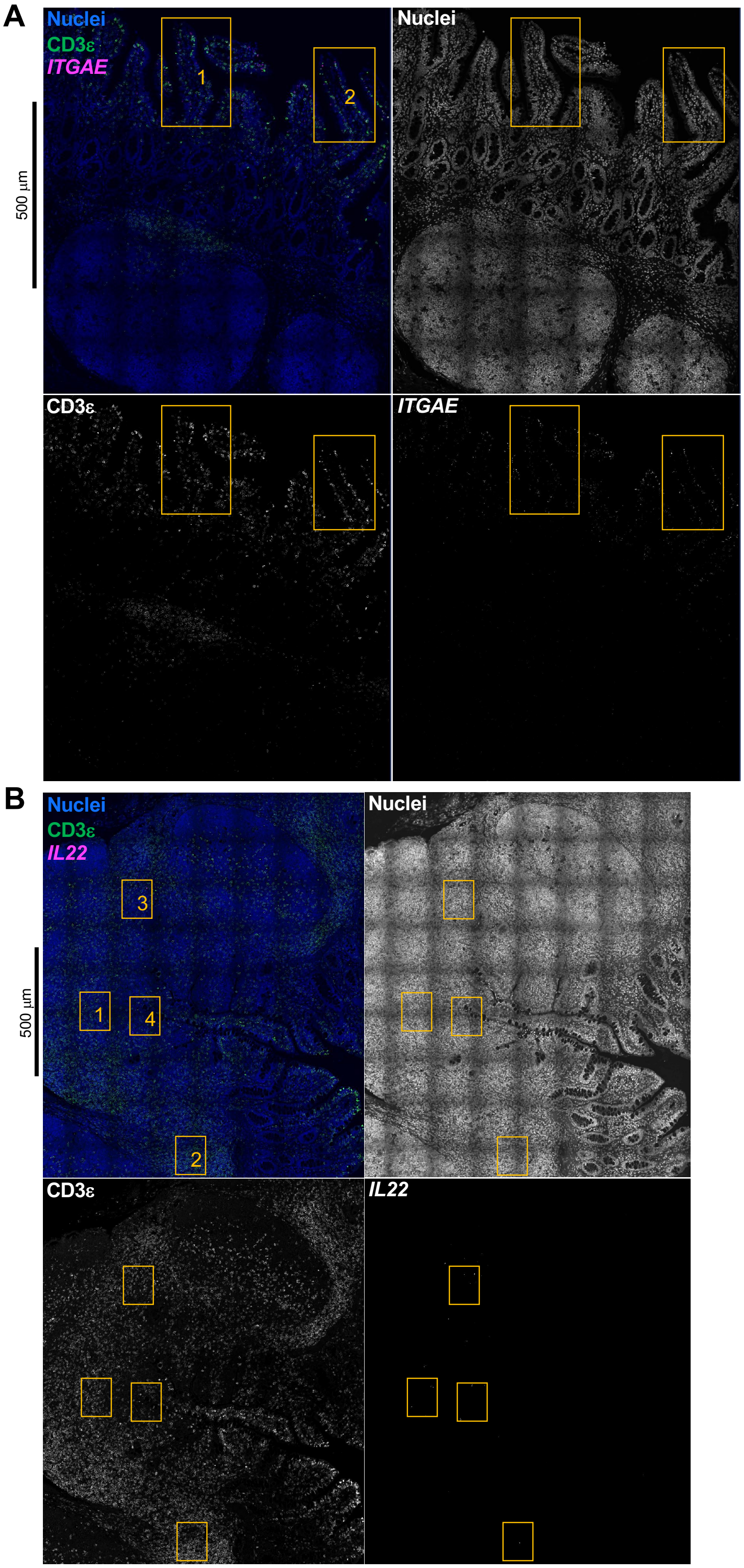

**Supplement to Figure 5-Figure 2. Microscopy images of *in situ* ILC detection.**
Confocal images for *in situ* detection of group 1 ILCs (**A**) and group 3 ILCs (**B**) in porcine ileum. Individual image frames were acquired at 60X magnification and stitched together. In both **A** and **B**, the upper left image shows overlay of all stains together, including nuclei (DAPI staining; blue), CD3ε protein (green), and RNA for *ITGAE* **(A)** or *IL22* **(B)** shown in magenta. The upper right image shows only nuclei staining (white); the lower left image shows only CD3ε protein staining (white); the lower right image shows only *ITGAE* **(A)** or *IL22* **(B)** RNA staining (white). All images in **A** and all images in **B** show the same captured frames. Yellow boxes indicate tissue areas shown at higher magnification in **Figure 5D-E**. Number of a box in the upper left panels corresponds to frame number shown in **Figure 5D-E**.
Dual IF/ISH experiments shown in **A-B** were conducted using a seven-week-old pig used for ileum scRNA-seq.
Abbreviations: IF (immunofluorescence); ILC (innate lymphoid cell); ISH (*in situ* hybridization)

### Supplement to Figure 6-Figure 1

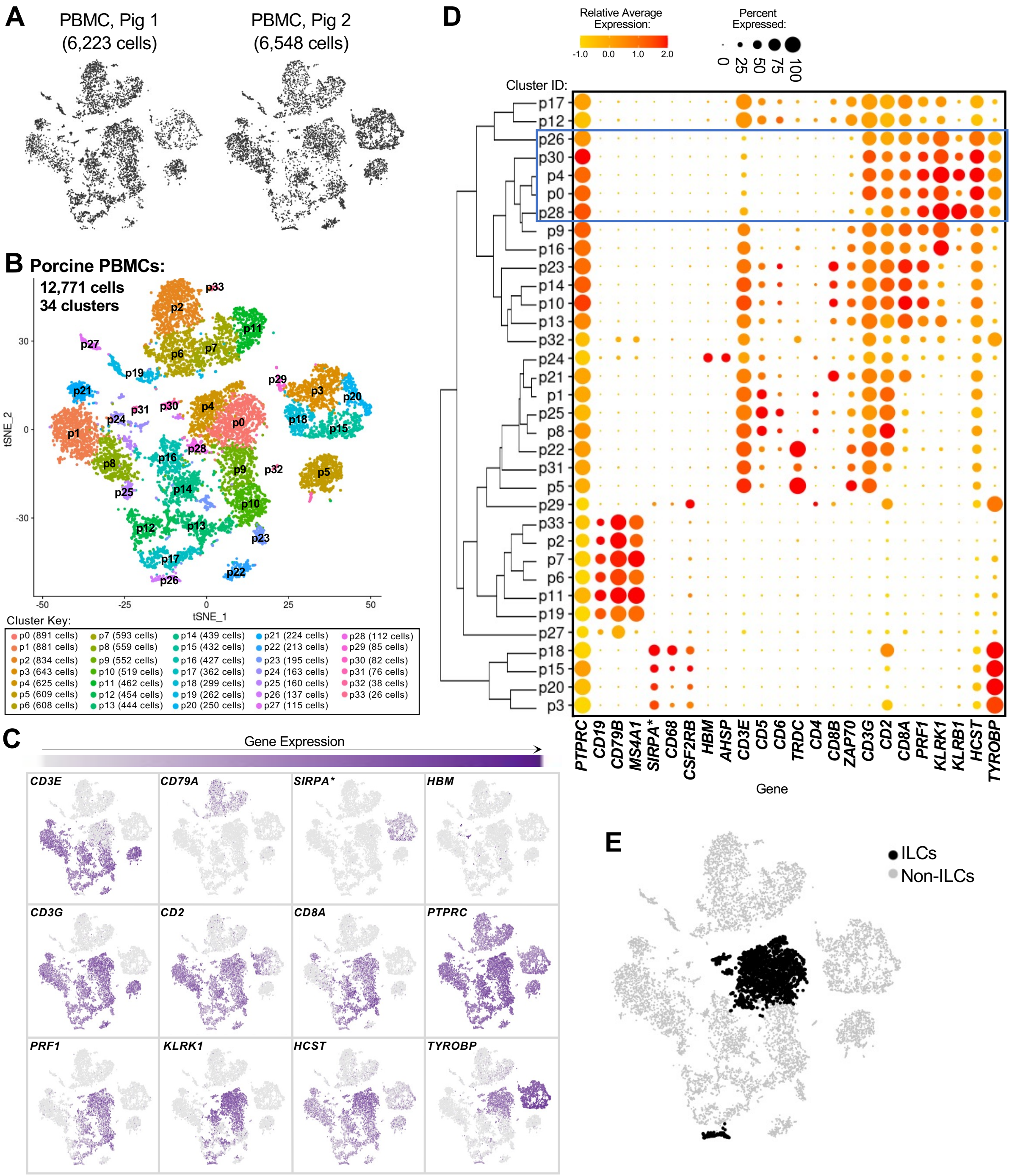

**Supplement to Figure 6-Figure 1. Identification of peripheral ILCs from porcine PBMC scRNA-seq data.**

**(A)** Two-dimensional t-SNE visualization of porcine PBMCs subjected to scRNA-seq and included in a final dataset following data processing and quality filtering. Each point represents a single cell. Plots show whether cells were derived from pig 1 (6,223 cells; left) or pig 2 (6,548 cells; right).

**(B)** Two-dimensional t-SNE visualization of 12,771 porcine PBMCs (combined cells from pig 1 and pig 2 shown in **A**). Each point represents a single cell; color of a point corresponds to one of 35 cell clusters a cell belongs to, with more transcriptionally similar cells belonging to the same cell cluster. The number of cells belonging to each cell cluster is listed in the cluster key.

**(C)** Expression of a subset of canonical genes used to identify ILCs overlaid onto two-dimensional t-SNE visualization coordinates of cells shown in **B**. Color of a point corresponds to expression level of a specified gene within a cell relative to all other cells in the dataset shown in **A**.

**(D)** Hierarchical relationship of cell clusters in porcine PBMCs shown in a dendrogram (left) and dot plot showing gene expression patterns within each cell cluster shown in **B** (right). In the dot plot, gene expression patterns of canonical genes used to identify ILCs (x-axis) across cell clusters shown in **B** are on the y-axis. Within the plot, size of a dot corresponds to the percentage of cells expressing a gene within a cell cluster; color of a dot corresponds to average expression level of a gene for those cells expressing it within a cell cluster relative to all other cells in the dataset shown in **B**. A blue box is drawn around clusters identified as ILCs.

**(E)** Overlay onto two-dimensional t-SNE visualization shown in **B** of cells annotated as ILCs (black; cell clusters p0, p4, p26, p28, p30) in **D**. All cells not annotated as ILCs (non-ILCs) are shown in light grey. Each point represents a single cell.

scRNA-seq data shown in **A-E** were derived from PBMCs of two seven-week-old pigs. Ileum and PBMC samples for scRNA-seq were collected from the same two pigs and processed in parallel.

*Ensembl identifiers found in gene annotation were converted to gene symbols; refer to methods section ‘*Gene name modifications*’ for more details

Abbreviations: ILC (innate lymphoid cell); PBMC (peripheral blood mononuclear cell); scRNA-seq (single-cell RNA sequencing); t-SNE (t-distributed stochastic neighbor embedding)

### Supplement to Figure 6-Figure 2

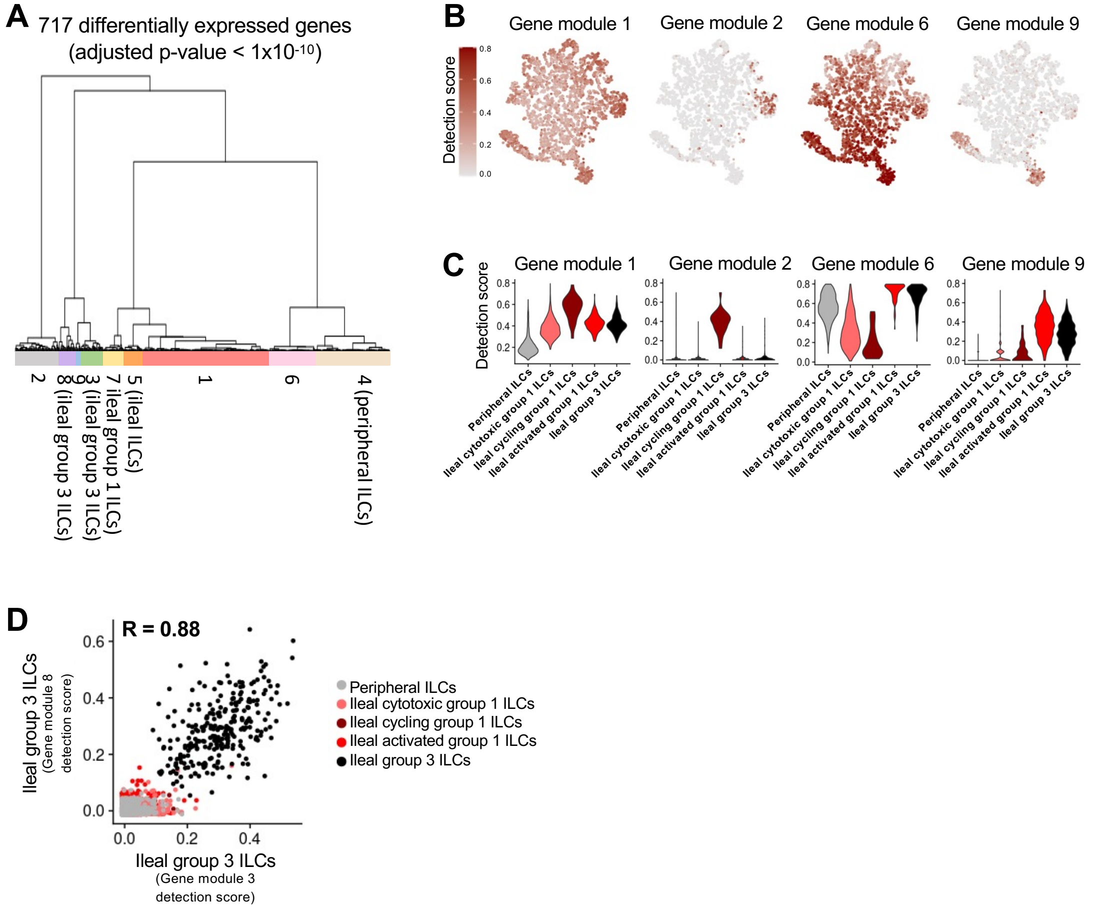

**Supplement to Figure 6-Figure 2. Gene module detection in ILCs from porcine ileum and PBMC scRNA-seq data.**

**(A)** Dendrogram of the top differentially expressed genes ($p < 1x10^{-10}$) recovered through multidimensional differential gene expression analysis. Based on the dendrogram, genes were grouped into nine gene modules, as indicated at the bottom of the dendrogram.

**(B)** Selected gene module detection scores from multidimensional differential gene expression analysis of cells shown in **Figure 6A** overlaid onto two-dimensional t-SNE visualization coordinates. Color of a point corresponds to detection score for a gene module within a cell.

**(C)** Violin plots summarizing gene module detections scores shown in **B** (y-axis) across annotated ILC types shown in **Figure 6A** (x-axis).

**(D)** Scatter plot of gene module 8 detection scores (y-axis) versus gene module 3 detection scores (x-axis) for all cells shown in **Figure 6A**. Each point represents a single cell; color of a point corresponds to cell type annotations shown in **Figure 6A**. Correlation value (R) is shown at the top of the plot.

scRNA-seq data shown in **A-D** were derived from ileum and PBMCs of two seven-week-old pigs. Ileum and PBMC samples for scRNA-seq were collected from the same two pigs and processed in parallel.

Abbreviations: ILC (innate lymphoid cell); PBMC (peripheral blood mononuclear cell); scRNA-seq (single-cell RNA sequencing); t-SNE (t-distributed stochastic neighbor embedding)
